# Inverse relationship between mitochondrial and non-coding nuclear DNA content in free-living kinetoplastids

**DOI:** 10.64898/2026.09.29.755400

**Authors:** Daryna Zavadska, Borys Olifirov, Julia A. Packer, Daria Tashyreva, Ryoma Kamikawa, Alastair G. B. Simpson, Daniel J. Richter

## Abstract

The evolution of genome size and eukaryotic genome organisation has puzzled evolutionary biologists for decades. Are differences in DNA content and protein-coding capacity of eukaryotic genomes a result of natural selection, or neutral processes? If they were shaped by natural selection, which features might be connected with these differences: cell volume, ecological strategy, availability of nutrients, etc.?

Kinetoplastids are protists with compact nuclear genomes that almost entirely lack introns, but whose mitochondrial DNA content (kDNA) can rival that of their nuclei.

We hypothesised that the reduction in non-protein coding nuclear DNA content during the evolution of kinetoplastids is connected with an increase in kDNA content: species with larger fractions of total cellular DNA that is allocated to kDNA will have concomitantly smaller fractions of non-coding DNA in the nuclear genome.

We developed a framework for estimation of DNA content in both the mitochondria and nuclei from quantitative fluorescence microscopy data, and inferred the fraction of non-coding DNA in nuclear genomes based on the combination of transcriptomic and fluorescence imaging data.

Across 10 genetically distant isolates from diverse free-living kinetoplastid lineages, there is indeed a trend towards a lower proportion of non-protein-coding nuclear DNA when kDNA proportion is higher relative to the total DNA in the cell. This observation is consistent with the hypothesis that in free-living kinetoplastids, mitochondrial and nuclear genomes might co-evolve in terms of their DNA content. This finding raises the possibility that natural selection acts on genome size evolution in kinetoplastids, and contributes evidence and methodology to address this question more broadly in eukaryotes.

## 1 Introduction

Kinetoplastids are a diverse group of free-living or parasitic unicellular flagellates that represent a monophyletic lineage of Euglenozoa, Discoba Adl et al. (2019); Gibson (2017). Kinetoplastids are a goldmine of bizzare biological characters: they posess glycosomes, modified peroxisomal organelles that compartmentalise much of the glycolytic pathway Hannaert and Michels (1994); Hannaert et al. (2003), a paraflegellar rod Maharana et al. (2015); Cachon et al. (1988), an unconventional kinetochore Akiyoshi and Gull (2014); Tromer et al. (2021), the non-canonical base J in nuclear DNA Borst and Sabatini (2008), and many other distinctive features of cell biology and metabolism (for example, Butenko et al. (2020); Vesteg et al. (2019); Donelson et al. (1999)).

The organization and function of kinetoplastid genomes is just as peculiar as their cellular structures and metabolic features. Kinetoplastid nuclear genomes have functionally unrelated genes arranged in polycistronic clusters, their nuclear mRNA undergoes trans-splicing via spliced leader addition, and transcriptional regulation of gene expression is largely absent (for example, Clayton (2016); Campbell et al. (2003)).

Notably, kinetoplastid genome sequences that are currently available have very few introns (as few as three per genome) Kostygov et al. (2024b). Intron poverty suggests that, as a lineage, kinetoplastids could have undergone “genome compaction” or “genome streamlining“: an increase in gene density and reduction of overall genome size, without the reduction of genome coding capacity Keeling (2004); Keeling and Slamovits (2005).

Genome compaction is not unique to kinetoplastids. Compact, intron-lacking genomes are also found in several other protist lineages Keeling (2004); Cavalier-Smith (2005); Derelle et al. (2006); López-Escardó et al. (2026); McGrath and Katz (2004). Compact genomes of protists (as well as converse cases of extremely expanded, giant genomes) are particularly interesting for studies addressing the evolution of genome organisation and cellular DNA content McGrath and Katz (2004). Many hypotheses and models that aim to explain evolution of total cell DNA content, protein-coding gene content, and its fraction relative to the total nuclear DNA have been proposed (see Discussion, Blommaert (2020); Lynch and Conery (2003); Rajic et al. (2005); Lynch (2006b); Elliott and Gregory (2015); Cavalier-Smith and Beaton (1999); Cavalier-Smith (1980, 2005); Lynch (2025)), with most of these explanations being subjects of ongoing debate.

It remains unclear whether DNA content and genome compaction has any meaningful fitness effect that can be subject to natural selection. Ecological, cellular, and molecular features of a given lineage could define fitness cost (if any) of a certain genome size and organisation. Nucleus/cell/body size, population-genetic characteristics, metabolic rate, trophic mode, division rate and ecological strategy have all been proposed to be connected with DNA content and genome organisation Hessen et al. (2010); Gregory (2001); Blommaert (2020); Keeling and Slamovits (2005); Cavalier-Smith and Beaton (1999); Cavalier-Smith (1980, 2005); Lynch (2025, 2006b); López-Escardó et al. (2026); Lynch and Marinov (2015); Jeffares et al. (2006).

Possibly the most remarkable lineage-specific feature of kinetoplastids is their so-called “kinetoplast DNA” (kDNA), which is in fact a mitochondrial genome. Kinetoplast DNA content is comparable to that of their nuclear DNA Lukeš et al. (2018). Since kDNA is such a significant part of the total DNA content of kinetoplastid cells it is worth considering when addressing the evolution of compact nuclear genomes and cellular DNA content in this lineage. In the kinetoplastids in which kDNA has been well characterised, thousands of small circular plasmids, known as “minicircles”, constitute the majority of kDNA, with the dozens of larger circular molecules, known as “maxicircles” making up a negligible fraction of kDNA content Lukeš, J., Lys Guilbride, D., Votypka, J., Zíková, A., Benne, R., and Englund, P. T. (2002); Liu et al. (2005); Hajduk and Ochsenreiter (2010). Maxicircles encode the primary mRNAs of mitochondrial rRNAs and proteins. However, mRNAs transcribed from maxicircles often need to be heavily edited (by uridine residue deletions/insertions) in order to become functional transcipts. This editing process is directed by the guide RNAs encoded on minircles Lukeš, J., Lys Guilbride, D., Votypka, J., Zíková, A., Benne, R., and Englund, P. T. (2002); Liu et al. (2005); Hong and Simpson (2003); Hajduk and Ochsenreiter (2010). The editing of primary mRNA is a stepwise process Hajduk and Ochsenreiter (2010); Simpson et al. (2000): if one of the minicircles/guide RNAs is missing, the upstream mRNA cannot be edited correctly, which (in the absence of correct guide RNA which allows re-editing) results in synthesis of altered mitochondrial proteins Wang et al. (2025). The evolutionary history and origin of such an excessively complex biological feature as kDNA is unclear Benne (1990); Simpson et al. (2000, 2002); Lukeš, J., Lys Guilbride, D., Votypka, J., Zíková, A., Benne, R., and Englund, P. T. (2002); Flegontov et al. (2011); Speijer (2006); mitochondrial genomes of many other eukaryotes encode dozens of proteins without the need for extensive mRNA editing and an accompanying enormous amount of DNA encoding for guide RNAs. Since the loss of minicircles and its encoded guide RNA(s) is highly likely to impair important biological functions Wang et al. (2025), loss of any kDNA fragment will likely face strong negative selection Lukeš, J., Lys Guilbride, D., Votypka, J., Zíková, A., Benne, R., and Englund, P. T. (2002); Covello and Gray (1993); cases of kDNA loss or reduction are known among parasitic trypanosomes, but these species are unable to perform aerobic respiration and are restricted to exist only in the anaerobic life stage Lai et al. (2008). So, throughout evolution, once RNA editing (and minicirles encoding for functional guide RNAs) appear, they are unlikely to be lost Covello and Gray (1993). The presence of exquisitely complex protein machinery for kDNA replication that ensures inheritance of a complete minicircle repertoire is plausibly an adaptation to prevent the loss of kDNA fragments Lukeš, J., Lys Guilbride, D., Votypka, J., Zíková, A., Benne, R., and Englund, P. T. (2002); Liu et al. (2005); Amodeo et al. (2023). kDNA may be a system of “irremediable complexity” evolving as a “ratchet”, in which complexity of the system is more likely to increase during the process of evolution Lukeš et al. (2011); Gray et al. (2010); Flegontov et al. (2011). In the case of kDNA, the immediate increase in complexity could possibly be connected with the increase in the number of guide RNAs, and therefore, kDNA content. Therefore, compacted nuclear genomes alongside the massive kDNA content in kinetoplastids present an interesting system for understanding genome size and DNA content evolution.

Based on the assumptions that (i) kDNA content is much more likely to increase in the process of evolution rather than decrease and (ii) nuclear genomes of kinetoplastids undergo (or have undergone) compaction by the loss of non-protein coding DNA, we hypothesise that the expansion of kDNA is connected with the compaction of the nuclear genome in free-living kinetoplastids. In this hypothesis, kDNA and nuclear genomes of free-living kinetoplastids co-evolve: the ratchet-like increase in kDNA content is compensated by the loss of non-protein-coding nuclear DNA, which leads towards compacted nuclear genomes. If this hypothesis cannot be rejected, it would also imply that the total DNA content per cell is shaped by natural selection in this particular protist lineage (see Discussion).

One way to test our hypothesis is to compare the fractions of total cellular DNA occupied by kDNA to nuclear protein-coding and nuclear non-coding DNA, across multiple isolates (“species”) of free-living kinetoplastids. If our hypothesis were true, we would expect isolates with a larger fraction of kDNA to have less non-protein-coding nuclear DNA and vice versa.

To test whether such an inverse relationship exists, we obtained data on kDNA and nuclear DNA content (including both coding and non-coding for the latter) and nuclear genome ploidy for 10 phylogenetically diverse free-living kinetoplastid isolates. For this, we developed a methodology for (i) quantitative fluorescent widefield imaging of the DNA-containing compartments of protist cells (modifying protocols from Wheeler et al. (2012); Lukeš et al. (2018)); (ii) semi-automatic discrimination of DNA-containing compartments in microscopy images and quantification of fluorescence intensity in separate compartments; (iii) inference of DNA content from fluorescence intensity using a robust statistical framework; (iv) verification of ploidy by combining transcriptomic sequencing data and Sanger sequencing of selected regions; (v) reliable estimation of the protein-coding nuclear DNA content from transcriptomic sequencing data. While (i)-(iii) allowed us to estimate the total nuclear and kDNA contents of non-axenic, free-living kinetoplastid isolates, (iv)-(v) permitted estimation of the protein-coding nuclear DNA content. Combining the fluorescence-based estimates of total DNA content and sequencing-based estimates of protein-coding DNA content, we calculated the fractions of kDNA and non-protein-coding nuclear DNA (estimated by substracting the lengths of predicted proteome from total nuclear DNA) relative to total cellular DNA for all 10 kinetoplastid isolates.

We found a negative relationship between the fractions of kDNA content and non-protein-coding nuclear DNA content, when these features are compared across the 10 isolates. This finding is consistent with the possibilty that expansion of kDNA is connected with the compaction of nuclear genomes in free-living kinetoplastids. This opens a discussion on on whether natural selection shapes the total cellular DNA content in kinetoplastids, with potential implications for the evolution of organelle DNA amount and structural complexity in eukaryotes.

## 2 Materials and Methods

### 2.1 Sampling and cultivation

### 2.2 Estimating length of protein-coding nuclear DNA

Transcriptome assemblies and predicted proteomes of the free-living kinetoplastid isolates were produced and described in Zavadska et al. (2025) and are publicly available via Figshare at DOI: 10.6084/m9.figshare.30214630. Briefly, raw reads were trimmed with fastp v0.20.1 Chen et al. (2018) and assembled using RNAspades v3.13.1 Bushmanova et al. (2019). The resulting assemblies were screened for contamination by identifying 18S/16S rRNA sequences belonging to contaminants. The taxonomic identity of contaminants was established based on 18S/16S rRNA, and a custom database of contaminant genomes corresponding to those 18S/16S rRNA sequences was produced. Contigs from assemblies matching to the custom database (via sequence homology) were removed. Cross-contamination among co-sequenced samples was eliminated using WinstonCleaner https://github.com/kolecko007/WinstonCleaner. Protein-coding sequences were predicted from decontaminated assemblies using TransDecoder v5.0.1 https://github.com/TransDecoder/TransDecoder.

Predicted proteomes were subjected to redundancy reduction using CD-hit v4.8.1 Li and Godzik (2006): proteins with more than 90% global amino acid sequence identity were reduced to a single representative in each proteome. Redundancy reduction at the proteome level did not result in the reduction of overall BUSCO Manni et al. (2021) completeness scores, but substantially reduced the number of duplicated BUSCO genes Figure S9. A set of Metakinetoplastina-specific Orthogroups (OGs) was inferred *de novo* from the proteomes of isolates used in this study and other proteomes of metakinetoplastid representatives using OrthoFinder v2.5.5 Emms and Kelly (2019) (see Figure S14, Section S8.8.4 for details). For the proteomes of strains used in this study, we identified short (*<*200 amino acid) proteins not assigned to any of the Metakinetoplastina-specific OGs, and short proteins assigned to OGs with sequences from only one species (“singletons”) Figure S15. Such proteins likely represent artefacts of assembly or protein prediction, since they have no homologs in related species and are short. The total length of these two groups of proteins was subtracted from the total proteome length in the final calculation of the protein-coding nuclear DNA content.

The total lengths of the redundancy-reduced proteomes (after removing artefactual sequences) was multiplied by 3 to obtain the total length of protein-coding nuclear DNA in the studied isolate to produce a final estimate of protein-coding nuclear DNA content.

To account for the possibility of obtaining incomplete assemblies/proteomes from transcriptome data, we first inspected the applicability of BUSCO datasets for measuring the completeness of assemblies/proteomes of free-living kinetoplastids using a set of controls Section S8.8.2, Figure S10. After discovering that BUSCO has limited applicability to our species, we developed a set of single-copy orthologous genes which can be used to infer more specific and accurate completeness metrics for free-living Metakinetoplastids Figure S14, Section S8.8.4. With this set, we verified that the proteomes used in this study do not require correction for completeness Section S8.8.4. Finally, we tested whether our pipeline of *de novo* transcriptome assembly, decontamination and protein prediction could have introduced systematic biases into our estimates of proteome completeness and total length. For this, we applied our pipeline to process publicly available transcriptomic reads of several protists for which well-annotated genomes were also available Section S8.8.3. We compared completeness and lengths of *de novo* transcriptome proteomes with the corresponding reference genome proteomes, and although we found large differences from the reference in some cases, this variation was not systematic Section S8.8.3.

### 2.3 Fluorescent staining and imaging

#### 2.3.1 Staining

All kinetoplastid cultures used for staining were grown in RS medium with the same concentration of Nutrient component https://mediadive.dsmz.de/medium/P5, diluted 1:100 in either the Non-Nutrient component https://mediadive.dsmz.de/medium/P4 or analogous oligotrophic media with fully defined composition, see Section S3.3. Kinetoplastid cultures were pre-adapted in a medium used for the experiment for at least two passages preceding the experimental one. Species with known genome sizes, used as controls, were grown in various suitable media, see Table S2.

Each culture used in staining was passed into fresh medium either 5-6, or 2-3 (depending on the species) days before the day of staining, to ensure, as much as possible, that (i) the cell population was in the stage of active growth, and cells were not starving and (ii) cell populations from different biological replicates were at approximately similar growth stages. The full record of passage dates, growth media, and other information on cultures used in all the biological replicates is available at https://github.com/beaplab/kineto_stats/blob/main/experimental_replicates_data.xlsx.

The staining protocol was adapted, with major modifications, from Wheeler et al. (2012). Cells were harvested by centrifugation at 13000 *g*, +17*^◦^*C, pellets were collected and pipetted onto 24×24 mm coverslips (thickness 0.13-0.16mm, COVN-024-200). The cells were fixed on the coverslip by the addition of an equal volume of 8%PFA (Sigma-Aldrich, F8775) in PBS, so that the final PFA concentration was 4%. Cells on coverslips were incubated in 4% PFA solution for 30 min at room temperature (RT); the fixed cells would passively sediment and attach to coverslips over this time. After 30 min of fixation, the liquid was removed from the coverslip, and the coverslip was fully immersed in 100% methanol (Sigma-Aldrich, 113351) which had been pre-cooled to −20*^◦^*C. A 4 h incubation in 100% methanol at −20*^◦^*C followed, to achieve membrane permeabilisation. Following the incubation in methanol, coverslips were immersed for 30min in large volumes (*>*5 ml/coverslip) of PBS at RT to wash methanol out of the sample. Coverslips were then incubated with 50 µg/ml RNAse A (Invitrogen PureLink, 12091-021) in PBS for 2 h at RT: this step was performed to digest unwanted cytoplasmic RNA, in order to prevent its staining with 4,6-diamidino-2-phenylindole (DAPI) in the following step. The RNAse A solution was removed from the coverslip, and a solution with 125 µg/ml propidium iodide (PI; Invitrogen, P1304MP) and 60 µg/ml DAPI (Sigma-Aldrich, MBD0015) in PBS was applied to the samples. Coverslips were incubated in this solution for 30min in darkness at RT, and afterwards transferred to glass slides, without any washing and without addition of mounting media. Therefore, the cells on the glass slides remained immersed in the staining solution from the moment of staining, during slide storage (at +4*^◦^*C), and throughout the process of imaging. This prevented possible biases that could have occurred due to the dissociation of either stain from the DNA over time. The concentrations of the stains in the solution exceeded the ones theoretically required to stain the DNA of the cells on the coverslip by orders of magnitude Section S4.4.

Following preparation, coverslip edges were sealed with nail polish, slides were stored between 6 and 20 hours at +4*^◦^*C, and were subsequently imaged within 48h after preparation.

#### 2.3.2 Imaging

Imaging was performed using an Axio Observer.Z1 widefield inverted microscope with a Plan-Apochromat 63x/1.4 Oil DIC M27 epifluorescence objective and Halogen X-Cite Xylis LED illumination system as a light source. The imaging setup was equipped with Hamamatsu HDCamC13440-20CU camera, and ZEN image acquisition software was used for the setup control.

Images were acquired from multiple fields of view (approximately 15 to 50) of each slide. Fields of view were distributed on the slide as evenly as possible. For each field of view, a z-stack of 8 slices, with a 1.2 µm interval between slices (see Section S5.5), with 1×1 binning, was acquired in three channels. All slices from one channel were imaged together, with channels in the following order: (1) DIC; (2) PI fluorescence (filter cube excitation 538-562nm, emission 570-640nm);(3) DAPI fluorescence (filter cube excitation 335-420nm, emission 383-470nm).The exposure time and the intensity of the excitation laser used for the PI fluorescence detection were constant throughout all the experimental series (DAPI intensity was not used for quantification, so constant exposure/intensity for this channel was not critical, but still maintained in the majority of imaging sessions - see https://github.com/beaplab/kineto_stats/blob/main/experimental_replicates_data.xlsx for details on acquisition settings).

#### 2.3.3 Principle of staining and selection of dyes

The DNA-binding dyes used for the staining, propidium iodide (PI) and DAPI, bind to different sites in the DNA molecule. PI is an intercalating stain, therefore, its binding to DNA is indifferent to sequence composition. These features make PI well-suited for the quantification of DNA content, and PI is widely used for this purpose, mainly, but not exclusively, in flow cytometry Noirot et al. (2002). DAPI is a minor-groove binding stain, and it binds preferentially to AT-rich sequences Larsen et al. (1989). The kDNA of kinetoplastids is known to be more AT-rich than the nuclear DNA, in all species in which it has been investigated Wheeler et al. (2012); Lukeš et al. (2018). The use of DAPI and PI in combination allows semi-automatic discrimination between the two DNA-containing compartments: kDNA and nuclear DNA, by estimating the ratios of fluorescence intensities of these two stains (see the section on the quantification of fluorescence intensity below).

### 2.4 Quantification of fluorescence intensity

The entire image analysis pipeline is implemented as a plugin for napari, an open-source multidimensional image viewer. It allows for flexible and robust semi-automatic analysis, with each step implemented as a separate widget within the plugin. Source code and comprehensive documentation are available via GitHub https://github.com/wisstock/kinetofluo-napari under an open-source licence.

For primary cell detection, Otsu thresholding was applied to the local gradient image of the DIC brightfield image. Brightfield image artefacts on the initial mask were filtered by comparison with fluorescent channels; only cell mask candidates that included fluorescence signals were kept. To avoid the influence of local background and fluorescent debris on the subsequent analysis steps, each candidate cell was segmented and analyzed separately within the cell mask only.

Nuclei of the control species were detected from the summed images of DAPI and PI spectral channels by simple Otsu thresholding and morphological filtering. To distinguish kDNA from nuclei in kinetoplastid isolates, we used a rudimentary blind unmixing technique. The compartments of interest should differ by the ratio of DAPI and PI signals, due to their differing AT-content. An AT-rich compartment should have higher absolute DAPI intensity and a higher DAPI/PI intensity ratio. To obtain an unmixed image of the AT-rich compartment (*I_ATrich_*) with enhanced contrast, the initial DAPI intensity images were multiplied by the DAPI/PI intensity ratio calculated on a pixel-by-pixel basis (see equation 1 below). The same logic was applied to obtain an unmixed image of the AT-poor compartment (*I_ATpoor_*); the initial PI intensity image was multiplied by the PI/DAPI ratio image (see equation 2 below). As the current segmentation implementation cannot handle overlapped compartments, overlapped areas were ignored in the final masks to prevent intensity overestimation.

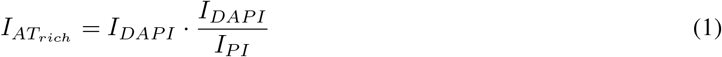

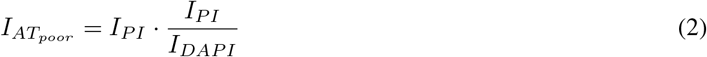

Species annotation was performed manually with the corresponding widget. In the process of manual annotation, we discarded cells in which kDNA, nuclear DNA, or cytoplasmic bacteria spatially overlapped. These annotations were saved to the final data frames alongside the summed PI intensity of the nucleus (AT-poor compartment) and kDNA (AT-rich compartment) for each individual cell. All tables with intensity measurements (used as an input for statistical analysis below) are stored on FigShare at DOI: 10.6084/m9.figshare.29088251.

### 2.5 Genome size prediction and statistical analysis

For the procedures described below, raw fluorescence intensities of nuclei/kDNA of all cells from all experimental series of each species were treated as one dataset (for a justification of this choice, see below, and Figure S5).

1. Fluorescence intensity values either below or above the 2nd or 3rd quartile *±* 1.5xIQR (interquartile range), were removed as outliers.
2. The cell cycle stage (“G1” or “G2”) was assigned to every nucleus/kDNA based on k-means clustering into two clusters. This was done to obtain a proxy for the copy number of genomes present in each cell: throughout the cell cycle, the amount of DNA in the genomes doubles, and cell cycles of the populations used in our experiments were not synchronised (see Section S6.6.2). After assigning the cell cycle stage, for each nucleus of the control species, the expected DNA content could be estimated based on the reference DNA content from genome sequencing data Table S2 and the cell cycle stage (two times the DNA content for the “G2” nuclei).
3. From a distribution of intensities within each of the two separate clusters of “G1” and “G2” compartments (nuclei and, if applicable, kDNA) intensity values either below or above the 2nd or 3rd quartile *±* 1.5xIQR (interquartile range), were removed as outliers.
4. A linear relationship between fluorescence intensities and DNA content was inferred from the intensity measurements and DNA content known for the control species. Quantile regression (using the “rq” function from the quantreg R package with default parameters) was used to model the DNA content’s dependence on the fluorescence intensity. Based on the regression model, the DNA content of nuclei and kDNA could be predicted for each kinetoplastid cell.
5. Finally, the median values of nuclear and kDNA content were estimated from the distributions of predicted DNA content for each kinetoplastid isolate. Only the cells assigned to the “G1” cell cycle stage were considered. The confidence intervals of the medians were estimated using the bootstrap bias-corrected, accelerated (BCa) method with 5000 bootstrap replicates (“groupwiseMedian” from the rcompanion R package with default parameters).

For performing the statistical analysis described above, we assessed different options for subsetting the dataset for both clustering (combined with outlier filtering) and regression inference: these procedures could be performed (1) on the intensity measurements obtained from all the experiments (biological replicates) combined; (2) separately on a subset of measurements from each experiment (biological replicates); or even (3) separately on a subset of measurements from each slide imaged during each experiment (technical replicates).

We evaluated the performance of each subsetting option by treating one out of the four control species as unknown and then predicting its DNA content from fluorescence intensities, by using the other three control species to build the corresponding regression. These predicted DNA contents were compared against those known from genome sequencing data. A detailed explanation and justification of the final approach used for DNA content prediction (in which all cells from all experimental series for each species were treated as a single dataset) is provided in Section S6.6.

The performance of the k-means clustering procedure used for cell cycle stage assignment was additionally assessed by running a series of stainings where the cell cycle propagation of one control species was blocked by hydroxyurea, as described in Section S6.6.2. Based on this experimental validation, we concluded that clustering performs best when fluorescence intensity data from all experiments and all technical replicates is analysed together as a single distribution, corroborating the results of the leave-one-out experiment on control species Figure S6, Figure S7.

## 3 Results

### 3.1 Validating the method

We ran a series of validations to assess the accuracy of DNA content estimation by measuring control species with known genome sizes. In each validation, one of the control species with known nuclear DNA content was treated as “test” isolate: its nuclear DNA content was estimated *de novo* based only on the fluorescence intensities that were experimentally measured for this species. The other three control species were used for linear model inference in such validations. The distributions of predicted DNA content were then compared with the known genome sizes of each species Figure 1. We found that the distributions of estimated DNA content had peaks equal to or close to the values expected according to genome sequencing. In order to produce point estimates of DNA content, distributions of the predicted DNA content values were further summarised as medians, as explained in Methods, and in Section S6.6. Medians of the estimated “G1” DNA content deviated from the expected values by a maximum of 10% of the expected DNA content Figure 1.

**Figure 1:**
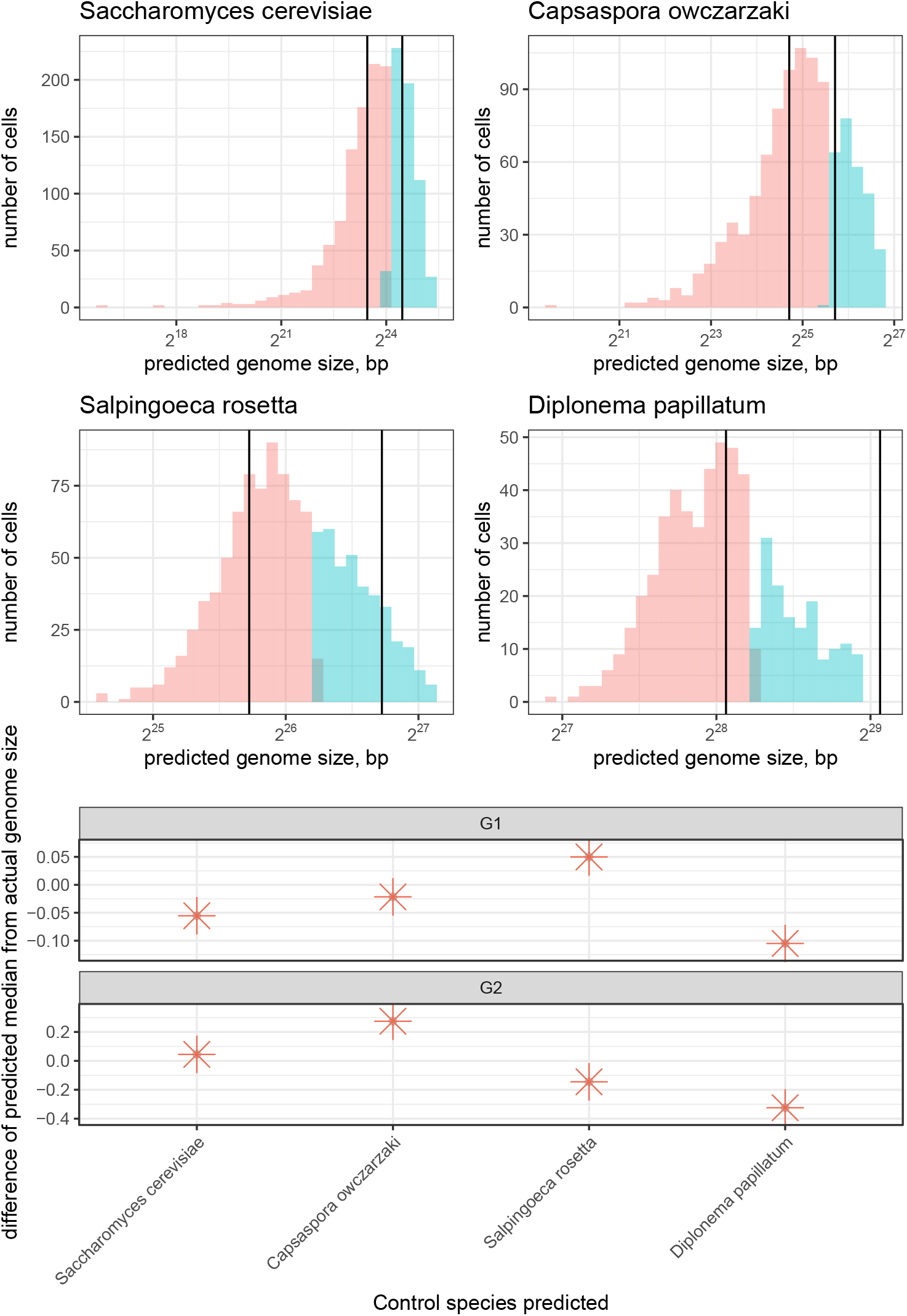
Top: Distribution of DNA content predicted for control species from fluorescence intensities, compared to their sequenced genome sizes. The vertical axis represents the number of G1 (red) and G2 (blue) cells with a given predicted DNA content (horizontal axis). Solid vertical lines indicate the known sizes of control species’ genomes for both G1 and G2 cells. The distributions tend to center around the vertical lines, indicating that the predicted DNA content generally is close to the sequenced genome size for control species. **Bottom: Difference between median estimated DNA content and expected value.** Each point represents a median obtained from the distribution of predicted DNA content, from which the sequenced genome size was subtracted, and the difference normalised by the sequenced genome size: (*Median_predicted_ − bp_actual_*)*/bp_actual_*.

**Figure 2:**
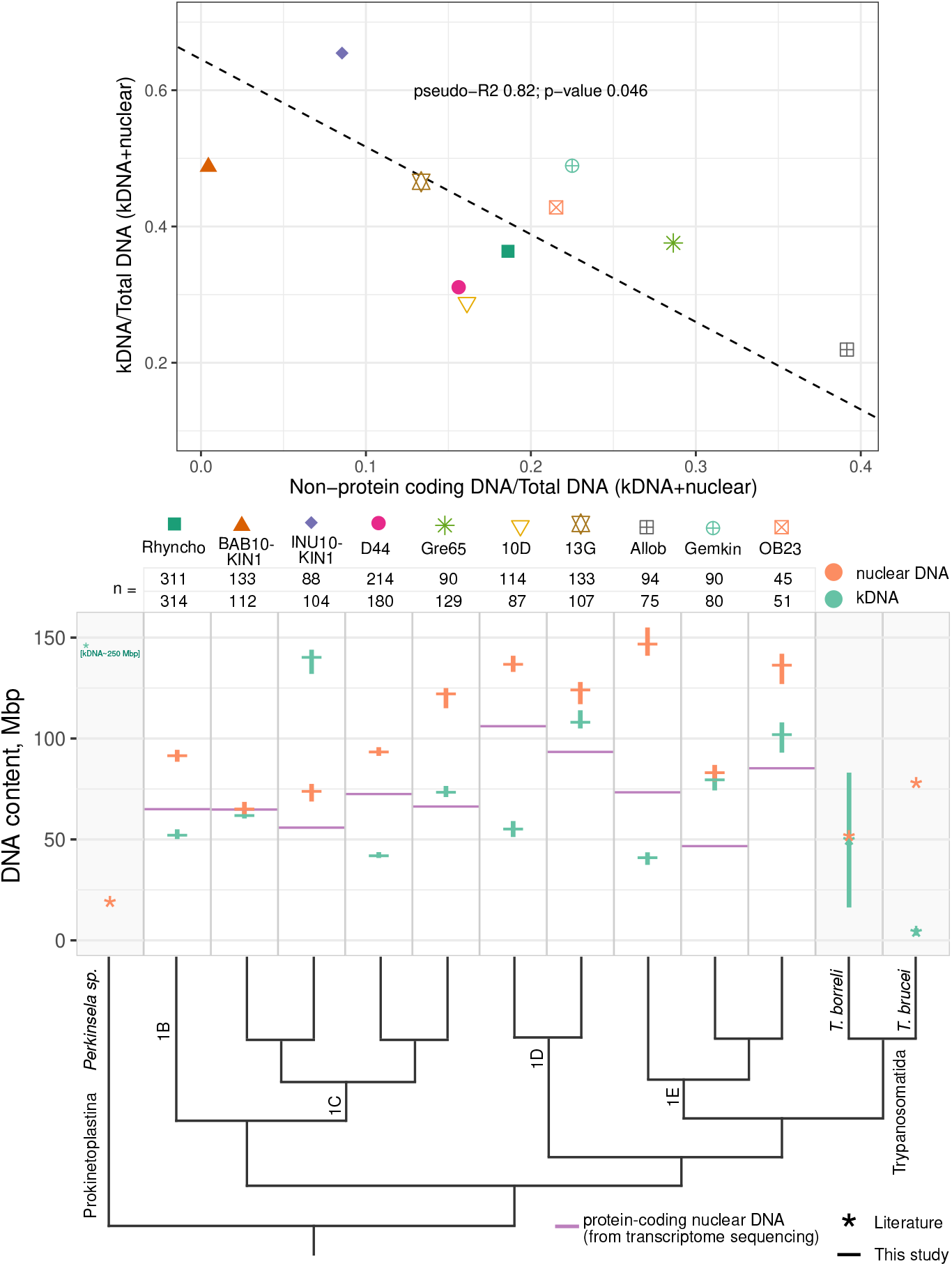
Top: Non-protein-coding nuclear and kDNA content relative to the total cellular DNA content of each isolate. The vertical axis represents the fraction of cellular DNA constituted by the kDNA; the estimates are obtained from the medians of predicted DNA content of G1 cells, for each kinetoplastid isolate: 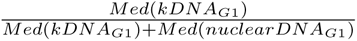. The horizontal axis represents the fraction of cellular DNA constituted by non-protein-coding nuclear DNA: 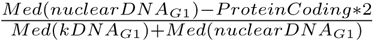. The size of the non-protein-coding part is calculated by subtracting the total length of the non-redundant predicted proteome, multiplied by 2 (as our kinetoplastid isolates are diploid), from the total nuclear DNA content predicted from fluorescence intensity. The grey line is a linear fit obtained from orthogonal distance regression (see text). **Bottom: Nuclear, protein-coding nuclear and kDNA content of kinetoplastid isolates.** Horizontal lines represent a median of predicted nuclear (red) and kDNA (blue) content of G1 cells. Each column contains estimates for a given kinetoplastid isolate, along with the three other species for which the estimates are available from the literature (grey background, asterisks) Lukeš et al. (2018). Vertical whiskers represent confidence intervals of the median, calculated by bootstraping (see Methods) for the isolates from this study; whiskers of estimates from the literature indicate the standard errors reported in Lukeš et al. (2018). Note that the standard error was not depicted for the kDNA of *Perkinsela sp.*, as it is 100Mbp and would therefore extend above the limit of the vertical axis Lukeš et al. (2018). The size of the non-redundant protein-coding part for **G1, diploid** cells, measured from transcriptome sequencing, is indicated by the wide purple line. The counts above the plot show the total number of G1 nuclei (top row) and kDNA (bottom row) of a given isolate used in median estimation (counts of nuclei and kDNA from a same isolate differ due to outlier filtering). The cladogram represents the evolutionary relatedness of the species, according to a phylogenomics study based on 444 single-copy orthologous genes Zavadska et al. (2025). Nodes of the major metakinetopla_1_st_0_id lineages are labelled: our isolates belong to neobodonid clades 1B, 1C, 1D and 1E.

### 3.2 Genome sizes of free-living metakinetoplastids

We estimated kDNA and nuclear DNA contents for 10 isolates of the free-living kinetoplastids. In parallel, we estimated the size of the protein-coding portion of their nuclear genomes from the total lengths of their redundancy-reduced proteomes. We ensured that redundancy reduction with CD-hit prevents over-estimation of the total length of protein-coding part without affecting completeness Figure S9. Benchmarking BUSCO metrics Manni et al. (2021) has shown that neither Euglenozoa nor Eukaryota BUSCO(v.10) gene sets are good indicators of completeness for euglenozoans other than trypanosomatids Section S8.8.4. So we additionally verified completeness of our proteomes using a custom set of Metakinetoplastid single-copy orthologs (M3USCO), which has shown that all proteomes used in our study are *>* 90% complete, with exempt of Gemkin isolate, which is *>* 85% complete Figure S14. We also verified that all 10 kinetoplastid isolates used in this study are diploid Section S7.7.

Fluorescence-based estimates of the total nuclear DNA content are concordant with the estimates of the length of the protein-coding nuclear DNA; none of the nuclear DNA content estimates implied a genome size smaller than the estimated protein-coding content Section S7.7. In addition, the confidence intervals of our estimates are small relative to the numerical value of each estimate, and do not exceed 10%.

We next interpreted these robust estimates for the sizes of the genomes and their protein-coding proportions for representatives of multiple lineages of kinetoplastids in the context of kinetoplastid evolution. The cladogram in Section S7.7 represents the phylogenetic relationships among the 10 isolates from this study (based on phylogenomics of 444 single-copy *de novo* inferred orthologous genes Zavadska et al. (2025)), and 3 more species for which DNA content estimates were previously available in the literature. Some general observations on the DNA contents across free-living kinetoplastids:

1. Estimated total kDNA content (of G1 cells) ranges from 41 to 144 Mbp.
2. Estimated total nuclear DNA content (of **haploid, G1** cells) ranges from 37 to 74 Mbp.
3. The total length of the protein-coding portion of the nuclear genome (of **haploid** G1 cells, based on the redundancy-reduced predicted proteome) ranges from 23 to 53 Mbp.
4. Neither the DNA contents (both kDNA and nuclear) nor length of the protein-coding part are conserved within major free-living kinetoplastid clades;
5. Kinetoplast DNA content is larger than that of the nucleus in 1 isolate out of 10 examined; 2 more isolates have almost equal kDNA and nuclear DNA contents, while the rest have nuclear DNA content higher than that of kDNA.

Finally, for each isolate, we estimated the fraction of the total cell DNA that is represented by kDNA and non-protein-coding nuclear DNA. The overall trend is that the larger the fraction of total cellular DNA that is occupied by kDNA, the smaller is the fraction represented by non-protein-coding nuclear DNA Section S7.7. Although the number of isolates (species) is relatively limited, there is a negative correspondence between the two variables: orthogonal distance regression slope of −1.29, pseudo-R^2^ of 0.82, p-value of 0.046 based on 5000 bootstrap replicates (“odregress” function from pracma R package, default parameters).

## 4 Discussion

### 4.1 Possible drivers of DNA content (co)evolution in free-living kinetoplastids

Our primary interest in this study is to test whether there is a relationship between kDNA content and non-protein coding nuclear DNA content. This could imply that kDNA and nuclear genome size have experienced adaptive evolution, either together or independently.

The inverse correspondence between the kDNA and non-coding nuclear DNA content that we observe Figure 1 is consistent with the possibility of nuclear and kDNA content co-evolution: the larger is the kDNA, the smaller is the non-coding portion of the nuclear genome in a given species. This indicates the total DNA content per cell could have been restricted to some limit during kinetoplastid evolution.

What are the factors that could influence or constrain the total amount of DNA in kinetoplastid cells? Multiple evolutionary forces, cellular/genomic features and environmental parameters have been proposed as the determining factors influencing genome size evolution and, specifically, genome compaction (see Introduction and Hessen et al. (2010); Gregory (2001); Blommaert (2020); Keeling and Slamovits (2005); Cavalier-Smith and Beaton (1999); Cavalier-Smith (1980, 2005); Lynch (2025, 2006b); López-Escardó et al. (2026); Lynch and Marinov (2015); Jeffares et al. (2006)). Blommaert (2020) provides a thorough overview and classification of this varied assemblage of hypotheses and models. These hypotheses assume different impacts of natural selection and neutral processes on the evolutionary changes in DNA content Blommaert (2020). Below we briefly outline a few hypotheses, existing experimental evidence to support or contradict them, and discuss how they may apply to the free-living metakinetoplastids.

#### 1. Resource and energy limitation

For prokaryotes, the fraction of a cell’s energy invested in the replication was found to be sufficiently large to be the target of natural selection; for a single unicellular eukaryote considered, (*S. cerevisiae*), this was a borderline case Lynch and Marinov (2015).

It has been claimed that energetic costs for genome maintenance constitute a significant fraction of the cell’s energy for prokaryotes, but are negligible for eukaryotic cells Lane and Martin (2010), this supposition was criticised, however (for example, Lynch (2025); Schavemaker and Muñoz-Gómez (2022)). The relative fraction of cellular energy invested in genome maintenance can vary across protist lineages, as each cellular trait has a certain energetic cost Schavemaker and Lynch (2025), and protists from different lineages can possess different unique (and possibly energetically costly) traits. kDNA is a unique trait of kinetoplastids. It is not unreasonable to suppose that genome sizes of the free-living kinetoplastids could be constrained by the amount of energy required to maintain them. To find out the extent of such a constraint, measurements like the ones on yeast Lynch and Marinov (2015), should be performed on kinetoplastids.

Besides energetic constraints, the availability of nucleic acid building blocks may be a selective constraint (for example, Hessen et al. (2010)). Several studies of chrysophyte genome sizes suggest availability of phosphorous and nitrogen along with the trophic mode (hetero-vs mixo-vs autotrophy) can be connected with the DNA content evolution in the lineage Čertnerová et al. (2025); Olefeld et al. (2018). Kinetoplastids lack the *de novo* purine biosynthesis pathway Butenko et al. (2020), so they can only obtain building blocks for DNA from external sources. Free-living kinetoplastids might therefore face a DNA building block shortage when food is limited. Concentrations of bacteria normally found in marine water are much lower than those that support maximum ingestion rates in heterotrophic nanoflagellates (to which free-living kinetoplastids belong) Tophøj et al. (2018). The assumption has been made that heterotrophic nanoflagellates generally face food limitation Tophøj et al. (2018), however, heterotrophic nanoflagellates can also feed on bacteria associated with solid particles or surfaces, involving more complex, not solely concentration-dependent, predator-prey interactions; for example, Erken et al. (2011, 2012); Kiørboe et al. (2004).

#### 2. Cell volume, replication and growth rates

The karyoplasmic ratio should be maintained within a certain range for the cell to proceed through the cell cycle Cavalier-Smith (2005). Consequently, the volume of the cell and the genome size may co-evolve, and many studies have discussed that DNA content generally correlates with both cell volume and division rates (for example, Elliott and Gregory (2015); Blommaert (2020); Cavalier-Smith (2005); Gregory (2001); Poulíèková et al. (2014); Šímová and Herben (2012); Malerba et al. (2020). According to the nucleoskeletal and nucleotypical DNA hypotheses, a natural selection for the cell volume or division (growth) rate could be reflected in the genome size Hof and Sparrow (1963); Cavalier-Smith (2005); Gregory (2001).

Based on the available data, the replication of kDNA and nuclear DNA occurs at a certain timing relative to one another, well-orchestrated throughout the cell cycle (for example, Woodward and Gull (1990); Bhattacharjee and Biswas (2023); Mensa-Wilmot et al. (2019); Cosgrove and Skeen (1970)). If there is a selective pressure towards shorter division times and/or smaller cell volumes, the sizes of the two genomes could co-evolve in complex feedback loops.

An inverse correlation between the intron density and the logarithm of generation time has also been reported Jeffares et al. (2006). It has already been proposed that the lack of introns in kinetoplastids could be associated with positive selection on the cell proliferation rate, but it implied another assumption of kinetoplastids being r-selected organisms Kostygov et al. (2024b).

#### 3. Adaptive versus neutral processes

According to the mutational burden/hazard hypothesis, the accumulation of non-coding DNA has weak deleterious effects, which would only be targeted by natural selection in species with sufficiently large effective population sizes Lynch (2002); Lynch and Conery (2003); Lynch (2006a,b). The mutational burden Lynch and Conery (2003); Lynch (2006a,b) or mutational equilibrium Petrov (2002) hypotheses can only be tested when the effective population size and mutation rates are known for the species. These parameters need to be measured experimentally and are largely unavailable for protists. Due to the lack of data, we are yet unable to evaluate the applicability of such hypotheses as the mutational burden/hazard Lynch and Conery (2003); Lynch (2006a,b), for free-living kinetoplastids.

Alternatively, experimental evolution and empirical measurement of some fitness parameters can be performed to see whether evolved lineages with smaller genomes have higher fitness. This approach does not require knowledge on population genetics parameters for a given protist, but still permits testing whether genome size is associated with fitness and can be a target of natural selection. Such studies have been performed on the green alga *Dunaliella* Malerba et al. (2020), and show that fitness of lineages with smaller genomes increases. Experimental designs similar to those described in Malerba et al. (2020) are suitable in principle for further studying DNA content evolution of free-living kinetoplastids.

### 4.2 Evolution of genome size in Metakinetoplastids

Kinetoplastids as a clade are divided into two sister lineages: prokinetoplastids and metakinetoplastids. Within metakinetoplastids, three monophyletic clades: trypanosomatids (well-studied parasitic species), and parabodonids then eubodonids as sister to trypanosomatids, branch from within multiple lineages of so-called neobodonids, which are, in fact, a paraphyletic assemblage of several subclades. Among trypanosomatids, haploid nuclear genome sizes range roughly from 14 to 50 Mbp Kostygov et al. (2024a). The only free-living metakinetoplastids with sequenced genomes, the eubodonid *Bodo saltans* (GCA_001460835.1), and the neobodonid *Cruzella marina* (GCA_977035905.1) have haploid assembly sizes of rougly 40Mbp and 100Mbp, respectively.

In this study, we obtained genome size estimates for free-living kinetoplastids from various deep-branching lineages of ‘neobodonids’, ranging roughly from 33 to 74 Mbp (which would make *Cruzella marina* the free-living metakinetoplastid with the largest genome, based on its 100 Mbp genome assembly). As a side effect of addressing our main hypothesis, our study informs genome size evolution in metakinetoplastids, accounting for the full diversity of the lineage, not only the currently highly sampled trypanosomatids. It is obvious that nuclear and kDNA contents are not conserved within major lineages of metakinetoplastids. Given their phylogenetic position, it is possible that trypanosomatids as a lineage have undergone an overall reduction in nuclear DNA content; perhaps this preceeded the origin of trypanosomatids and was already the case for the common ancestor of trypanosomatids and eubodonids. Taxon sampling of the free-living lineages remains insufficient to reconstruct DNA content changes with precision, and the aforementioned suggestions must remain speculative. The method proposed in this study can be applied in the future to measure the kDNA and nuclear DNA content of additional free-living metakinetoplastids. Even without transcriptome sequencing data, it can provide insights on DNA content evolution in this lineage, and its possible connection with ecological features, including parasitic lifestyles. Populations of parasitic species (like trypanosomatids) are frequently facing bottlenecks Pérez et al. (2019); Oberle et al. (2010), and bottlenecks increase the impact of genetic drift (for example, Katju and Bergthorsson (2019)). Therefore, signatures of selection on the DNA content, if any, may be observed less consistently in this lineage (which was one of the reasons trypanosomatids were not included in this study).

### 4.3 Advantages and pitfalls of methodology used for DNA content estimation

Sequencing of protist genomes remains infamously challenging. Difficulties in obtaining enough material, presence of prey or symbiotic organisms inseparable from a target species, and lack of culturing approaches are only a few of many factors that complicate the process of obtaining high-quality protist genome assemblies (for example, Schoenle et al. (2025)). The nuclear genome sequencing data currently available for the free-living (non-axenic, and putatively bacterivorous) kinetoplastids is severely limited: only 2 genomes (GCA_001460835.1 and GCA_977035905.1) were publicly available as of summer 2026. Precise estimates of kDNA content also cannot be easily retrieved by sequencing approaches, as the organisation of kDNA, where minicircles contain conserved and variable regions would result in assembly difficulties, although it is not impossible (see Wang et al. (2025)). Using fluorescence quantification-based approach, only 4 estimates have been produced specifically for the kDNA content of kinetoplastids Lukeš et al. (2018). Approximate kDNA content could be infered based on kDNA molecular weight (summarised for different species in Borst and Hoeijmakers (1979)), or the kDNA organisation and plasmid copy number (based on high-quality kDNA assemblies and subsequent read mapping for the copy number evaluation, for example, Wang et al. (2025)), but the accuracy of such estimates cannot be assessed.

To sum up, precise DNA content estimates were previously only available for a small number of free-living kinetoplastid species, and direct sequencing of the genome is not always a straightforward approach for quantifying both nuclear and kDNA content. Alternatively, DNA content can be estimated by staining nuclei with a fluorescent dye which binds DNA in a quantity-dependent manner. The fluorescence intensity of stained nuclei (which is proportional to DNA content) can then be assessed by flow cytometry or fluorescent microscopy (for example, Munyenyembe et al. (2021); Wheeler et al. (2012); Broemsen et al. (2021); Parrow and Burkholder (2002); Zubáčová et al. (2008); Noirot et al. (2002); Parfrey and Katz (2010); Mukherjee et al. (2009); Von Dassow et al. (2008); Timmons et al. (2024); Kuroiwa et al. (2015))

Although the number of cells screened by fluorescent microscopy is expected to be lower compared to flow cytometry, previous studies have shown that estimates obtained by these two methods are comparable in accuracy Wheeler et al. (2012); Broemsen et al. (2021), furthermore, developing (semi-)automated image analysis pipelines can substantially increase the cell numbers screened by microscopy (see below).

Common flow cytometry techniques have considerable limitations when applied to kinetoplastids: they do not allow discrimation between the kDNA and nuclear DNA, and further difficulties can arise when working with polyxenic cultures of presumably bacteriovourous small flagellates. In fact, many protists from diverse lineages cannot be reliably screened for nuclear DNA content by flow cytometry: DNA of their endosymbionts/prey/organelles significantly influences total fluorescence measurements Broemsen et al. (2021); Kuroiwa et al. (2015) and cells can be multinucleated Munyenyembe et al. (2021); Parfrey and Katz (2010); Mukherjee et al. (2009). Imaging flow cytometry, which combines cell sorting and imaging of cells could in theory distinguish between kDNA and nuclear DNA fluorescence signal, and protocols for such techniques were recently established for a parasitic axenic kinetoplastid *Leishmania mexicana* Howell et al. (2024). However, to our knowledge, these approaches have not been tested on non-axenic bacteriovorous kinetoplastids, and they also require specialised equipment.

Some past studies have estimated both the kDNA and nuclear DNA content of kinetoplastids based on fluorescent microscopy data Lukeš et al. (2018); Wheeler et al. (2012), and analogous approaches have been used to quantify nuclear DNA content of protists from other lineages Broemsen et al. (2021); Munyenyembe et al. (2021); Parfrey and Katz (2010); Timmons et al. (2024); Mukherjee et al. (2009); Zubáčová et al. (2008); Kuroiwa et al. (2015).

Previous fluorescence intensity quantifications were performed manually in all Broemsen et al. (2021); Munyenyembe et al. (2021); Parfrey and Katz (2010); Timmons et al. (2024); Mukherjee et al. (2009); Zubáčová et al. (2008); Kuroiwa et al. (2015) but one study Wheeler et al. (2012) (the same method used in Lukeš et al. (2018), and part of an analogous approach was later described in Wheeler (2020)). The image analysis plugin from Wheeler et al. (2012) was initally developed to detect DNA content differences from images containing cells of single species, without bacteria, and typically with a known genome sequence. Its algorithm uses typical DAPI/PI intensities of bright regions on the image as reference values for discriminating kDNA from nuclear DNA. This otherwise useful plugin is suboptimal for analysing images from our study: free-living neobodonids have kDNA less condensed than that of trypanosomatids Simpson et al. (2002); Lukeš, J., Lys Guilbride, D., Votypka, J., Zíková, A., Benne, R., and Englund, P. T. (2002), in our isolates, it is located adjacent to the nucleus, and, because their genome sizes are unknown, are present on the same slide together with several control species of various morphologies and sizes (for large species, z-stacks are required to fully capture nuclei).

The experimental and data analysis frameworks released with this study can be applied to other protist lineages, and would be especially useful for those where several compartments with comparable DNA content are present (for example, species with nucleomorphs). The open-source plugin presented in this study https://github.com/wisstock/kinetofluo-napari allows manual verification and curation of the automatically identified DNA-containing compartments. Sripts for the subsequent statistical analysis of raw intensity measurements are openly available as well: https://github.com/beaplab/kineto_stats/blob/main/FINAL_neat_knit_clustered_FUNCTION-BASED.Rmd and https://github.com/beaplab/kineto_stats/blob/main/FINAL_Post-process_NEW_by_distr.Rmd.

Some important considerations should be taken into account when applying the technique from this study:

1. Depending on the imaging setup, there can be lower and upper limit of DNA content that can be reliably quantified.
2. For validation in which DNA content of one control species is predicted based on the linear model inferred from other controls, the number of control species should be =*>* 3. Furthermore, when control species with the smallest and largest DNA content are treated as unknowns, their predicted DNA content can deviate from the sequenced genome size, because the fluorescence intensity values would be below/above the range of calibration.
3. Cells where the two stained compartments overlap have to be manually identified. If skipped at the manual species annotation step, they will be omitted in further analysis. In cases of morphologies which almost always result in overlapping compartments when imaged, the method is poorly applicable (for example, too many prey bacteria in cytoplasm, polykinetoplast kDNA organisation).
4. Optimal subsetting parameters may differ depending on the extent of technical errors during slide preparation and image acquisition. The dataset presented in this study had negligible error within and among biological replicates Figure S4; however, microscopy setups may differ and it is advisable to identify the optimal way of subsetting data for every individual experimental setup.
5. Different species imaged on the same slide should preserve some morphological features after fixation, so they can be discriminated from one another. Alternatively, control and test species can be imaged on seperate slides. In this case, a different statistical framework that accounts for errors in technical replicates would be necessary.

## 5 Summary and Outlook

We established an open-source framework for estimating DNA content of different cell compartments based on quantitative fluorescence imaging. It is applicable to heterotrophic protists, such as free-living metakinetoplastids, that are co-cultured with their microbial prey. Using this framework, we produced fluorescence-based estimates of kDNA and nuclear DNA content, together with transcriptome-based estimates of nuclear protein-coding DNA content and ploidy, for 10 phylogenetically diverse free-living kinetoplastid isolates.

We hypothesised that, in the process of evolution, different free-living kinetoplastids might have come to the increase in the proportion of cell DNA content represented by kDNA, and this might be connected with a reduction in the fraction of non-protein coding nuclear DNA (“compact” nuclear genomes). We compared how the fractions of the kDNA content and non-protein-coding nuclear DNA content relative to the total cell DNA content vary across the isolates examined, and discovered an inverse relationship between these two parameters. This might mean that higher kDNA content is associated with lower non-protein-coding nuclear DNA content, and that DNA contents of these two compartments could co-evolve.

The taxon sampling of our study covers the diversity of free-living metakinetoplastids in breadth, but not in depth. However, with our framework, data from more isolates could be easily added to the current set, given more time and resources. With more isolates sampled, the hypotheses on kDNA and nuclear DNA content co-evolving in this lineage could be further verified, or, conversely, disproved. If confirmed, very dense taxon sampling may provide sufficient data to robustly infer the directionality of evolutionary change in selected clades (for example, did a given clade experience an increase in kDNA, leading to a decrease in non-coding nuclear DNA, or did a reduction in kDNA mass relax constraints on non-coding nuclear DNA).

Finally, our framework can be applied to other protists to quantify and study the evolution of DNA content in various lineages. It would be especially valuable for studying DNA content of multiple compartments present within the same cell: protists with intracellular symbionts or parasites, or with multinucleated cells, as well as other protist lineages with remarkably large organellar DNA content.

Most importantly, in this study we attempted a hypothesis-based, experimental data-driven project on the evolution of protist lineages that are otherwise undersampled and neglected, and we hope that future studies can make use of our data and scripts, or take broadly similar approaches to different questions in evolutionary protistology.

## 6 Data availability

The image analysis plugin is available on GitHub https://github.com/wisstock/kinetofluo-napari. Raw imaging data, as well as graphical outputs of image analysis (manually curated) are deposited to BioStudies database Sarkans et al. (2018) and will be available under accession S-BSST3293. All files and scripts used for transcriptome analysis, downstream analysis of fluorescence intensity values and ploidy estimation are available either via the GitHub repository https://github.com/beaplab/kineto_stats, via Figshare at DOI: 10.6084/m9.figshare.29088251, or as the Supplementary Material in Zavadska et al. (2025).

## 7 Acknowledgments

We are grateful to Koryu Kin, Gonzalo Bercedo Saborido, Núria Ros-Rocher and Guifré Torruella i Cortés for sharing strains of protists with known genome sizes (not all of them were used for the final prediction since some had uncertain ploidy). We thank Igor Kostikov, Lada Isakova and Michelle Leger for valuable comments on the draft text.

## 9 Supplemental information

### Section S1.1 Relationship between the kDNA and non-protein-coding DNA content fractions under alternative estimates of proteome lengths

**Figure S1:**
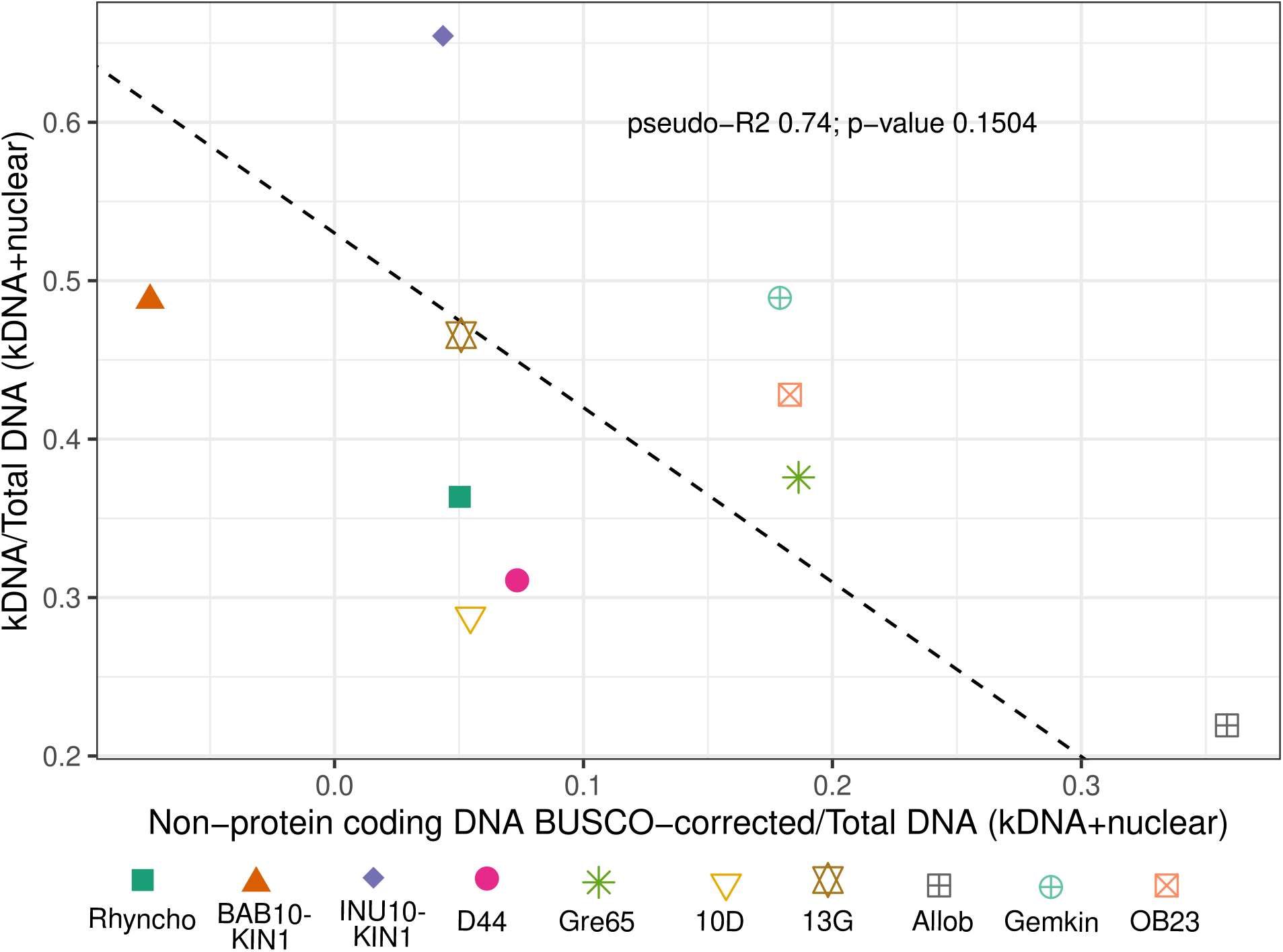
Non-protein-coding nuclear and kDNA content relative to the total cellular DNA content of each isolates. The estimate of non-protein-coding nuclear and kDNA content was corrected for BUSCO completemess of non-redundant proteomes. See Section S8.8.3 for the details on the correction for completeness; we note that such correction represents rather bold estimate, and BUSCO completeness is not the most suitable measure for assessing completeness of free-living kinetoplastid proteomes: see below.

### Section S2.2 DNA content estimates of 10 free-living kinetoplastid isolates

**Table S1:**
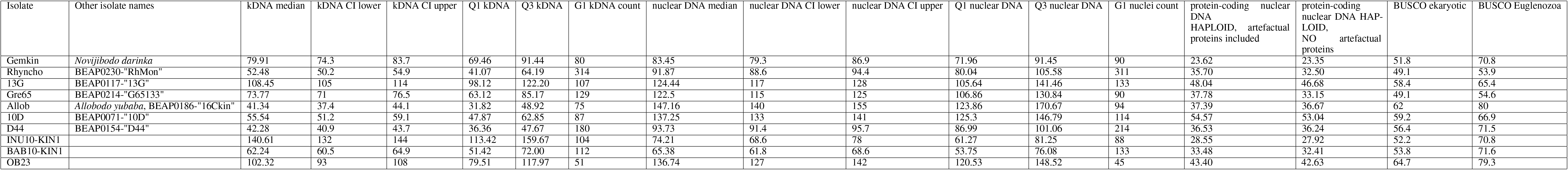
kDNA, total nuclear DNA content, and protein-coding content of 10 free-living kinetoplastid isolates. All DNA content values are in Mbp. For fluorescence-based estimates (columns 2-6 and 8-12), DNA content of **G1, diploid** compartments is shown. For transcriptome sequencing-based estimates, length of **haploid (G1)** protein-coding nuclear genome is shown. “Q1/Q3 kDNA/nuclear DNA” columns show 1st and 3d qunatile of predictes DNA content distributions (where predicted DNA content of single compartment of one cell represents a datapoint). “kDNA/nuclear DNA CI lower/upper” colums show the upper and confidence interval of the median of predicted DNA content.

### Section S3.3 Culturing conditions and information on isolates for control species with known genome size and kinetoplastids

Marine kinetoplastid isolates (Rhyncho, D44, 13G, Gre65, 10D, 13G, Allob, OB23) were cultured in “classical” RS medium with the same concentration of Nutrient component https://mediadive.dsmz.de/medium/P5, diluted 1:100 in of the Non-Nutrient component https://mediadive.dsmz.de/medium/P4 (the latter mimicking seawater, 36ppt). The other three kinetoplastid isolates were also grown with Nutrient component of RS medium in 1:100 dilution as carbon source, while the Non-Nutrient component was substituted with mineral-based media of salinity and pH that corresponds to the optimal growth conditions of the isolate. For Gemkin, this was either “classical” RS medium used for other marine strains, or: CR medium, salinity 20ppt Gigeroff et al. (2023) or Non-Nutrient component of RS diluted 1:1 in distilled water. For INU10-KIN1 and BAB10-KIN1, 3N Bold’s Basal Medium Bischoff (1963) and 1:100 diluted Nutrient component with distilled water as a solvent were used. For Gemkin, this was: CR medium, salinity 20ppt Gigeroff et al. (2023) or Non-Nutrient component of RS diluted 1:1 in distilled water. For INU10-KIN1 and BAB10-KIN1, 3N Bold’s Basal Medium Bischoff (1963) and 1:100 diluted Nutrient component with distilled water as a solvent were used.

Additional information on kinetoplastid isolates used in this study (transcriptome sequencing data, live imaging, isolation sites, etc.) can be found in Zavadska et al. (2025).

**Table S2:**
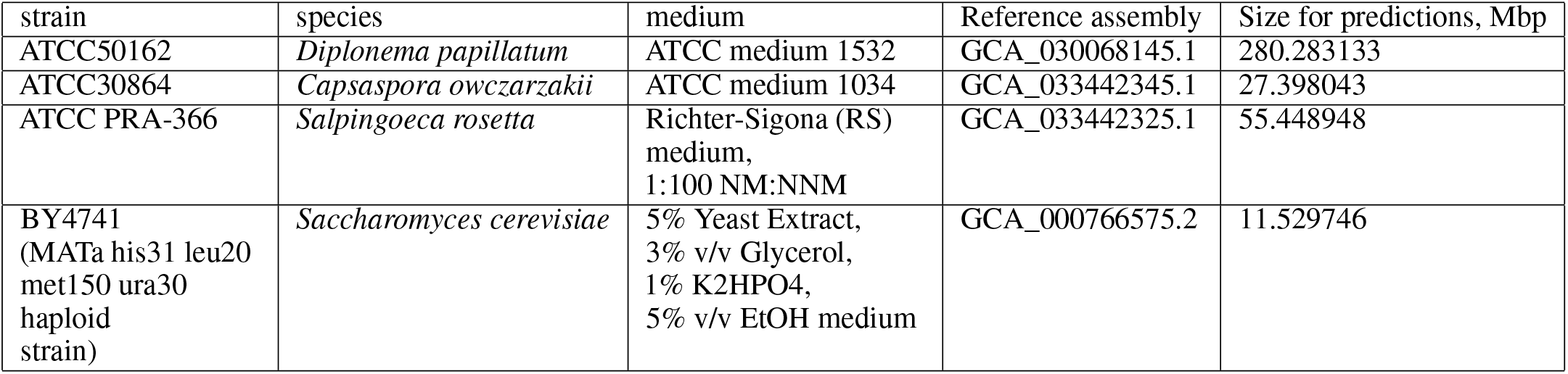
Information on strains with known genome size used as a control. Strain, species, culture medium, reference GenBank IDs and genome size used for calculations are listed. Laboratory strains derived from *Salpingoeca rosetta* strain ATCC PRA-366 are known to be both haploid and/or diploid populations Levin and King (2013); our population was predominantly haploid, according to the Sanger sequencing of polymorphic loci (see below). The genome size in the table corresponds to than of haploid assembly, 1n (“G1”) cells.

#### Section S4.4 Stain concentrations

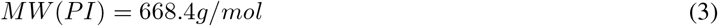

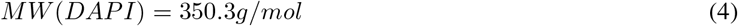

The concentrations of dyes in the staining solutions:

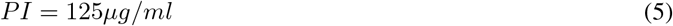

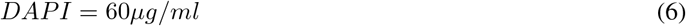

Assuming at least 10 µl of liquid remains on 24mmx24mm coverslip (usually it is around 12-15 µl), the number of PI molecules will be:

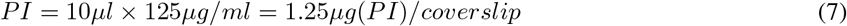

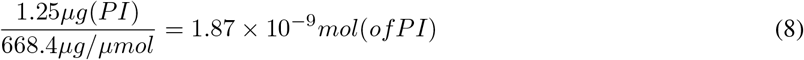

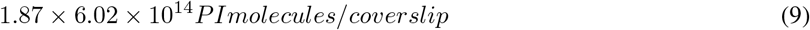

The number of DAPI molecules will be:

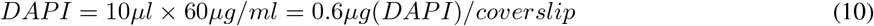

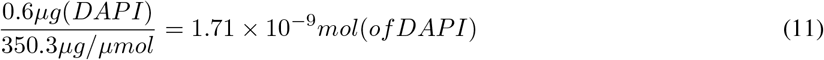

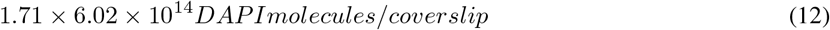

Assume the nuclei of *Diplonema papillatum*, the species with the largest genome size used in this study, are squares of 5µmx5µm, they cover all the surface of the 24mmx24mm coverslip, and all of them are in the G2 phase of the cell cycle, containing 560Mbp of DNA each. The hypothetical maximum number of *Diplonema papillatum* nuclei which fits on the coverslip would be:

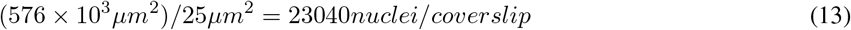

All nuclei that could be allocated on one coverslip (as a monolayer), would contain

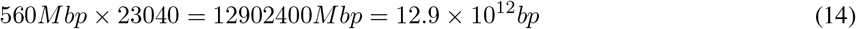

PI:DNA binding stoichiometry 1 PI molecule : 4-5 basepairs (as mono-intercalating dye; we take 4 as minimum) Lin et al. (2003)

DAPI:DNA binding stoichiometry 1 DAPI molecule : 4 basepairs Larsen et al. (1989)

So, given the number of DNA basepairs which could be positioned on one coverslip (organised as a monolayer of square nuclei), the number of dye molecules in the staining solution in which the coverslip is immersed, and the stoichiometry of PI and DAPI binding with DNA, the number of molecules required to bind all the DNA on the coverslip, will be, for either DAPI or PI:

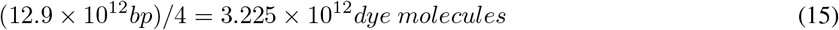

The number of both DAPI (1.03 *×* 10^15^) and PI (1.13 *×* 10^15^) molecules in staining solution exceeds the minimum required number of molecules, 3.225 *×* 10^12^, by a magnitude of more than two orders, even given the most unrealistic, worst-case estimates of the amount of DNA and staining solution present on the slide.

### Section S5.5 Image acquisition setup - interval between slices in z-stack

#### Section S5.5.1 Resolution in Z-plane

The axial resolution, or depth of field, of wide-field fluorescent microscopy can be estimated using the equation 16, where *n* is the refractive index of the sample, *λ* is the fluorescence wavelength, and *NA* is the numerical aperture of the objective.

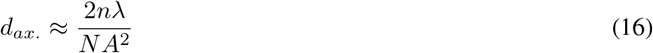

The estimated depth of focus for the emission maxima of DAPI (*λ_DAP I_ ≈* 465nm) and PI (*λ_PI_ ≈* 620nm) was 640 nm and 840 nm (17, 18).

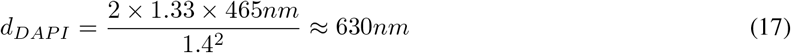

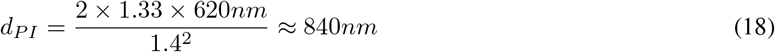

The interval of 1.2 µm between the individual focal planes exceeds the estimated depth of field, even for the red PI, and each z-stack contains 8 slices. This enables us to effectively collect signals from cells of various sizes that may be located at slightly different depths within the samples. Using a large interval also prevents signal overlap and possible overestimation of total signal values.

#### Section S5.5.2 Fluorescence intensities depending on the intervals between slices and the number of slices

To identify the optimal interval between slices, we acquired z-stacks with two intervals: 1.2 µm interval, 8 slices per stack (used in the final setting), and 0.24 µm interval, 37 slices per stack (the one estimated as “optimal” by Zen Blue image acquisition software). The stacks were acquired from the same slide to minimise the impact of technical error on the comparison.The images were analysed using the custom plugin; the fraction of out-of-focus slices which were dropped before the intensity estimation was constant regardless of the number of slices: i.e from 1 to 2 slices were normally dropped from 8-slice stack, correspondingly, 5 to 10 slices were dropped from 37-slice stack. The intensity counts obtained from both type of stacks were analysed as described in the main text.

The resulting distribution of raw fluorescence intensities are visualised in Figure S2. Although adding more slices in the stack contributes towards the overall higher intensities, the **relative differences in intensities** between different species/compartments and different cell cycle stages remains unaffected. Based on this observation, on the estimates presented above, as well as practical considerations (the size of the image files, time of acquisition, chance of bleaching while acquiring 37 slices, and system memory requirements for image processing), we selected the setting with 1.2 µm interval, 8 slices per stack.

**Figure S2:**
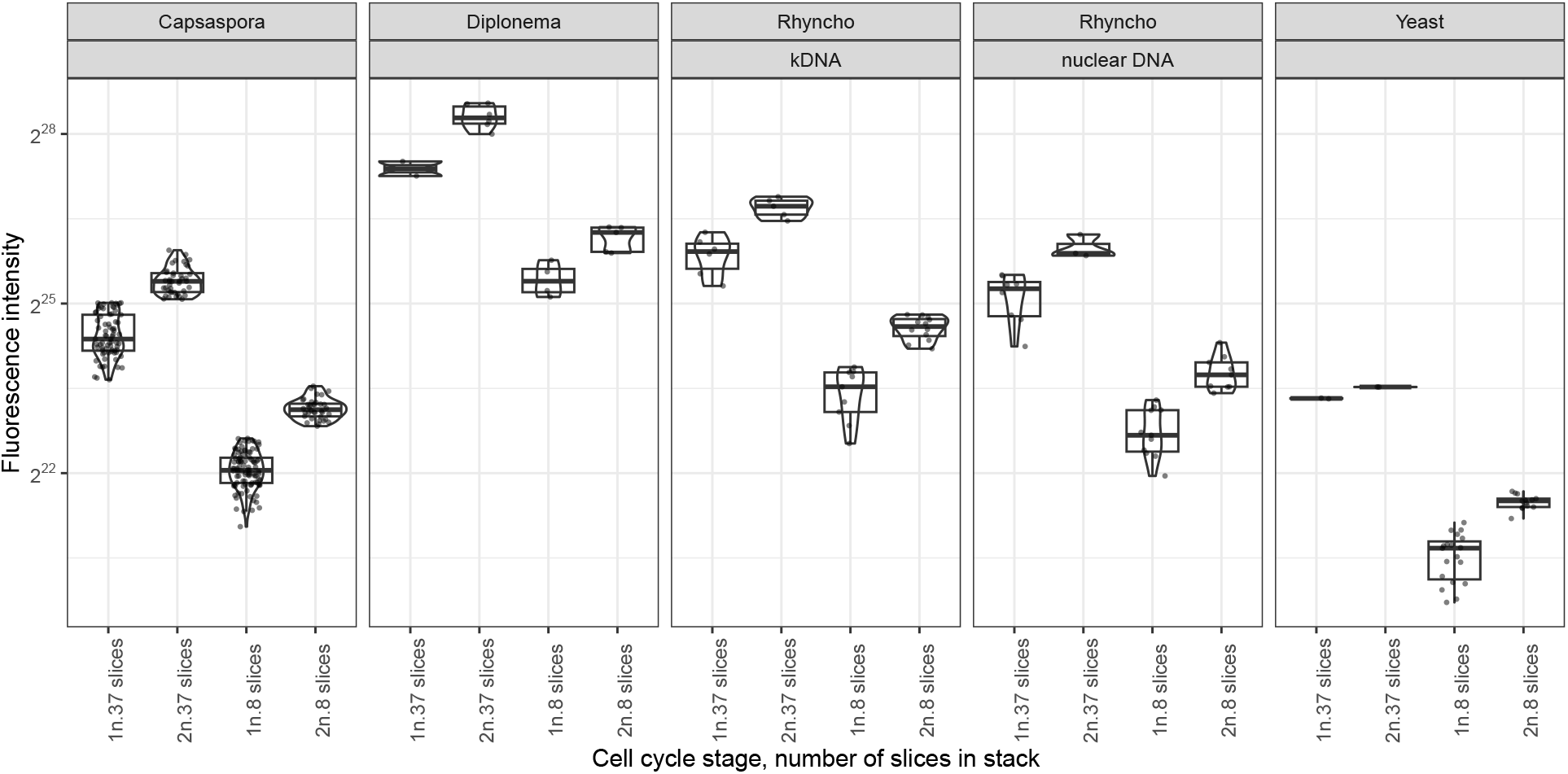
Distributions of fluorescence intensities depending on the number of slices and interval of the stack. Each facet represents values of fluorescence intensities (y-axis) obtained for a given compartment of a given species; each dot represents a measurement from one cell. On x-axis, the values are split depending on the assigned cell cycle stage (“G1” or “G2”) and the stack setting (“8 slices” or “37 slices”). Note the log2 scale.

### Section S6.6 Statistical analysis benchmark

#### Section S6.6.1 Subsets used in data analysis

The overall experimental design assumed obtaining measurements from several independent sets of cell populations (eukaryotic cell “passages”) of control plus test species. These sets were passed, grown, put on the same slide, fixed, stained and imaged together. These independent sets are biological replicates of the study. Within each biological replicate, technical replicates have been present in the majority of cases: two different slides with the same populations of cells were prepared.

As a result, the raw data used in the analysis, represented by the fluorescence intensities of nuclei or kDNA of individual cells, had the underlying structure of a technical replicate (= glass slide) *<* biological replicate (same populations of cells and same day/batch of cell fixation and staining) Figure S3.

**Figure S3:**
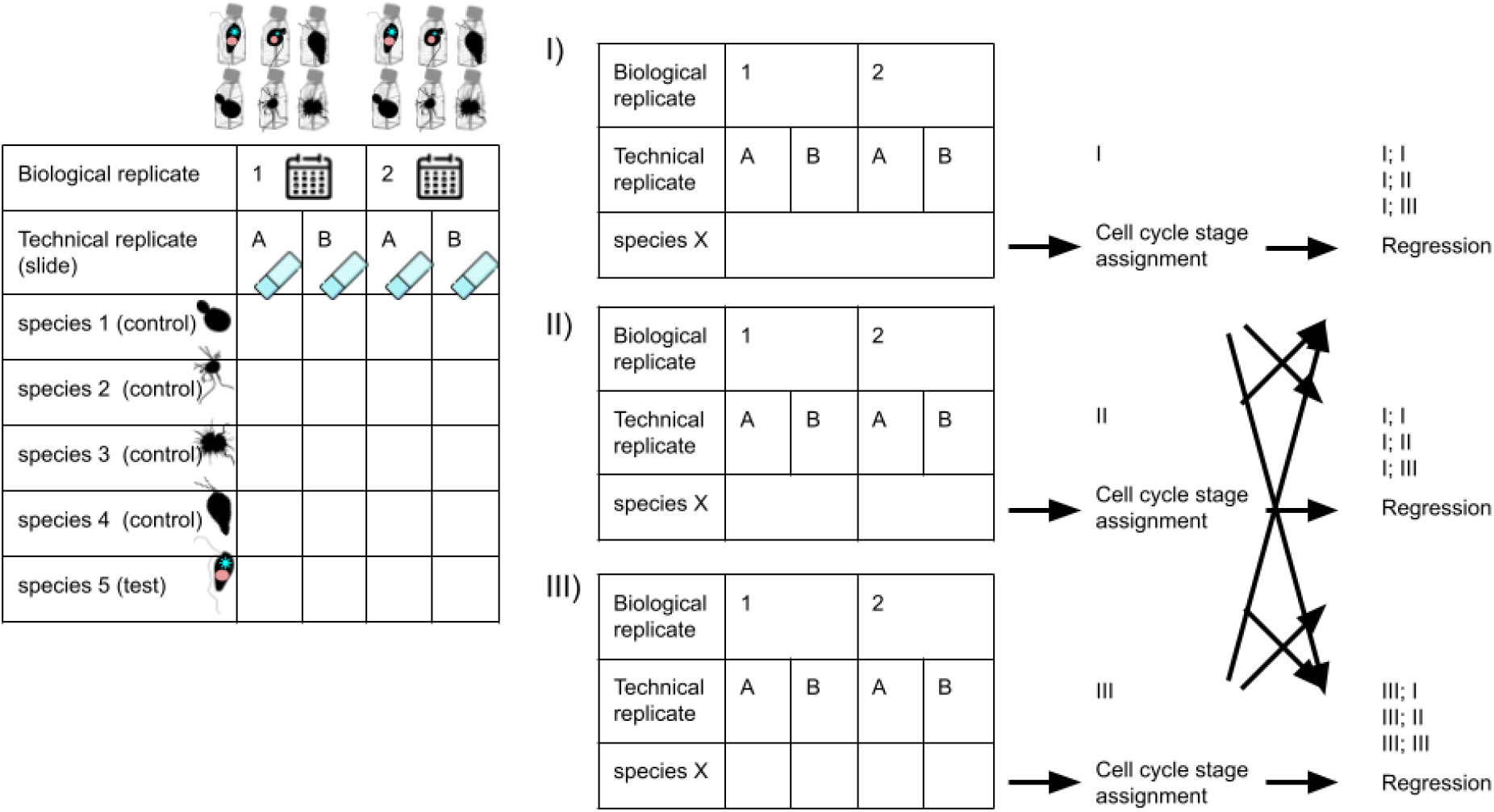
Schematic illustration of data structure and nature of replicates. Further description in the text.

Procedures of outlier filtering combined with the cell cycle stage assignment, and regression could be performed separately on different data subsets: separate sets of cells from every technical replicate (III in Figure S3), separate sets of cells from every biological replicate (II in Figure S3), or all cells with no subsetting (I in Figure S3). Furthermore, these sets could be combined: while outlier filtering with the cell cycle stage assignment is performed on a given subset (for example, all cells: “I”), the regression can be performed on a different one (for example, separately on values within each biological replicate: “II”). Using different subsets for different procedures can have advantages, given the nature of the data and the putative error sources. For example, combining cells of a given species from all biological replicates to predict the cell cycle stage can give more accurate result, as cell populations from different experiments might have the majority of cells shifted to only one stage in the cell cycle, G1 or G2.

All possible ways of subsetting data (12 combinations) were tested for the accuracy of genome size predictions the same way as described in the main text: DNA content of one out of three control species was predicted based on the regression infered using the rest three control species. The accuracy of predictions obtained for each subset combination is summarised in Figure S4 and Figure S5.

**Figure S4:**
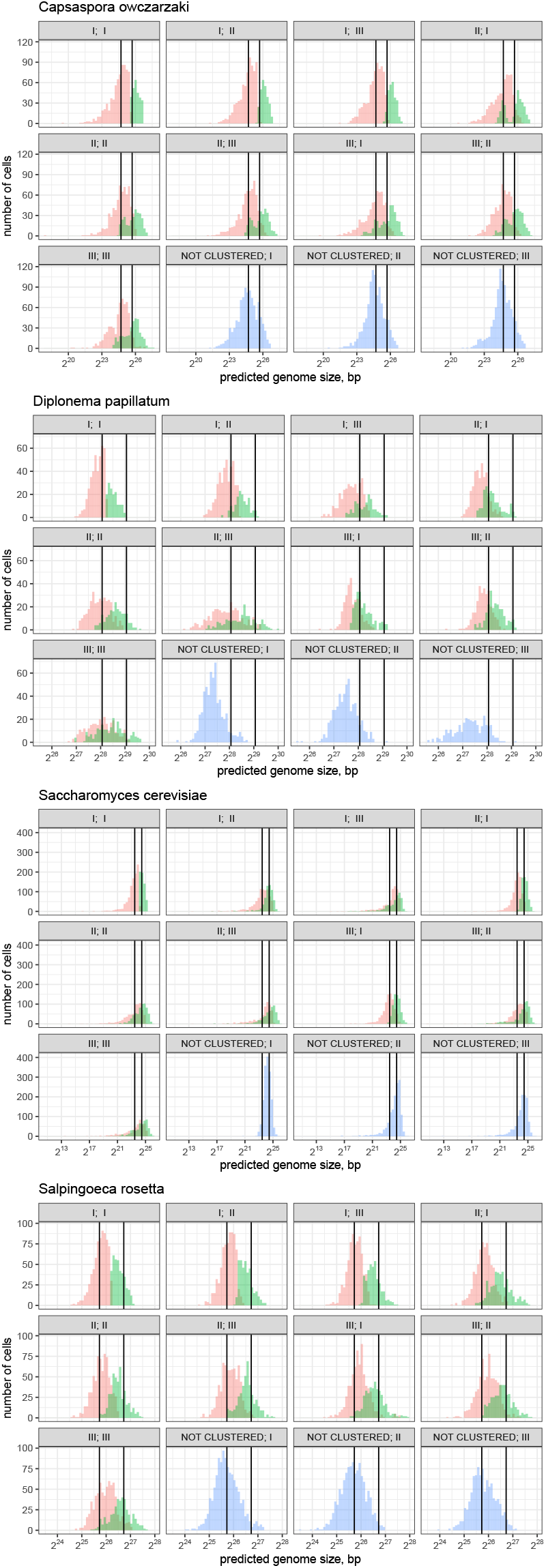
Distribution of DNA content predicted for the control species based on fluorescence intensities, compared to the expected genome sizes. FThe y-axis represents the number of G1 (red) and G2 (green) cells with a predicted DNA content (x-axis). The vertical solid lines indicate the actual sizes of control species genomes - for the G1 and G2 cells; the distributions tend to center around the vertical line, indicating that the predicted genome sizes, on average, correspond to the real ones for control species. Facets represent data subsets used for outlier filtering combined with the cell cycle stage assignment (first number in the facet header), and for regression inference (second number in the facet header); to match the number with the type of subset, refer to the Figure S3 and the description in text.

**Figure S5:**
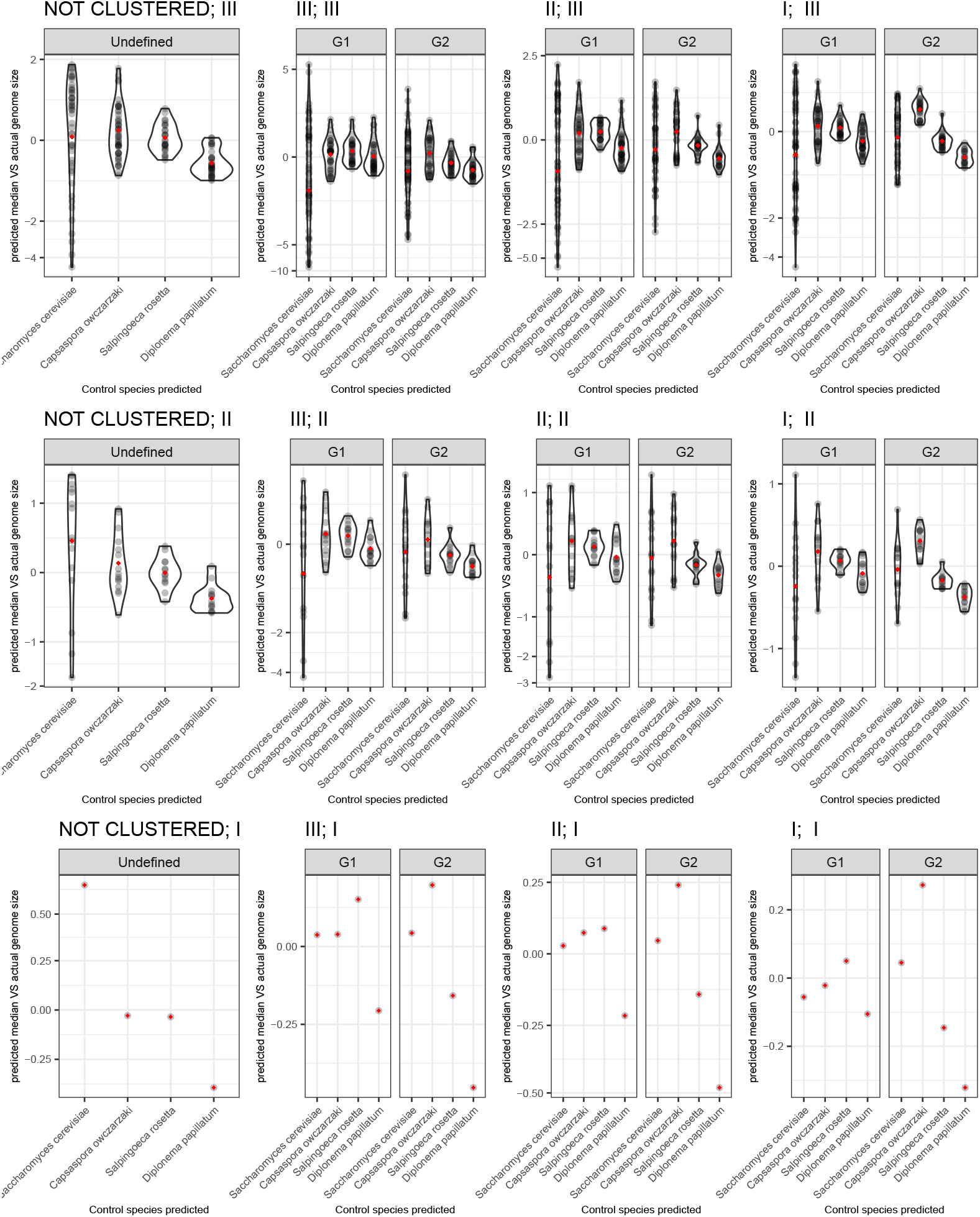
Difference of medians of genome sizes predicted for the control species from the expected genome sizes, as a fraction of the expected genome size. Each dot represents a median obtained from the predicted values of cells from either a single technical (top panel), biological (middle panel) replicates, or all the measurements taken together (bottom panel), from which the actual genome size was subtracted, and the difference is normalised by the expected genome size: (*Median_predicted_ − bp_actual_*)*/bp_actual_*. Red diamonds represent the means of medians from all biological/technical replicates, from which the actual genome size was subtracted, and the difference is normalised by the expected genome size. Facets represent subsets used for outlier filtering combined with the cell cycle stage assignment (first number in the facet header), and for regression inference (second number in the facet header); to match the number with the type of subset, refer to the Figure S3. Note different scales on the y-axis. The inverse hyperbolic sin transformation was used for y-axis values in this case, to efficiently represent both scattered values for *S. cerevisiae* and less scattered values from all other species.

Cell cycle stage assignment expectedly improves the accuracy of DNA content predictions, compared to the predictions which were done without the correction for the cell cycle stage (the latter are labelled as “NOT CLUSTERED; X”) Figure S4, Figure S5. Treating each technical replicate as a separate dataset (“III”) either for clusterization with outlier filtering or for regression may overall reduce the accuracy of prediction Figure S4, Figure S5. In majority of cases, for our data, there are no substantial differences in the distribution of predicted DNA content values across either combination of biological replicate subset (“II”) and/or no subsetting (“I”). To simplify downstream data analysis we decided to treat all biological replicates from experiments as a single dataset in (setting “I;I”) for obtaining the final estimates of DNA content presented in Figure 1, Figure 2, Table S1. Note that scripts used for statistical analysis have a built-in framework for running the validation of all vs all subsetting combinations and it is advisable to do it and select the optimal subsetting option for a new dataset, if replicating the method.

#### Section S6.6.2 Cell cycle stage assignment

In order to verify whether (i) cell cycle stage assignment using k-means clusterisation corresponds with the biological reality, and (ii) which subsetting option is the best for the cell cycle stage assignment, the experiment with blocking the cell cycle was performed.

The cell cycle was blocked in one of the control species, *Capsaspora owczarzaki*, using hydroxyurea, which has been previously used to arrest the cell cycle at G1 phase in this species Pérez-Posada et al. (2020); Sebé-Pedrós et al. (2013). For hydroxyurea treatment, protocols from Pérez-Posada et al. (2020); Sebé-Pedrós et al. (2013) were adapted with modifications: 5-day old split of *C. owczarzaki* culture, grown in ATCC medium 1034 with Fetal Bovine Serum (“FBS+”), was centrifuged at 11kg for 8min, and cell pellet was resuspended in 8ml of ATCC medium 1034, but without FBS, to prevent the aggregation of cells. 2ml of 0.5M Hydroxyurea (Sigma-Aldrich, Cat. No. H8627) stock (also in ATCC 1034 medium with no FBS) was added to the resuspended culture, final Hydroxyurea concentration being 100mM, and the culture was left in the flask, for a 24h incubation, prior to being harvested, fixed and stained by a procedure described in the main text. The control population of *C. owczarzaki* cells was grown in the same initial flask, but cells were resuspended with the 8ml of fresh ATCC 1034 medium, with addition of FBS, 24h prior to harvesting, fixation, and slide preparation. Both test and control populations of *C. owczarzaki* were harvested, fixed and stained on the same day, simultaneously, on separate slides, but stained in the same “batch“; cells of other control species in this batch originated from the same passage.

After obtaining the raw fluorescence intensities of hydroxyurea-treated and control *C. owczarzaki* nuclei, these intensity values were subject to the outlier filtering and cell cycle stage assignment, while included into different subsets of data (see Figure S3 and corresponding explanation in text). DNA content was predicted for these cells, using regressions inferred based on the data from the other three control species, as described above.

Figure S6 and Figure S7 illustrate that:

- Both raw intensities and predicted genome sizes are lower for Hydroxyurea-treated cells compared to the untreated ones.
- Majority of Hydroxyurea-treated cells are classified as G1 stage, regardless of the subsets of data used.
- However, for subsets II and III Figure S6, based on the raw intensity values, few Hydroxyurea-treated cells are still classified as G2; while for the subset I the number of incorrectly classified cells is lowerFigure S6.

We can conclude that outlier filtering and cell cycle stage assignment may be prone to artefacts when performed within individual subsets of biological (“II”) or technical (“III”) replicates: if cells of a given cell cycle stage are overrepresented in the subset, the clusterisation by k-means could incorrectly classify cells. Instead, when the measurements from all cells across all replicates (“I”) are taken into account, the overrepresentation of a single cell cycle stage in one replicate does not influence the outcome of cell cycle stage assignment. Subsetting data for the cell cycle stage assignment procedure could slightly influence the accuracy of DNA predictions, as ilustrated in Figure S4. In the majority of cases, performing outlier filtering and cell cycle stage assignment on all cells from all replicates subtly increases the accuracy of predictions Figure S4. From a biological perspective, every biological replicate can differ slightly in terms of the number of cells in each cell cycle stage, and we are unaware on the details of this dynamics.

To sum up, performing outlier filtering and cell cycle stage assignment on all cells of a given species from all replicates at once (subset “I” Figure S3) is the best option, which has been selected for obtaining the final predictions of DNA content. Yet, the number of incorrectly classified Hydroxyurea-treated cells remains relatively low even when the cell cycle stage assignment is done for separate technical (“III”) or biological (“II”) replicates. There is also a possibility that some fraction of cells just escapes the cell cycle block. So for data from other experiments, depending on the extent of technical errors and cell cycle dynamics of cultures, subsetting measurements by technical or biological replicates should still produce realistic classification.

**Figure S6:**
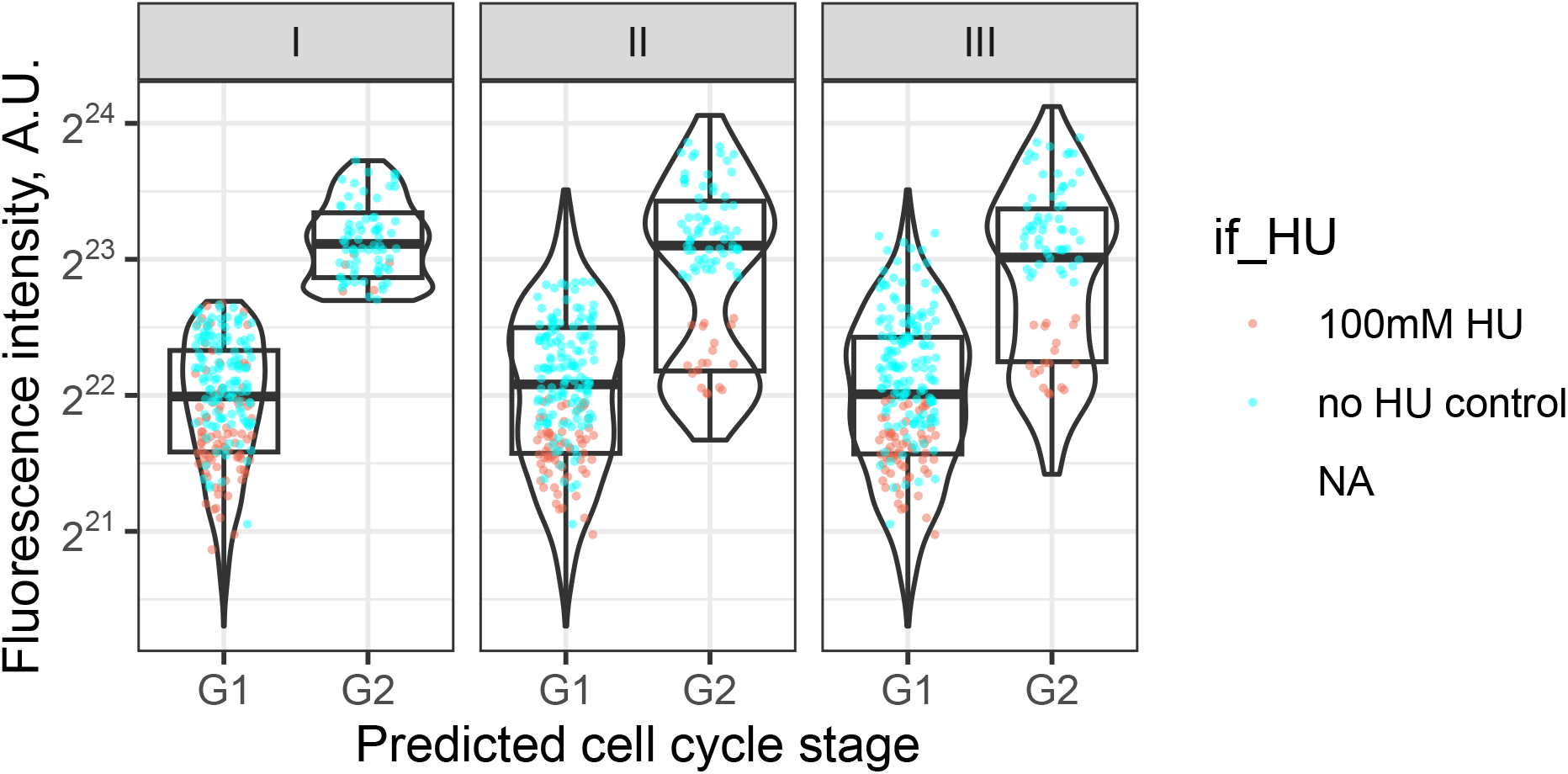
Distributions of raw fluorescence intensities of *C. owczarzaki* nuclei, across cell cycle stages assigned, for different subsets used for classification. Violin and boxplots represent the overall distribution of intensities from **all cells** measured across **all biological replicates**. The dots in red and represent the intensities measured for Hydroxyurea-treated cells and the dots in blue - intensities from control (non-treated) cells from a given biological replicate. The majority of red dots should be G1 cells, and are expected to be classified as such, while blue dots are expected to be both G1 and G2 cells. Facet headers show the subset of cells used in outlier filtering and cell cycle stage assignment procedure; to match the number with the type of subset, refer to the Figure S3. Note that the total number of cells might differ between subsets, as some values can be discarded as outliers in one subset, but not in the other.

**Figure S7:**
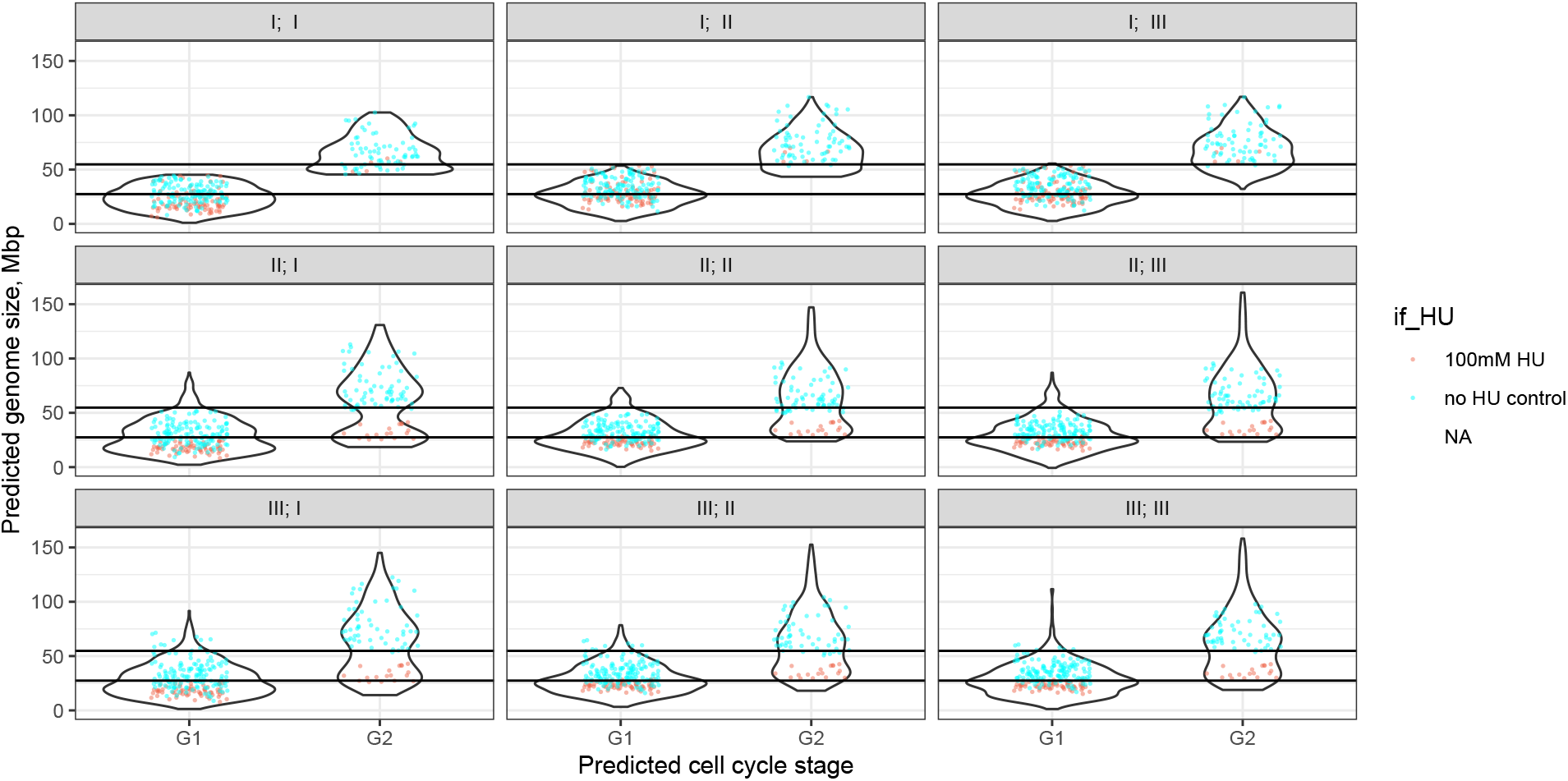
Distributions of predicted DNA content of *C. owczarzaki* nuclei, across cell cycle stages assigned, for different combinations of subsets. Violin and boxplots represent the overall distribution of predicted genome sizes from **all cells** measured across **all biological replicates**. The dots in red and represent the intensities measured for Hydroxyurea-treated cells and the dots in blue - intensities from control (non-treated) cells from a given biological replicate. The majority of red dots should be G1 cells, are expected to be classified as such and to have predicted DNA content closer to that expected for G1 cells, while blue dots are expected to be both G1 and G2 cells. Facet headers show the subset of cells used in outlier filtering and cell cycle stage assignment procedure (first number) and for regression inference (second number); to match the number with the type of subset, refer to the Figure S3. Note that the total number of cells might differ between subsets, as some values can be discarded as outliers in one subset, but not in the other.

### Section S7.7 Ploidy

The verification of ploidy levels was done in two steps: (i) transcriptomic read mapping against the predicted CDS, SNP calling and filtering by mapping quality and estimating allele frequency distribution and (ii) experimental verification of mapped SNPs by amplifying single-copy regions and Sanger sequencing resulting amplicons.

#### Section S7.7.1 Verification of ploidy - analysis *in silico*

The mapping of SNPs was performed for the transcriptomes of the species used in this study, and for the transcriptomes of various protist species with known ploidy. The species with known ploidy served as a control of whether the SNP detection based on **transcriptomic** read mapping works correctly and can be a good predictor of an organism’s ploidy. For control, we selected Illumina SRA studies of various protist species, from various supergroups, optionally with datasets from both haploid and diploid life stages of the same species. The raw reads for each control species were downloaded from NCBI, and assembled *de novo* using RNAspades v3.13.1 Bushmanova et al. (2019), default parameters (all combinations of read orientations and stranded vs unstranded modes were performed on all the reads; the proteins were predicted from the assemblies using TransDecoder v5.0.1 https://github.com/TransDecoder/ TransDecoder with the default settings, and the best assembler settings settings were identified in each case based on the number of complete BUSCO genes in assemblies produced by every setting Manni et al. (2021)).

See https://github.com/beaplab/kineto_stats/blob/main/GitHub_PLOIDY/mapping/Ploidy_Bioinf_assemblies.xlsx for the details on the species and assembly settings used as controls in this section.

**No redundancy removal was applied to CDS sets**, for both the control species and free-living kinetoplastids (we used the TransDecoder output before the CD-hit filtration was applied).

Raw reads were mapped against the CDS from transcriptome assemblies with bowtie2 v2.4.4 Langmead and Salzberg (2012), maximum fragment length (“-X” flag) set to 2000, other parameters default, and sorted with samtools v1.13 Danecek et al. (2021).

The mapped reads were filtered to retain only those with mapping quality >25 (“samtools view -b -q 25”). This mapping data was used as a source of information for finding PCR target genes which are expected to have biallelic SNPs (see the next section). The statistics on the mapped reads, filtered by quality, and the fasta of non-redundancy reduced CDS sets are available on the Figshare at DOI: 10.6084/m9.figshare.29088251.

To further check the global patterns of SNP frequency distributions, biallelic SNPs were called and coverages of major and minor alleles were estimated using a combination of bcftools commands mpileup (“-a AD,DP -Ou” flags) and view (“-V indels -Ov -o” flags). Variants were called while ignoring indels, filtering for a minimum read depth of 5, a minimum of four alternate bases before calling a SNP, accepting only bi-allelic SNPs, and only accepting SNPs with a MAF(=minor allele frequency) of 2%, and a maximum missing data of 50%.

For visualisation on Figure S8, the annotated SNPs were filtered so that only those with the sum of minor+major allele coverage >12 are plotted.

The majority of control species, with a few exceptions, that are known to be diploid show the peaks of frequency ratios of two alleles centered around 1 Figure S8, as would be expected for diploid genomes (and now, this has proven to be the case for the majority of diploid transcriptomes). In contrast, the control haploid species do not show a frequency ratios peak around 1; instead, they are distributed equally Figure S8. **All of the kinetoplastid species from this study show peaks of frequency ratios of two alleles centered around 1, similat those of control diploid species, which strongly suggests that they are diploid**.

The scripts for obtaining and visualising estimates from this section are available on Github https://github.com/beaplab/kineto_stats/tree/main/GitHub_PLOIDY.

We also validated the presence of SNPs experimentally: see below.

**Figure S8:**
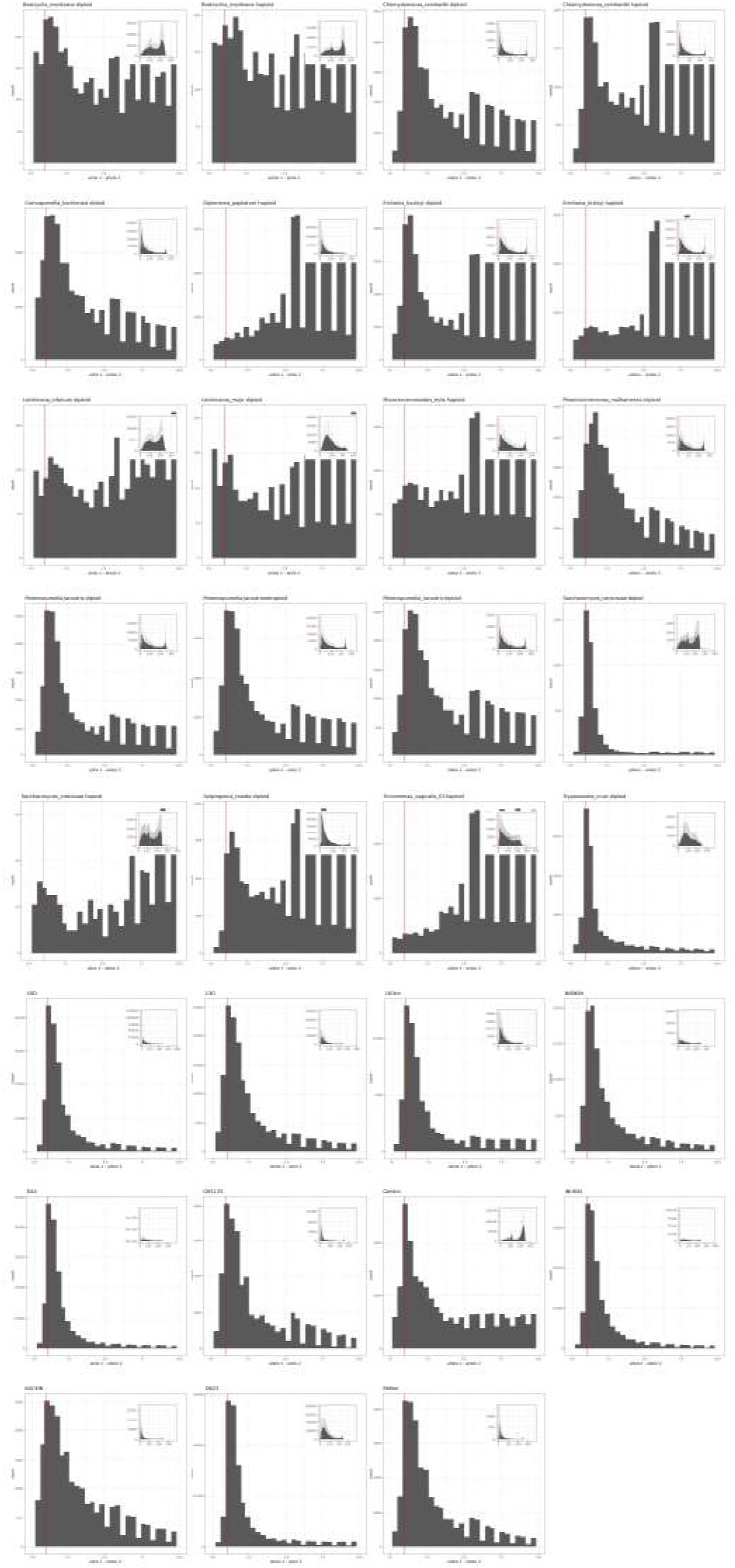
Distributions of major and minor allele frequency ratios in SNPs detected by transcriptomic read mapping. Each facet represents the distribution for each species; for the control species with known ploidy, the ploidy is also indicated. The red line corresponds to the ratio of 1: where the peak of distribution is expected for the diploid organisms. The distribution of the whole range of ratios is shown at the right upper corner of each facet, while large plots indicate the distribution of ratios ranging from 0 to 10. Frequency ratios higher than 10 mean that SNPs are less likely to result from the presence of multiple alleles (they could be artefacts of data analysis, sequencing, mutations in the process of transcription, etc).

#### Section S7.7.2 Verification of ploidy - PCR and Sanger sequencing

To ensure that the SNPs originate from the different alleles of the same gene and not from multiple copies of the same gene, we used the genes from the euglenozoan BUSCO (v.10) gene set as candidate targets for PCRs. From these genes, we selected the ones which would be (i) present as “Complete” and “Single-Copy” (not “Duplicated“!)in both redundancy-reduced and original proteomes of at least 8 (>70%) species from the study, and (ii) present as “Single-Copy” in the biggest number of Kinetoplastids that originally included in the euglenozoan BUSCO dataset (v.10). For each species, genes satisfying both criteria were additionally checked for the coverage, and presence of SNPs in high-coverage regions other than the end/start of the mapped reads, using the Interactive Genome Viewer software Robinson et al. (2011). Primers were designed for the selected genes, with the primer3plus web tool https://www.primer3plus.com/, so that the amplicons would cover as many polymorphic regions as possible. The list and sequences of genes and primers used, expected amplicon lengths, etc. are available on GitHub https://github.com/beaplab/kineto_stats/tree/main/GitHub_PLOIDY/PCRs.

In addition to the kinetoplastid species used in this study, we verified the ploidy of the *S. rosetta* strain used as a control; for this species, we selected two random genes which would have SNPs and sufficient coverage (according to the transcriptome mapping described in the section above), and which would not produce any significant matches with other genes of *S. rosetta* upon the BLAST search. Haploid strain of *S. cerevisiae* (the one also used as a control for staining), and a diploid sample of human were used as a haploid and diploid controls.

Total DNA was extracted from monoclonal cultures of kinetoplastid isolates and from controls with DNeasy Power-Soil Pro Kit (QIAGEN, cat. no. 47017, Hilden, Germany). Target regions were amplified using Phusion High-Fidelity DNA polymerase (ThermoScientific, cat. no. F-530S), with the addition of DMSO, according to the manufacturer’s protocol (the number of cycles, extension time and primer annealing temperature varied for different samples). The length of the amplicons was verified on a 1% agarose gel. Amplicons with expected length were either purified from the gel (if the presence of unspecific PCR products was revealed on the gel), or directly from the PCR mix (if there was no unspecific amplification), using Gelpure kit (NZYtech, cat. no MB01101). Amplicons were each Sanger-sequenced (Eurofins Genomics, Cologne, Germany) from both ends.

Sanger chromatograms were manually checked using FinchTV v.1.3.0 viewer for the presence of SNPs, for whether the position of each SNP on the chromatogram corresponds to that expected from the read mapping. The sequences of the genes amplified, the data on SNPs identified in amplicons, and it’s correspondence with the read mapping are available via GitHub https://github.com/beaplab/kineto_stats/blob/main/GitHub_PLOIDY/PCRs/Ploidy_PCRs_CLEAN.xlsx (sheet 2 of the table). Raw Sanger sequencing data (.ab1 files) are available via Figshare at DOI: 10.6084/m9.figshare.29088251.

### Section S8.8 Total lengths of protein-coding genes: completeness, gene count, redundancy

The protein-coding nuclear DNA content (=total length in bp) in this study is estimated based on proteomes predicted from *de novo* transcriptome assemblies. Some proteins might be either missing or over-represented in *de novo* assemblies, which may result in under- or overestimation of the protein-coding DNA content. Assembly “completeness”, distributions of protein lengths and sequence similarity within assemblies were accurately inspected, using control datasets from other protist species, to identify what affects the final estimate of protein-coding DNA content.

Benchmarking Universal Single-Copy Orthologs (BUSCO) Manni et al. (2021) is a frequently used measure to assess the completeness of assemblies and proteomes. However, for protist sequencing data, BUSCO is rarely higher than 90%, due to limited taxonomic representation of protists in reference databases from which BUSCO gene sets are formed Schoenle et al. (2025). Even though a set of BUSCO genes putatively specific to euglenozoa is available, 29 out of 31 species included in the Euglenozoa (v.10) BUSCO dataset belong to trypanosomatids - a monophyletic lineage of kinetoplastids, represented by parasitic species. On the other side, the “universal” eukaryotic BUSCO dataset (v.10) does not include a single representative of Discoba. As a result, BUSCO datasets that can be used to assess the completeness of predicted proteomes of free-living bodonids from this study, “Eugleonozoa_odb10” set and “Eukaryota_odb10” are both expected to underestimate the completeness.

Bias in BUSCO datasets is one of the possible artefact sources in the completeness estimation. Data processing during assembly and/or protein prediction steps can also introduce artefacts and biases towards the completeness estimate. Finally, the source of data itself would matter: coverage depth, sequencing technology, etc. can directly affect the outcome of data processing.

To identify whether the pipeline for *de novo* assembly and protein prediction of transcriptomes from free-living kinetoplastids used in this study affects BUSCO completeness and/or total proteome length estimate, several species from both Kinetoplastea, and other protist lineages with available well-annotated reference genomes, were selected (see https://github.com/beaplab/kineto_stats/blob/main/BUSCO_TITRATION_CLEAN_NOV2025/assemblies_proteomes_length_completeness_LEGEND.xlsx for the species list). Publicly available RNA-Seq reads from these species have been assembled *de novo*, proteins were predicted, and redundancy reduction was performed. The process was identical to that used for the species described in Section S7.7, and, apart from the decontamination step, similar to that used for obtaining proteomes of free-living kinetoplastid isolates from this study (described in detail in Zavadska et al. (2025)). See Section S8.8.3.

The set of *de novo*-reference pairs of assemblies/proteomes was expanded by adding reference assemblies/proteomes from Discoba representatives (used in Zavadska et al. (2025)) and a few non-Discoba protists. This extended set was used to assess the overall performance of BUSCO for diverse protists, and the effect of taxa included in BUSCO database on the outcome. Proteomes in the set were obtained from phylogenetically diverse taxa, different data types (genomic, transcriptomic, genomic with transcriptomic evidence), from both model organisms for which transfection tools are available, and understudied organisms which have been cultured and described only recently. Importantly, the dataset contained some of the organisms which were used in the BUSCO reference database. See Section S8.8.2.

We also used the expanded set of proteomes/assemblies to compare the effect of redundancy reduction on the proteome lengths, duplicate frequency, and BUSCO completeness. Finally, we inspected how BUSCO completeness estimate changes depending on the type of dataset used: nucleotide vs amino acid, redundancy-reduced vs original. The redundancy reduction was performed on proteomes, with CD-HIT v4.8.1 Li and Godzik (2006); Fu et al. (2012); proteins <90% amino acid global sequence identity were removed. For each proteome in the dataset, both “ORIGINAL” and redundancy-reduced (“NR90”) versions have been inspected for BUSCO completeness and other metrics. See Section S8.8.1. For simplicity of display, only the estimates for redundancy-reduced proteomes are compared in Section S8.8.3 and Section S8.8.2.

Note that for the procedures described, BUSCO v5.2.2 and BUSCO databases v.10 have been used. Scripts used to produce data in this chapter are available via Github https://github.com/beaplab/kineto_stats/tree/main/BUSCO_TITRATION_CLEAN_NOV2025, and intermediate datasets are either available via Github and FigShare at DOI: 10.6084/m9.figshare.29088251 or can be (re)produced by running the provided scripts.

#### Section S8.8.1 How does redundancy reduction affect completeness and duplication rate?

Redundancy reduction of proteomes has no effect on BUSCO completeness, when compared to the “ORIGINAL”, non-redundancy reduced proteomes Figure S9. At the same time, redundancy reduction decreases the fraction of duplicated BUSCO genes Figure S9.

Using nucleotide transcripts vs predicted proteins in the majority of cases does affect completeness or the number of duplicates. For the Euglenozoa dataset, an overall trend would be a slight increase in completeness of proteomes compared to transcripts. There are certain exceptions, however, in which the completeness of the proteome is notably lower than that of nucleotide assemblyFigure S9. This likely represents an issue with the protein prediction step. As long as predicted proteomes from the test Kinetoplastid species that we use in this study do not exhibit such a drop in completeness after protein prediction, this issue (observed in *Azumiobodo hoyamushi, Bodo saltans, Cruzella marina* and *Cryptobia borreli* data) is none of our concern. Based on the mentioned observations, we conclude that the **most optimal dataset for completeness and length assessment would be a proteome that undegone redundancy reduction at a 90% aa identity threshold.**

**Figure S9:**
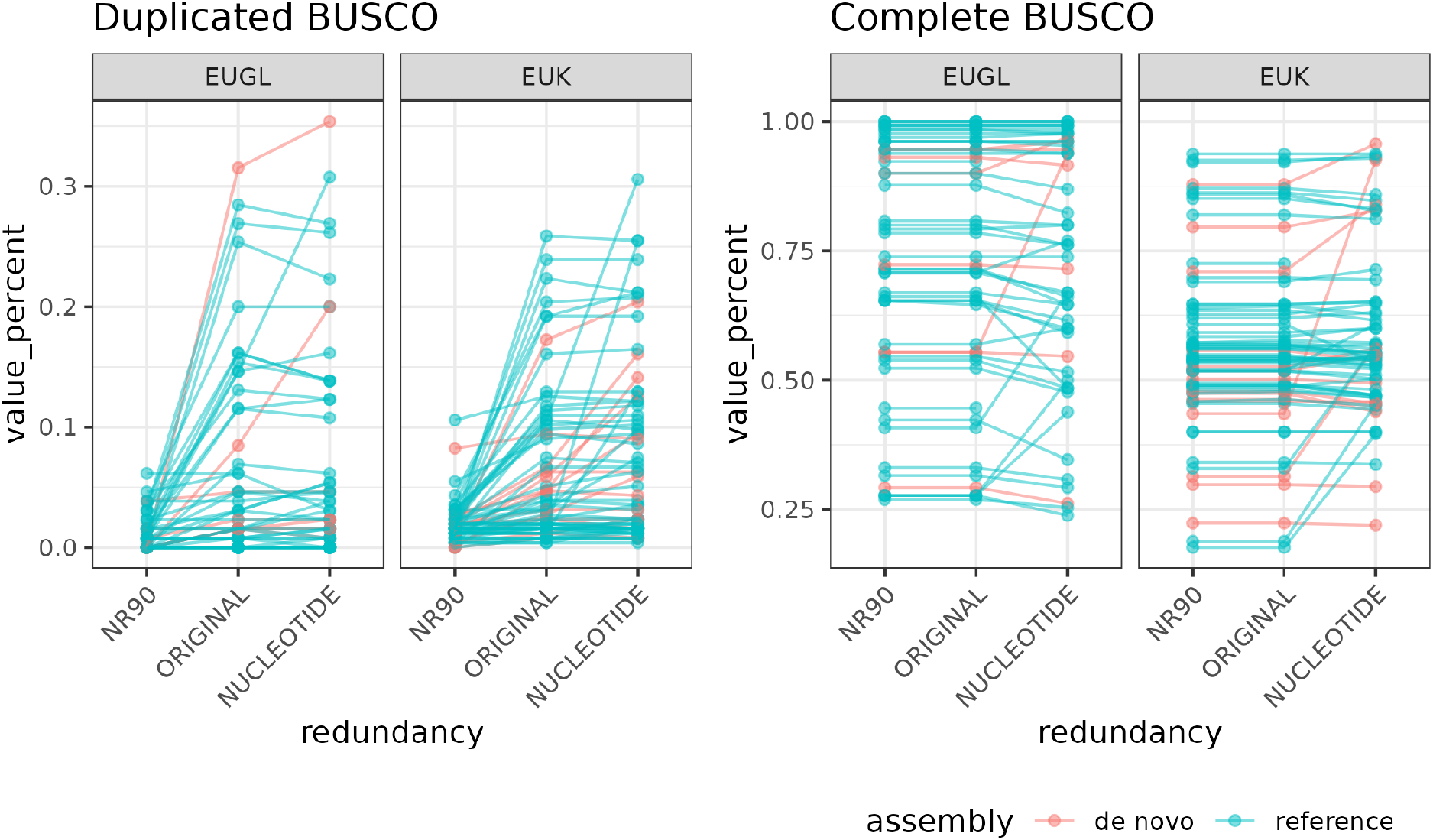
Effect of redundancy reduction and data type on the number of BUSCO duplicates and Complete BUSCO genes. Each dot represents a proteome/assembly from one species under one setting, and its position on the y-axis illustrates the percent of BUSCO genes (“Duplicated” or “Complete”). Dots from the same species are linked by lines. Colour represents whether the assembly/proteome is a part of *de novo* assembled/predicted set, used for a pipeline test in Section S8.8.3, or “reference” set. Each subplot represents BUSCO database used: “EUK” - Eukaryota BUSCO dataset; “EUGL” - Euglenozoa BUSCO dataset. The x-axis represent the combination of redundancy reduction setting and data type assessed: “ORIGINAL” - proteome with no redundancy reduction applied; “NR90” - proteome after discarding duplicates at CD-hit at 90% aa identity threshold; “NUCLEOTIDE” - nucleotide transcripts (for a few species, only CDSs were available).

#### Section S8.8.2 Is BUSCO completeness score a good indicator of assembly quality for diverse protist taxa?

Proteins that should be putatively present as a single-copy, conserved orthologs in a majority of representatives of a given taxonomic group form the BUSCO database. Yet, such proteins are identified based on the data from already sequenced and annotated species of a given taxonomic group. Unfortunately, the selection of species for forming BUSCO database is biased, in euglenozoa dataset, towards Trypanosomatids, and in the eukaryota dataset, towards multicellular organisms (Embryophyta, Metazoa and Eumycota).

The comparison of BUSCO scores for a broad selection of datasets from diverse species illustrates that the species included in the BUSCO databases demonstrate higher completeness than that of other representatives, even within the same clade Figure S10. All protists, exempt from those included in the eukaryota database, a Holozoan, and, for some reason, Heteroloboseans, have BUSCO completeness ranging between 50% and 75% for the “eukaryota” dataset Figure S10. This indicates that the “eukaryota” BUSCO dataset is likely not specific enough to accurately assess completeness of protist proteomes. “Euglenozoa” BUSCO dataset is, in contrast, based almost entirely on Trypanosomatid data. Expectedly, the majority of Trypanosomatid representatives have BUSCO scores close to 100%, whilst all representatives of all other major Euglenozoa lineages (Diplonemids, other Kinetoplastids and Euglenids) have “Euglenozoa” scores close to that of “Eukaryota” scores Figure S10. **Current BUSCO databases are bad indicators of protist proteome completeness - in particular, those of the free-living metakinetoplastids (“Bodonids”), on which this study is focused. Most likely, this is a result of biased taxon sampling used for database formation.** This highlights the importance of balanced taxon sampling when forming “universal” ortholog databases; accurate benchmarking of completeness assessment approaches for specificity/universality thresholds, and finally, correct indication of the lineages to which the given dataset should be applied. Due to the lack of viable alternatives to BUSCO completeness, we attempted to develop the Metakinetoplastid-specific ortholog set: see Section S8.8.4.

**Figure S10:**
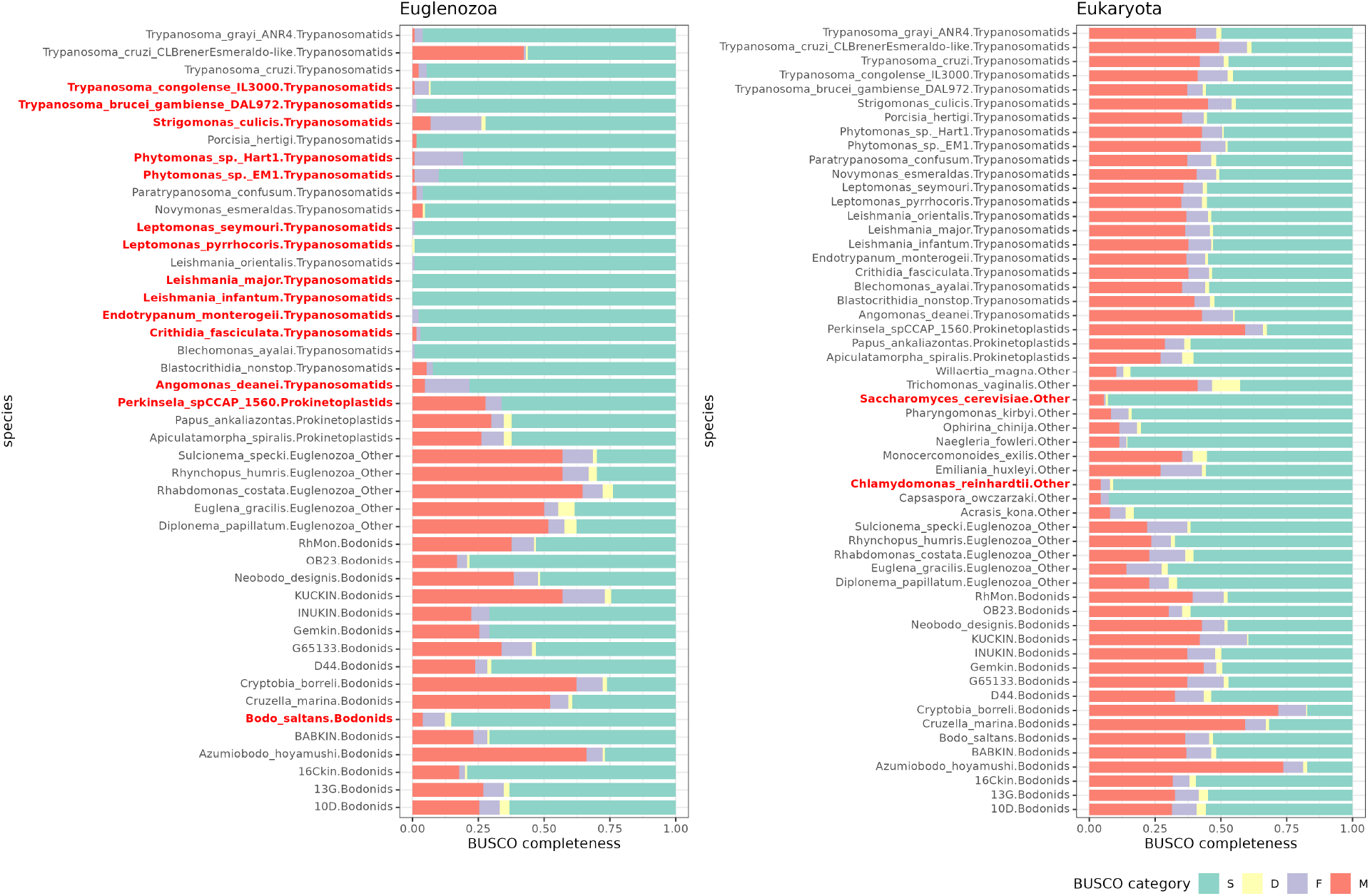
BUSCO completeness of proteomes from diverse protists. Facets represent completeness according to “Eukaryota” and “Euglenozoa” BUSCO datasets. The x-axis represents the fraction of each BUSCO gene category. Colours of the bars indicate fractions of: “S” - Complete and single-copy; “D” - Complete and duplicated; “F” - Fragmented; “M” - Missing BUSCO genes. On the y-axis, species and their taxonomic affiliation are indicated. Species which are included in “Eukaryota” or “Euglenozoa” BUSCO databases are highlighted in red.

#### Section S8.8.3 How complete and long are *de novo* assemblies/predictions?

BUSCO of the proteomes obtained *de novo* as described above was compared to that of proteomes based on the reference genomes of the same strain of the same species (the latter were obtained from either NCBI RefSeq database or TriTryp database) Figure S11-A.

Both total length of predicted proteomes and fraction of complete BUSCO genes was also compared between the *de novo* and reference proteomes; expressed on Figure S11-C, D as (*lengths_ref_ − lengths_denovo_*)*/lengths_ref_* and (*BUSCOcomplete_denovo_ − BUSCOcomplete_ref_*)*/BUSCOtotal* correspondingly. Technically, there is some correspondence between the BUSCO completeness estimated and total lengths of the proteome Figure S11-C: *de novo* proteomes with completeness lower than that of the reference from the same species tend to be shorter than reference in lengths. This indicates BUSCO score could be used to compare different assemblies/proteomes of the same species.

However, the relationship between the absolute (eukaryotic or Euglenozoan) BUSCO completness values of *de novo* proteomes and the difference in lengths between the *de novo* and reference proteomes is not as evident, especially for “eukaryota” database Figure S11-D. For “Euglenozoa” database, species with BUSCO scores *>*70% show deviation from the reference proteome lengths of only 10%. The two species with BUSCO scores *<*60% show deviation from the reference proteome lengths of *>*40%, which is a large fraction of the proteome. Theoretically, the dependence of the “euglenozoa” BUSCO score and the reduction in *de novo* assembled proteome lengths, illustrated in Figure S11-D, could be used to infer the coefficient for correcting the final estimate of proteome lenghts by “euglenozoa” BUSCO score. Using such a correction for proteomes of the test Kinetoplastid species in this study would be irrelevant though, because: (i) the datapoints for “euglenozoa” linear model as in Figure S11-D are very limited, therefore, (ii) standard errors of the fit are high, (iii) increasing the number of Euglenozoa species is not expected to improve the accuracy, as well-curated reference dataset are mainly available only for Trypanosomatids and (iv) BUSCO completeness is not the most reliable estimate when applied to non-Trypanosomatid Euglenozoa (see the section above). However, we did perform additional inference of our parameter of interest, the fraction of non-protein coding DNA content, using the BUSCO-corrected protein-coding DNA content estimate Figure S1. The protein-coding DNA content length of free-living kinetoplastids from this study was corrected according to the BUSCO completeness scores, using the parameters of linear fit from Figure S11-D for Euglenozoa dataset (slope = −0.83, intercept = 0.749). The inverse relationship between the proportions of kDNA content and non-protein coding nuclear DNA content is retained when BUSCO-corrected estimates are used Figure S1; although we note that these estimates are of limited precision due to the reasons listed above.

In the majority of cases, both BUSCO completeness and total proteome length of *de novo* assemblies are lower than that of the reference ones. However, the extent of difference varies greatly from species to species. This indicates that our assembly/protein prediction algorithm might introduce biases towards length estimation but these biases are not systematic, as they are substantial and evident only in several datasets.

Based on this conclusion, instead of implementing the completness-based correction for the total lengths, we inspected the putative sources of evident biases in *de novo* dataset lengths/completeness estimates.

Differences in BUSCO completeness and total proteome lengths between *de novo* and reference assemblies in principle has two main explanations: a lower total number of proteins, or a larger number of shorter proteins in *de novo* proteomes. The first case might be a consequence of lower read coverage; the second can be due to partial RNA degradation during extraction/library preparation. **Both reduction in average protein lengths and lower number of predicted proteins can also be an artefact of the assembly and annotation pipeline. However, in such a case, the same pipeline is expected to produce similar bias pattern for different input data, which was not the case for our pipeline.**

We investigated the distribution of protein and transcript lengths for species in our dataset Figure S12. *Diplonema papillatum*, *Bodo saltans* and *Leishmania infantum* are those euglenozoans which show the biggest reduction in both lengths and completeness of *de novo* proteomes when compared to the reference ones. *Leishmania infantum* has in total less contigs in *de novo* assembly, and less predicted proteins Figure S12. *De novo* assembly of *Diplonema papillatum* has shorter contigs than the reference one, and therefore, the length of predicted proteins is, on average, lower than that of the reference Figure S12. Finally, *Bodo saltans*, a close relative to the test species from our study with a good reference annotation publicly available, has lenghts distribution patterns that are hard to interpret. The number of long assembled contigs is **higher** in *de novo* assembly compared to the reference Figure S12. There are a lot of contigs with length *<*300bp in *de novo* assembly as well; but these contigs are filtered out in the protein prediction step (TransDecoder discards proteins *<*100aa). After the protein prediction step, the number of long predicted proteins in *de novo* predicted proteome is **less** than in the reference, while the number of short proteins is much higher in *de novo* proteome. These observations are hard to explain by solely the artefacts of data analysis performed here, and we assume that *Bodo saltans* case represents a combination of biases that could emerge from RNA extraction, library preparation, sequencing technology, gene expression patterns in the species and the annotation methods used to produce the reference.

**Overall, proteomes assembled/predicted *de novo* by the pipeline from this study have a slight overrepresentation of short protein sequences, when compared to references, or some proteomes produced in other studies Figure S12. However, the overall number of relatively long proteins is not systematically affected by the assembly pipeline. The increased fraction and number of short proteins can be a cue to an underestimation of proteome lengths.** Species analysed in this study do have some amount of shorter proteins, but the peaks in their number are not nearly as evident as those of *Diplonema papillatum* or *Bodo saltans* (KUCKIN is an exception, but the sequencing depth for this species was low, and this transcriptome is expected to be less complete than the others). We further investigate the origin of short proteins in the data from our test species, see the following section.

**Figure S11:**
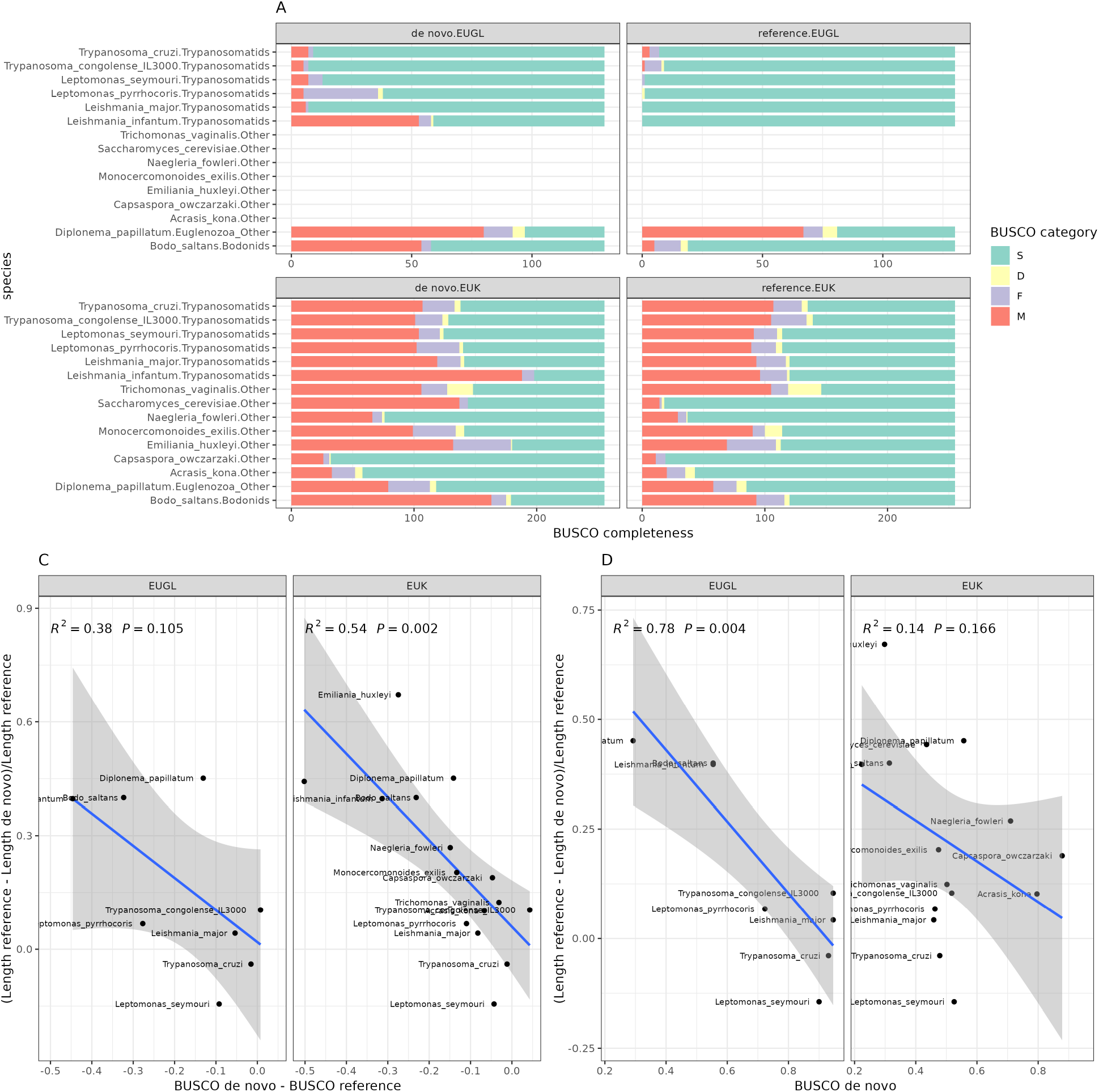
A - BUSCO completeness of *de novo* predicted proteomes and reference proteomes of the same species. “EUK” - Eukaryota BUSCO dataset; “EUGL” - Euglenozoa BUSCO dataset. Colours of the bars indicate fractions of: “S” - Complete and single-copy; “D” - Complete and duplicated; “F” - Fragmented; “M” - Missing BUSCOs. **C - Difference in length between *de novo* assembled proteome and the reference proteome, expressed as a fraction of reference proteome length (x-axis) vs difference in BUSCO completeness between *de novo* assembled proteome and the reference proteome (y-axis). D - Difference in length between *de novo* assembled proteome and the reference proteome, expressed as a fraction of reference proteome length (x-axis) vs difference in BUSCO completeness of the *de novo* assembled proteome (y-axis).** Linear regression fits on C and D were performed using the “geom_smooth” function from the ggplot2 R package, using “lm” method from stats R package for the fit.

**Figure S12:**
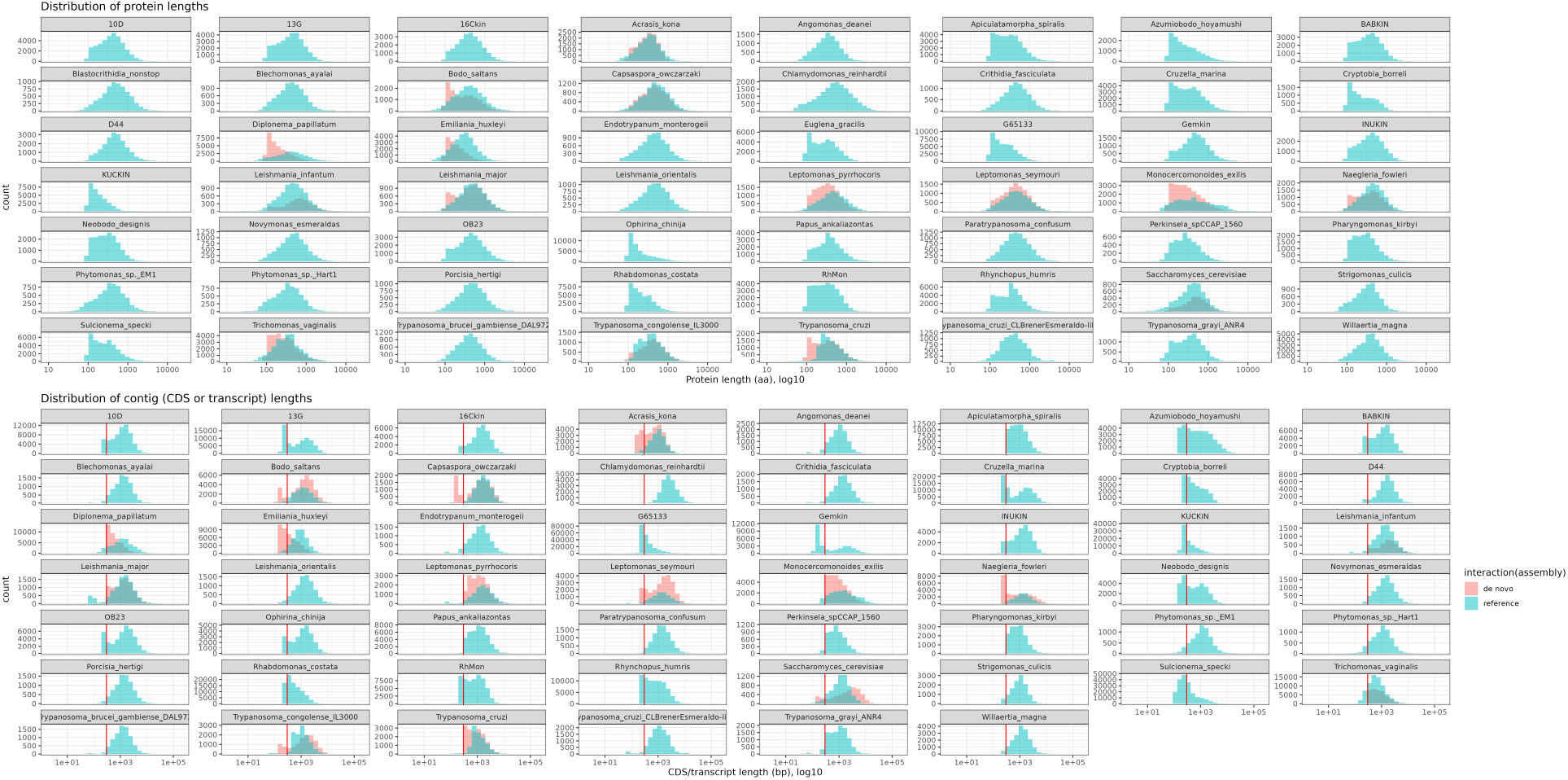
Length distribution of predicted proteins and contigs. Each subplot represents one species from the dataset; the x-axis represents lengths of proteins or contigs (note the log-10 scale). For protein length distribution, redundancy-reduced proteomes were used. The red vertical line on the contig lengths plot indicates the lengths of 300bp: proteins below 100aa are discarded at the protein annotation step; therefore, proteins from contigs shorter than 300bp are not included in the predicted proteome.

To summarise, based on the comparisons presented above:

- Redundancy reduction of proteomes decreases the number of duplicates without reducing the number of complete BUSCO genes; therefore, for proteome lengths estimation, redundancy-reduced proteomes should be used.
- BUSCO completeness score is highly biased due to poor taxon sampling: the “Eukaryota” BUSCO database scores underestimate completeness of the majority of protist proteomes, while the “Euglenozoa” database likely overestimates the completeness of Trypanosomatid proteomes, underestimating the rest of Euglenozoa. This means that BUSCO completeness score is not reliable when applied towards the proteome of free-living Metakinetoplastids investigated here; alternatives should be developed.
- *De novo* assembly and protein prediction pipeline used in our study may systematically produce big amount of short proteins, which are not detected in the reference “control” proteomes. However, the overall number of *de novo* assembled/predicted proteins, proteome total length and completeness, and the difference of these metrics from the reference proteomes, vary greatly depending on the species. This indicates that there is no systematic error in our data analysis pipeline, and the biases likely emerge earlier in the process of sequencing/sample preparation.

#### Section S8.8.4 *De novo* inferred Metakinetoplastina-specific orthogroups as an alternative completeness estimate

To create a set of Metakinetoplastina-specific OGs, a set of redundancy-reduced proteomes from Metakinetoplastina representatives used in Zavadska et al. (2025) was taken, and Orthogroups (OGs) were inferred *de novo* with OrthoFinder v2.5.5 software Emms and Kelly (2019), (“diamond_ultra_sens” option for the sequence search algorithm; “msa” option for gene tree inference; other options left as their defaults). This set of OGs was further reduced to contain only the OGs which would have genes present across the majority of Metakinetoplastid species, predominantly as single-copy genes (similar to the criteria used in BUSCO datasets). The reduced set of Metakinetoplastid-maybe/more-universal, single-copy orthologs (M3USCO) could serve as an alternative indicator of proteome completeness - since BUSCO datasets are not well-suited for this task, as was shown above.

To form M3USCO gene set, we selected OGs which would satisfy all the criteria: (i) mean number of proteins per single species between 0.9 and 1, inclusive, (ii) standard deviation of the number of proteins per single species less than or equal to 0.5, AND (iii) number of species with at least one protein present greater than 30 (out of 36 species in total). This approach has already been used in Zavadska et al. (2025). The distributions of the mean and standard deviation of the proteins per OG, along with the distribution of species representation per OG were checked before setting the threshold values Figure S13.

The resulting M3USCO set contained genes from 1413 OGs. Figure S14 shows the copy number and absence/presence of the genes in the M3USCO set across the proteomes of various Metakinetoplastids. **If we consider M3USCO as an alternative completeness metric, it clearly indicates several low-completeness proteomes (*Cruzella marina, Cryptobia borreli, Azumiobodo hoyamushi*), while others, including the ones from the isolates used in our study, are almost complete Figure S14. This further proves that there is no need for correcting proteome lengths based only on BUSCO completeness.** Yet, several proteomes of species from this study have slightly elevated numbers of M3USCO duplicates Figure S14; in addition, these proteomes have elevated numbers of short proteins Figure S12, when compared to others. This could point to a potential artefact in our dataset. Therefore, we attempted to establish the nature of short and duplicated proteins in our dataset based on *de novo* inferred OGs.

**Figure S13:**
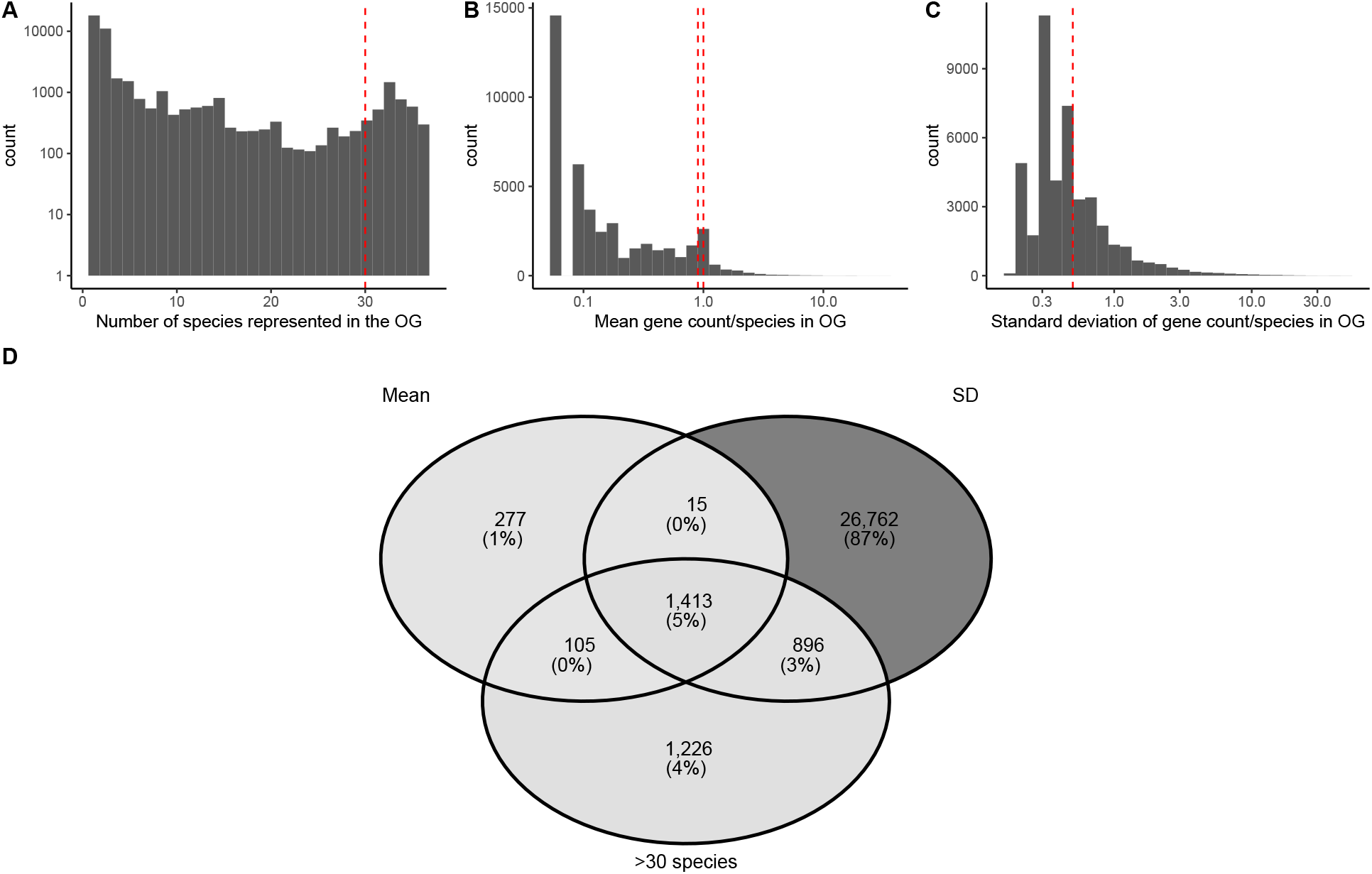
Subsetting OGs. **A-C**: distributions of parameters used to establish thresholds to subset OGs. The count of OGs with a given parameter value is represented on the vertical axis, and the values of the three different parameters are represented on the horizontal axis. The dashed red vertical lines in A-C represent thresholds used for subsetting OGs. Note log10 scales in A-C. **D**: the count of OGs with the mean number of proteins per single species between or equal to 1 and 0.9 (“Mean”), standard deviation of number of proteins per single species lower than or equal to 0.5 (“SD”), and the number of species with at least one protein in a given OG higher than 30 (“>30 species”).

#### Section S8.8.5 Short proteins in proteomes

High numbers of short proteins could affect the final estimate of total proteome length. Short proteins could either be artefacts of protein prediction software, which do not represent real sequences translated in the organism, or fragments of longer protein-coding genes, which do represent parts of “real” sequences. In the latter case, the sum lengths of shorter proteins should equal the lengths of a single longer protein, and protein fragmentation does not affect the total length of the proteome. The overrepresentation of short sequences which do not correspond to “real” proteins could lead to an overestimation of total proteome lengths. “Real” short protein sequences can be distinguished from artefactual ones by the presence of homology to the proteins of other related species (though some genes and proteins can appear spontaneously from the non-coding ancestral sequence, the majority of protein-coding genes still evolve from ancestral protein-coding sequences). In such case, the “real” proteins are likely to be included into OG with sequences from more than one species, when classified by OrthoFinder. The artefactual proteins, in contrast, would not have homologs in the related species, and would be assigned to “singletons”, or “species-specific OGs” - OGs consisting entirely of genes from one species. Alternatively, artefactual proteins would end up not being assigned to any OG.

**Figure S14:**
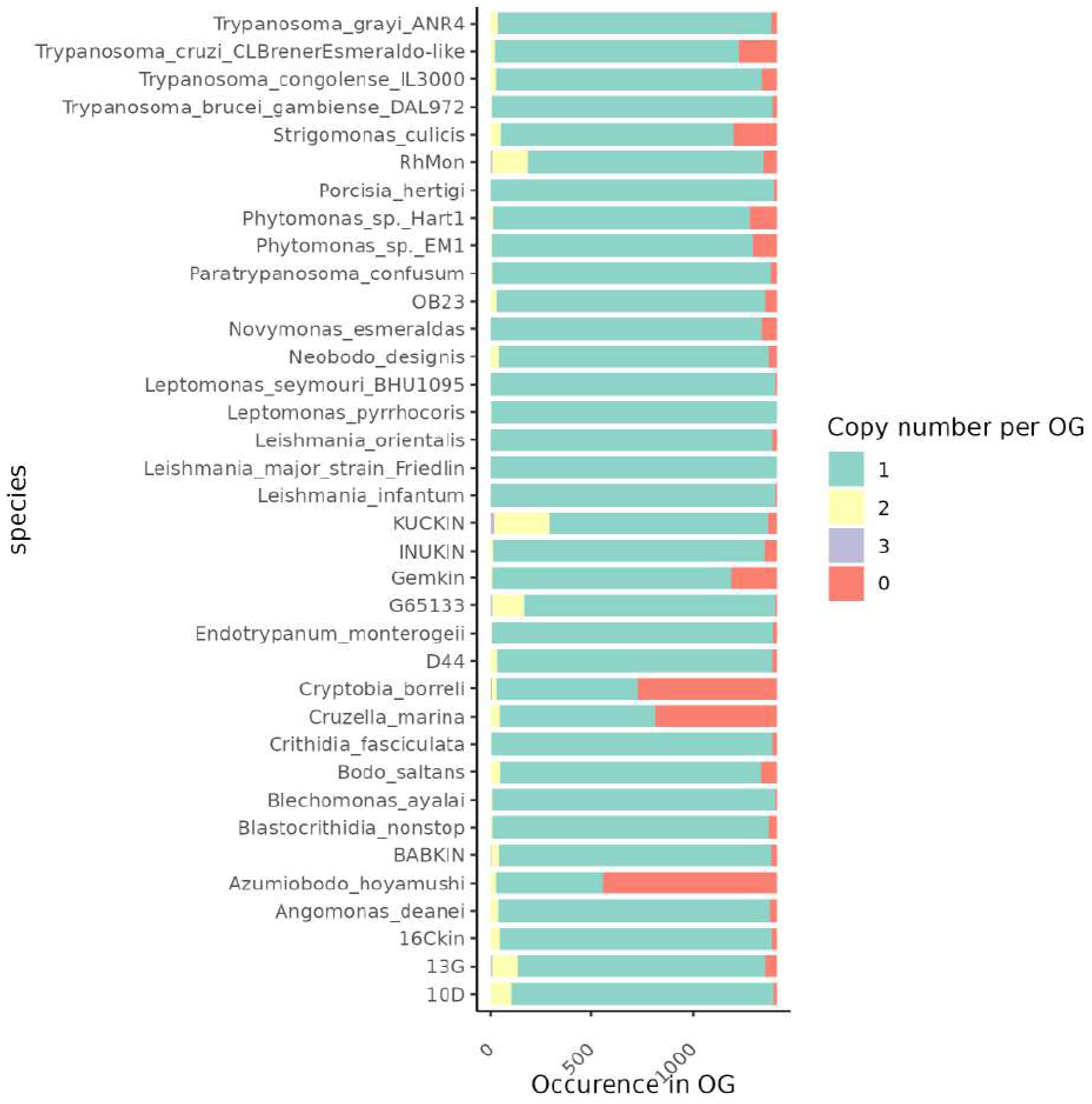
Representation of genes from M3USCO OG set across Metakinetoplaastid proteomes.

In the Metakinetoplastid OG dataset, described above, we analysed the total lengths of proteins shorter than 200aa, which belong to either singleton OGs or do not belong to any OG Figure S15. Among the proteomes used in this study, three species: G65133, RhMon and KUCKIN have a total length of short (*<*200 aa) proteins contributing more than 10% towards the total length of all the proteins Figure S15. As mentioned above, KUCKIN initially had lower read coverage than other species from the study, and its proteome length is not considered in the final estimations. For G65133 and RhMon, at least half of the total length of short proteins can be attributed to proteins in singletons, or not assigned to any OG: the “artefactual” proteins, without a homology to any sequences in related species Figure S15. In G65133, only 15% of the total non-redundant proteome lengths could be attributed towards the sequences which are most likely artefacts of data analysis and do not represent the underlying biological entities. For other species used in this study, the fraction of such artefacts is negligible, in terms of their lengths relative to the whole proteome.

**Figure S15:**
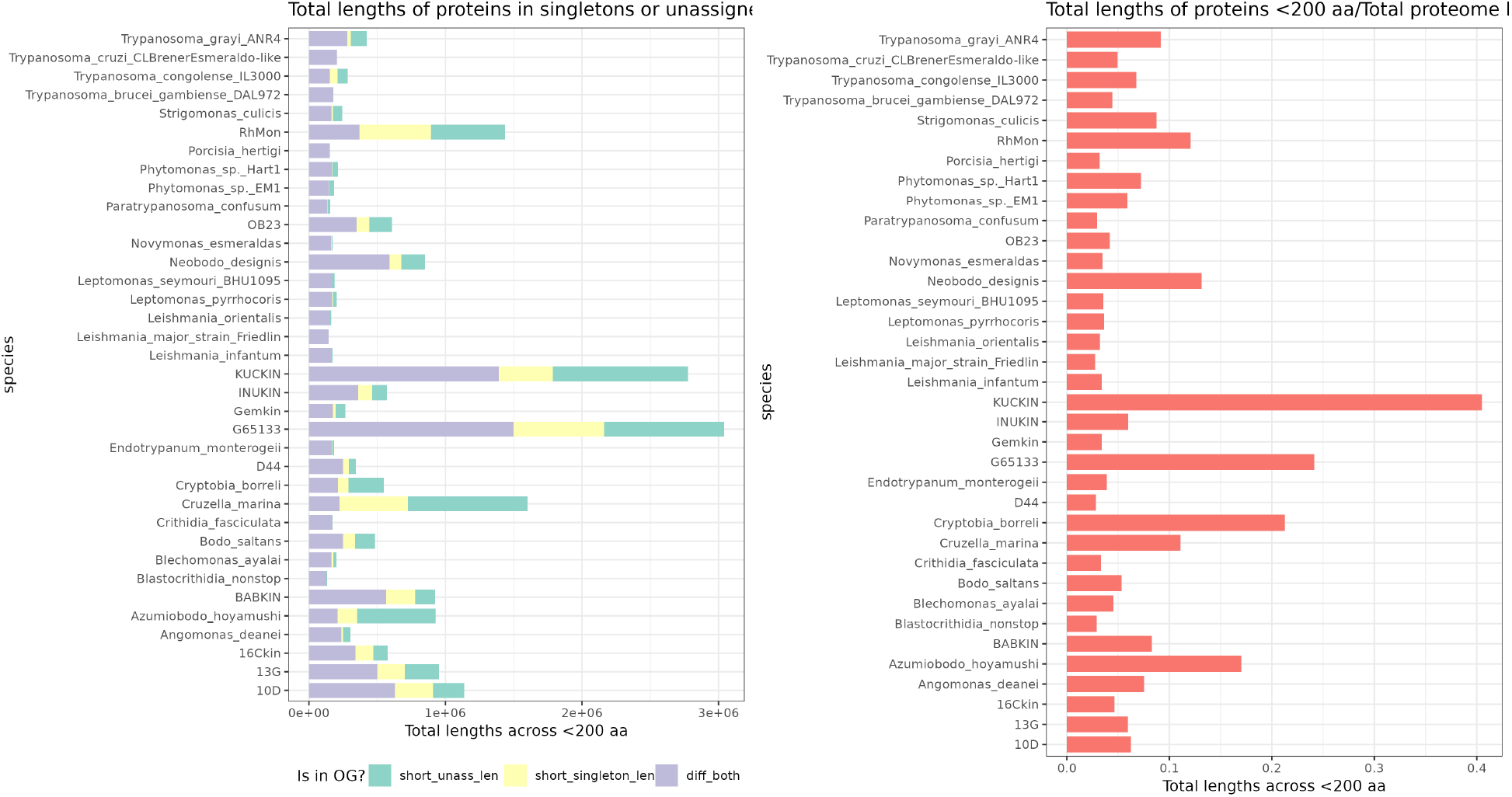
The total aa length of short (*<*200 aa) proteins, their OG affiliation, and the fraction of total proteome length constituted by the short proteins. The colours on the first subplot represent: “short_unass_len” - total length of short proteins which are not assigned to any OG; “short_singleton_len” - total length of short proteins which are assigned to sigletons; “diff_both” - total length of short proteins which are assigned to an OG with proteins from more than a single species.

## References

Adl, S. M., Bass, D., Lane, C. E., Lukeš, J., Schoch, C. L., Smirnov, A., Agatha, S., Berney, C., Brown, M. W., Burki, F., et al. (2019). Revisions to the classification, nomenclature, and diversity of eukaryotes. Journal of Eukaryotic Microbiology, 66(1):4–119.

Akiyoshi, B. and Gull, K. (2014). Discovery of unconventional kinetochores in kinetoplastids. Cell, 156(6):1247–1258.

Amodeo, S., Bregy, I., and Ochsenreiter, T. (2023). Mitochondrial genome maintenancethe kinetoplast story. FEMS Microbiology Reviews, 47(6):fuac047.

Benne, R. (1990). RNA editing in trypanosomes: is there a message? Trends in genetics, 6:177–181.

Bhattacharjee, A. and Biswas, A. (2023). Cycling within a cell: Cell cycle of intracellular kinetoplastid parasites. In Pathobiology of Parasitic Protozoa: Dynamics and Dimensions, pages 95–107. Springer.

Bischoff, H. W. Bold, H. C. (1963). Some soil algae from enchanted rock and related algal species. Phycological Studies, IV.

Blommaert, J. (2020). Genome size evolution: towards new model systems for old questions. Proceedings of the Royal Society B: Biological Sciences, 287(1933).

Borst, P. and Hoeijmakers, J. H. (1979). Kinetoplast DNA. Plasmid, 2(1):20–40.

Borst, P. and Sabatini, R. (2008). Base J: discovery, biosynthesis, and possible functions. Annu. Rev. Microbiol., 62(1):235–251.

Broemsen, E. L., Place, A. R., and Parrow, M. W. (2021). Quantitative nuclear dna content and cell cycle analysis of a mixotrophic dinoflagellate by image cytometry. Limnology and Oceanography: Methods, 19(4):253–266.

Bushmanova, E., Antipov, D., Lapidus, A., and Prjibelski, A. D. (2019). rnaspades: a de novo transcriptome assembler and its application to rna-seq data. GigaScience, 8(9):giz100.

Butenko, A., Opperdoes, F. R., Flegontova, O., Horák, A., Hampl, V., Keeling, P., Gawryluk, R. M., Tikhonenkov, D., Flegontov, P., and Lukeš, J. (2020). Evolution of metabolic capabilities and molecular features of diplonemids, kinetoplastids, and euglenids. BMC biology, 18(1):23.

Cachon, J., Cachon, M., Cosson, M.-P., and Cosson, J. (1988). The paraflagellar rod: a structure in search of a function. Biology of the Cell, 63(2):169–181.

Campbell, D. A., Thomas, S., and Sturm, N. R. (2003). Transcription in kinetoplastid protozoa: why be normal? Microbes and infection, 5(13):1231–1240.

Cavalier-Smith, T. (1980). R-and K-tactics in the evolution of protist developmental systems: cell and genome size, phenotype diversifying selection, and cell cycle patterns. Biosystems, 12(1-2):43–59.

Cavalier-Smith, T. (2005). Economy, speed and size matter: evolutionary forces driving nuclear genome miniaturization and expansion. Annals of botany, 95(1):147–175.

Cavalier-Smith, T. and Beaton, M. (1999). The skeletal function of non-genic nuclear dna: new evidence from ancient cell chimaeras. Genetica, 106(1):3–13.

Čertnerová, D., Škaloud, P., Jadrná, I., and Čertner, M. (2025). Large genomes are associated with greater cell size and ecological shift towards more nitrogen-rich and higher-latitude environments in microalgae of the genus *Synura*. Journal of Eukaryotic Microbiology, 72(4):e70026.

Chen, S., Zhou, Y., Chen, Y., and Gu, J. (2018). Fastp: an ultra-fast all-in-one fastq preprocessor. Bioinformatics, 34(17):i884–i890.

Clayton, C. (2016). Gene expression in kinetoplastids. Current opinion in microbiology, 32:46–51.

Cosgrove, W. B. and Skeen, M. J. (1970). The cell cycle in crithidia fasciculata. temporal relationships between synthesis of deoxyribonucleic acid in the nucleus and in the kinetoplast. The Journal of Protozoology, 17(2):172–177.

Covello, P. and Gray, M. (1993). On the evolution of RNA editing. Trends in Genetics, 9(8):265–268.

Danecek, P., Bonfield, J. K., Liddle, J., Marshall, J., Ohan, V., Pollard, M. O., Whitwham, A., Keane, T., McCarthy, S. A., Davies, R. M., et al. (2021). Twelve years of samtools and bcftools. Gigascience, 10(2):giab008.

Derelle, E., Ferraz, C., Rombauts, S., Rouzé, P., Worden, A. Z., Robbens, S., Partensky, F., Degroeve, S., Echeynié, S., Cooke, R., et al. (2006). Genome analysis of the smallest free-living eukaryote *Ostreococcus tauri* unveils many unique features. Proceedings of the National Academy of Sciences, 103(31):11647–11652.

Donelson, J. E., Gardner, M. J., and El-Sayed, N. M. (1999). More surprises from Kinetoplastida. Proceedings of the National Academy of Sciences, 96(6):2579–2581.

Elliott, T. A. and Gregory, T. R. (2015). What’s in a genome? the C-value enigma and the evolution of eukaryotic genome content. Philosophical Transactions of the Royal Society B: Biological Sciences, 370(1678):20140331.

Emms, D. M. and Kelly, S. (2019). Orthofinder: phylogenetic orthology inference for comparative genomics. Genome biology, 20(1):238.

Erken, M., Farrenschon, N., Speckmann, S., Arndt, H., and Weitere, M. (2012). Quantification of individual flagellate-bacteria interactions within semi-natural biofilms. Protist, 163(4):632–642.

Erken, M., Weitere, M., Kjelleberg, S., and McDougald, D. (2011). *In situ* grazing resistance of vibrio cholerae in the marine environment. FEMS microbiology ecology, 76(3):504–512.

Flegontov, P., Gray, M. W., Burger, G., and Lukeš, J. (2011). Gene fragmentation: a key to mitochondrial genome evolution in Euglenozoa? Current genetics, 57(4):225–232.

Fu, L., Niu, B., Zhu, Z., Wu, S., and Li, W. (2012). Cd-hit: accelerated for clustering the next-generation sequencing data. Bioinformatics, 28(23):3150–3152.

Gibson, W. (2017). Kinetoplastea. In Archibald, J. M., Simpson, A. G. B., and Slamovits, C. H., editors, Handbook of the Protists, pages 1–50. Springer International Publishing, Cham, 2 edition.

Gigeroff, A. S., Eglit, Y., and Simpson, A. G. (2023). Characterisation and cultivation of new lineages of colponemids, a critical assemblage for inferring alveolate evolution. Protist, 174(2):125949.

Gray, M. W., Lukeš, J., Archibald, J. M., Keeling, P. J., and Doolittle, W. F. (2010). Irremediable complexity? Science, 330(6006):920–921.

Gregory, T. R. (2001). Coincidence, coevolution, or causation? DNA content, cell size, and the C-value enigma. Biological reviews, 76(1):65–101.

Hajduk, S. and Ochsenreiter, T. (2010). RNA editing in kinetoplastids. RNA biology, 7(2):229–236.

Hannaert, V., Bringaud, F., Opperdoes, F. R., and Michels, P. A. (2003). Evolution of energy metabolism and its compartmentation in Kinetoplastida. Kinetoplastid biology and disease, 2(1):11.

Hannaert, V. and Michels, P. A. (1994). Structure, function, and biogenesis of glycosomes in Kinetoplastida. Journal of bioenergetics and biomembranes, 26(2):205–212.

Hessen, D. O., Jeyasingh, P. D., Neiman, M., and Weider, L. J. (2010). Genome streamlining and the elemental costs of growth. Trends in ecology & evolution, 25(2):75–80.

Hof, J. V. and Sparrow, A. H. (1963). A relationship between DNA content, nuclear volume, and minimum mitotic cycle time. Proceedings of the National Academy of Sciences, 49(6):897–902.

Hong, M. and Simpson, L. (2003). Genomic organization of *Trypanosoma brucei* kinetoplast DNA minicircles. Protist, 154(2):265–279.

Howell, J., Omwenga, S., Jimenez, M., and Hammarton, T. C. (2024). Analysis of the *Leishmania mexicana* promastigote cell cycle using imaging flow cytometry provides new insights into cell cycle flexibility and events of short duration. PLoS One, 19(10):e0311367.

Jeffares, D. C., Mourier, T., and Penny, D. (2006). The biology of intron gain and loss. TRENDS in Genetics, 22(1):16–22.

Katju, V. and Bergthorsson, U. (2019). Old trade, new tricks: insights into the spontaneous mutation process from the partnering of classical mutation accumulation experiments with high-throughput genomic approaches. Genome biology and evolution, 11(1):136–165.

Keeling, P. J. (2004). Reduction and compaction in the genome of the apicomplexan parasite *Cryptosporidium parvum*. Developmental Cell, 6(5):614–616.

Keeling, P. J. and Slamovits, C. H. (2005). Causes and effects of nuclear genome reduction. Current opinion in genetics & development, 15(6):601–608.

Kiørboe, T., Grossart, H.-P., Ploug, H., Tang, K., and Auer, B. (2004). Particle-associated flagellates: swimming patterns, colonization rates, and grazing on attached bacteria. Aquatic Microbial Ecology, 35(2):141–152.

Kostygov, A. Y., Albanaz, A. T., Butenko, A., Gerasimov, E. S., Lukeš, J., and Yurchenko, V. (2024a). Phylogenetic framework to explore trait evolution in Trypanosomatidae. Trends in Parasitology, 40(2):96–99.

Kostygov, A. Y., Skỳpalová, K., Kraeva, N., Kalita, E., McLeod, C., Yurchenko, V., Field, M. C., Lukeš, J., and Butenko, A. (2024b). Comprehensive analysis of the Kinetoplastea intron landscape reveals a novel intron-containing gene and the first exclusively trans-splicing eukaryote. BMC biology, 22(1):281.

Kuroiwa, T., Ohnuma, M., Nozaki, H., Imoto, Y., Misumi, O., and Kuroiwa, H. (2015). Cytological evidence of cell-nuclear genome size of a new ultra-small unicellular freshwater green alga, Medakamo hakoo strain M-hakoo 311 I. comparison with *Cyanidioschyzon merolae* and *Ostreococcus tauri*. Cytologia, 80(2):143–150.

Lai, D.-H., Hashimi, H., Lun, Z.-R., Ayala, F. J., and Lukeš, J. (2008). Adaptations of *Trypanosoma brucei* to gradual loss of kinetoplast DNA: *Trypanosoma equiperdum* and *Trypanosoma evansi* are petite mutants of *T. brucei*. Proceedings of the National Academy of Sciences, 105(6):1999–2004.

Lane, N. and Martin, W. (2010). The energetics of genome complexity. Nature, 467(7318):929–934.

Langmead, B. and Salzberg, S. L. (2012). Fast gapped-read alignment with bowtie2. Nature methods, 9(4):357–359.

Larsen, T. A., Goodsell, D. S., Cascio, D., Grzeskowiak, K., and Dickerson, R. E. (1989). The structure of DAPI bound to DNA. Journal of Biomolecular Structure and Dynamics, 7(3):477–491.

Levin, T. C. and King, N. (2013). Evidence for sex and recombination in the choanoflagellate *Salpingoeca rosetta*. Current Biology, 23(21):2176–2180.

Li, W. and Godzik, A. (2006). Cd-hit: a fast program for clustering and comparing large sets of protein or nucleotide sequences. Bioinformatics, 22(13):1658–1659.

Lin, Y.-W., Chiu, T.-C., and Chang, H.-T. (2003). Laser-induced fluorescence technique for DNA and proteins separated by capillary electrophoresis. Journal of Chromatography B, 793(1):37–48.

Liu, B., Liu, Y., Motyka, S. A., Agbo, E. E., and Englund, P. T. (2005). Fellowship of the rings: the replication of kinetoplast DNA. Trends in parasitology, 21(8):363–369.

López-Escardó, D., Obiol, A., Marimon, G., López-Alforja, X., Vaqué, D., Forn, I., Logares, R., Yau, S., Fornas, Ò., Martínez-García, M., et al. (2026). Single-cell sequencing reveals genome streamlining and functional diversity in ecologically dominant marine protists. bioRxiv, pages 2026–04.

Lukeš, J., Archibald, J. M., Keeling, P. J., Doolittle, W. F., and Gray, M. W. (2011). How a neutral evolutionary ratchet can build cellular complexity. IUBMB life, 63(7):528–537.

Lukeš, J., Wheeler, R., Jirsová, D., David, V., and Archibald, J. M. (2018). Massive mitochondrial DNA content in diplonemid and kinetoplastid protists. IUBMB life, 70(12):1267–1274.

Lukeš, J., Lys Guilbride, D., Votypka, J., Zíková, A., Benne, R., and Englund, P. T. (2002). Kinetoplast DNA network: evolution of an improbable structure. Eukaryotic cell, 1(4):495–502.

Lynch, M. (2002). Intron evolution as a population-genetic process. Proceedings of the National Academy of Sciences, 99(9):6118–6123.

Lynch, M. (2006a). The origins of eukaryotic gene structure. Molecular biology and evolution, 23(2):450–468.

Lynch, M. (2006b). Streamlining and simplification of microbial genome architecture. Annu. Rev. Microbiol., 60(1):327–349.

Lynch, M. (2025). Energetics and evolution: Response to Martin. Biochimica et Biophysica Acta (BBA)-Bioenergetics, page 149576.

Lynch, M. and Conery, J. S. (2003). The origins of genome complexity. science, 302(5649):1401–1404.

Lynch, M. and Marinov, G. K. (2015). The bioenergetic costs of a gene. Proceedings of the National Academy of Sciences, 112(51):15690–15695.

Maharana, B., Tewari, A., and Singh, V. (2015). An overview on kinetoplastid paraflagellar rod. Journal of parasitic diseases, 39(4):589–595.

Malerba, M. E., Ghedini, G., and Marshall, D. J. (2020). Genome size affects fitness in the eukaryotic alga *Dunaliella tertiolecta*. Current Biology, 30(17):3450–3456.

Manni, M., Berkeley, M. R., Seppey, M., Simão, F. A., and Zdobnov, E. M. (2021). BUSCO update: novel and streamlined workflows along with broader and deeper phylogenetic coverage for scoring of eukaryotic, prokaryotic, and viral genomes. Molecular biology and evolution, 38(10):4647–4654.

McGrath, C. L. and Katz, L. A. (2004). Genome diversity in microbial eukaryotes. Trends in ecology & evolution, 19(1):32–38.

Mensa-Wilmot, K., Hoffman, B., Wiedeman, J., Sullenberger, C., and Sharma, A. (2019). Kinetoplast division factors in a trypanosome. Trends in parasitology, 35(2):119–128.

Mukherjee, C., Majumder, S., and Lohia, A. (2009). Inter-cellular variation in DNA content of *Entamoeba histolytica* originates from temporal and spatial uncoupling of cytokinesis from the nuclear cycle. PLoS Neglected Tropical Diseases, 3(4):e409.

Munyenyembe, K., Timmons, C., Weiner, A. K., Katz, L. A., and Yan, Y. (2021). DAPI staining and DNA content estimation of nuclei in uncultivable microbial eukaryotes (arcellinida and ciliates). European journal of protistology, 81:125840.

Noirot, M., Barre, P., Louarn, J., Duperray, C., and Hamon, S. (2002). Consequences of stoichiometric error on nuclear dna content evaluation in *Coffea liberica* var. dewevrei using DAPI and propidium iodide. Annals of Botany, 89(4):385–389.

Oberle, M., Balmer, O., Brun, R., and Roditi, I. (2010). Bottlenecks and the maintenance of minor genotypes during the life cycle of *trypanosoma brucei*. PLoS pathogens, 6(7):e1001023.

Olefeld, J. L., Majda, S., Albach, D. C., Marks, S., and Boenigk, J. (2018). Genome size of chrysophytes varies with cell size and nutritional mode. Org Divers Evol, 18:163–173.

Parfrey, L. W. and Katz, L. A. (2010). Genome dynamics are influenced by food source in *Allogromia laticollaris* strain CSH (Foraminifera). Genome Biology and Evolution, 2:678–685.

Parrow, M. W. and Burkholder, J. M. (2002). Flow cytometric determination of zoospore dna content and population DNA distribution in cultured *Pfiesteria* spp.(Pyrrhophyta). Journal of experimental marine biology and ecology, 267(1):35–51.

Pérez, S. D., Grummer, J. A., Fernandes-Santos, R. C., José, C. T., Medici, E. P., and Marcili, A. (2019). Phylogenetics, patterns of genetic variation and population dynamics of *trypanosoma terrestris* support both coevolution and ecological host-fitting as processes driving trypanosome evolution. Parasites & vectors, 12(1):473.

Pérez-Posada, A., Dudin, O., Ocaña-Pallarès, E., Ruiz-Trillo, I., and Ondracka, A. (2020). Cell cycle transcriptomics of *Capsaspora* provides insights into the evolution of cyclin-CDK machinery. PLoS genetics, 16(3):e1008584.

Petrov, D. A. (2002). Mutational equilibrium model of genome size evolution. Theoretical population biology, 61(4):531–544.

Poulíèková, A., Mazalová, P., Vašut, R. J., Šarhanová, P., Neustupa, J., and Škaloud, P. (2014). DNA content variation and its significance in the evolution of the genus *Micrasterias* (Desmidiales, Streptophyta). PLoS One, 9(1):e86247.

Rajic, Z. A., Jankovic, G. M., Vidovic, A., Milic, N. M., Skoric, D., Pavlovic, M., and Lazarevic, V. (2005). Size of the protein-coding genome and rate of molecular evolution. Journal of human genetics, 50(5):217–229.

Robinson, J. T., Thorvaldsdóttir, H., Winckler, W., Guttman, M., Lander, E. S., Getz, G., and Mesirov, J. P. (2011). Integrative genomics viewer. Nature biotechnology, 29(1):24–26.

Sarkans, U., Gostev, M., Athar, A., Behrangi, E., Melnichuk, O., Ali, A., Minguet, J., Rada, J. C., Snow, C., Tikhonov, A., et al. (2018). The biostudies databaseone stop shop for all data supporting a life sciences study. Nucleic acids research, 46(D1):D1266–D1270.

Schavemaker, P. E. and Lynch, M. (2025). Bioenergetics and the evolution of cellular traits. Annual review of biophysics, 54(1):81–99.

Schavemaker, P. E. and Muñoz-Gómez, S. A. (2022). The role of mitochondrial energetics in the origin and diversification of eukaryotes. Nature ecology & evolution, 6(9):1307–1317.

Schoenle, A., Francis, O., Archibald, J. M., Burki, F., de Vries, J., Dumack, K., Eme, L., Florent, I., Hehenberger, E., Hoffmeyer, T. T., et al. (2025). Protist genomics: key to understanding eukaryotic evolution. Trends in Genetics.

Sebé-Pedrós, A., Irimia, M., Del Campo, J., Parra-Acero, H., Russ, C., Nusbaum, C., Blencowe, B. J., and Ruiz-Trillo, I. (2013). Regulated aggregative multicellularity in a close unicellular relative of metazoa. elife, 2:e01287.

Šímová, I. and Herben, T. (2012). Geometrical constraints in the scaling relationships between genome size, cell size and cell cycle length in herbaceous plants. Proceedings of the Royal Society B: Biological Sciences, 279(1730):867–875.

Simpson, A. G., Lukeš, J., and Roger, A. J. (2002). The evolutionary history of kinetoplastids and their kinetoplasts. Molecular Biology and Evolution, 19(12):2071–2083.

Simpson, L., Thiemann, O. H., Savill, N. J., Alfonzo, J. D., and Maslov, D. (2000). Evolution of RNA editing in trypanosome mitochondria. Proceedings of the National Academy of Sciences, 97(13):6986–6993.

Speijer, D. (2006). Is kinetoplastid pan-editing the result of an evolutionary balancing act? IUBMB life, 58(2):91–96.

Timmons, C., Le, K., Rappaport, H., Sterner, E. G., Maurer-Alcalá, X. X., Goldstein, S. T., and Katz, L. A. (2024). Foraminifera as a model of eukaryotic genome dynamism. Mbio, 15(3):e03379–23.

Tophøj, J., Wollenberg, R. D., Sondergaard, T. E., and Eriksen, N. T. (2018). Feeding and growth of the marine heterotrophic nanoflagellates, *Procryptobia sorokini* and *Paraphysomonas imperforata* on a bacterium, *Pseudoalteromonas* sp. with an inducible defence against grazing. PLoS One, 13(4):e0195935.

Tromer, E. C., Wemyss, T. A., Ludzia, P., Waller, R. F., and Akiyoshi, B. (2021). Repurposing of synaptonemal complex proteins for kinetochores in Kinetoplastida. Open Biology, 11(5).

Vesteg, M., Hadariová, L., Horváth, A., Estraño, C. E., Schwartzbach, S. D., and Krajčovič, J. (2019). Comparative molecular cell biology of phototrophic euglenids and parasitic trypanosomatids sheds light on the ancestor of Euglenozoa. Biological Reviews, 94(5):1701–1721.

Von Dassow, P., Petersen, T. W., Chepurnov, V. A., and Armbrust, V. E. (2008). Inter-and intraspecific relationships between nuclear DNA content and cell size in selected members of the centric diatom genus *Thalassiosira* (bacillariophyceae). Journal of Phycology, 44(2):335–349.

Wang, J.-F., Bi, Y.-X., Hammond, M., Lukeš, J., Lai, D.-H., and Lun, Z.-R. (2025). Kinetoplast DNA structure, RNA editing patterns and small respiratory Complex I in *Trypanosoma musculi*. Communications Biology, 8(1):1544.

Wheeler, R. J. (2020). Imagej for partially and fully automated analysis of trypanosome micrographs. In Trypanosomatids: Methods and Protocols, pages 385–408. Springer.

Wheeler, R. J., Gull, K., and Gluenz, E. (2012). Detailed interrogation of trypanosome cell biology via differential organelle staining and automated image analysis. BMC biology, 10(1):1.

Woodward, R. and Gull, K. (1990). Timing of nuclear and kinetoplast DNA replication and early morphological events in the cell cycle of *Trypanosoma brucei*. Journal of cell science, 95(1):49–57.

Zavadska, D., Packer, J. A., Simpson, A., and Richter, D. J. (2025). Phylogenomics of free-living neobodonids reveals they are a paraphyletic group from which all other metakinetoplastids are descended. bioRxiv, pages 2025–10.

Zubáčová, Z., Cimbrek, Z., and Tachezy, J. (2008). Comparative analysis of trichomonad genome sizes and karyotypes. Molecular and biochemical parasitology, 161(1):49–54.

