## Supplemental information for "Inverse relationship between mitochondrial and non-coding nuclear DNA content in free-living kinetoplastids"

### 9 Supplemental information

##### Section S1.1 Relationship between the kDNA and non-protein-coding DNA content fractions under alternative estimates of proteome lengths

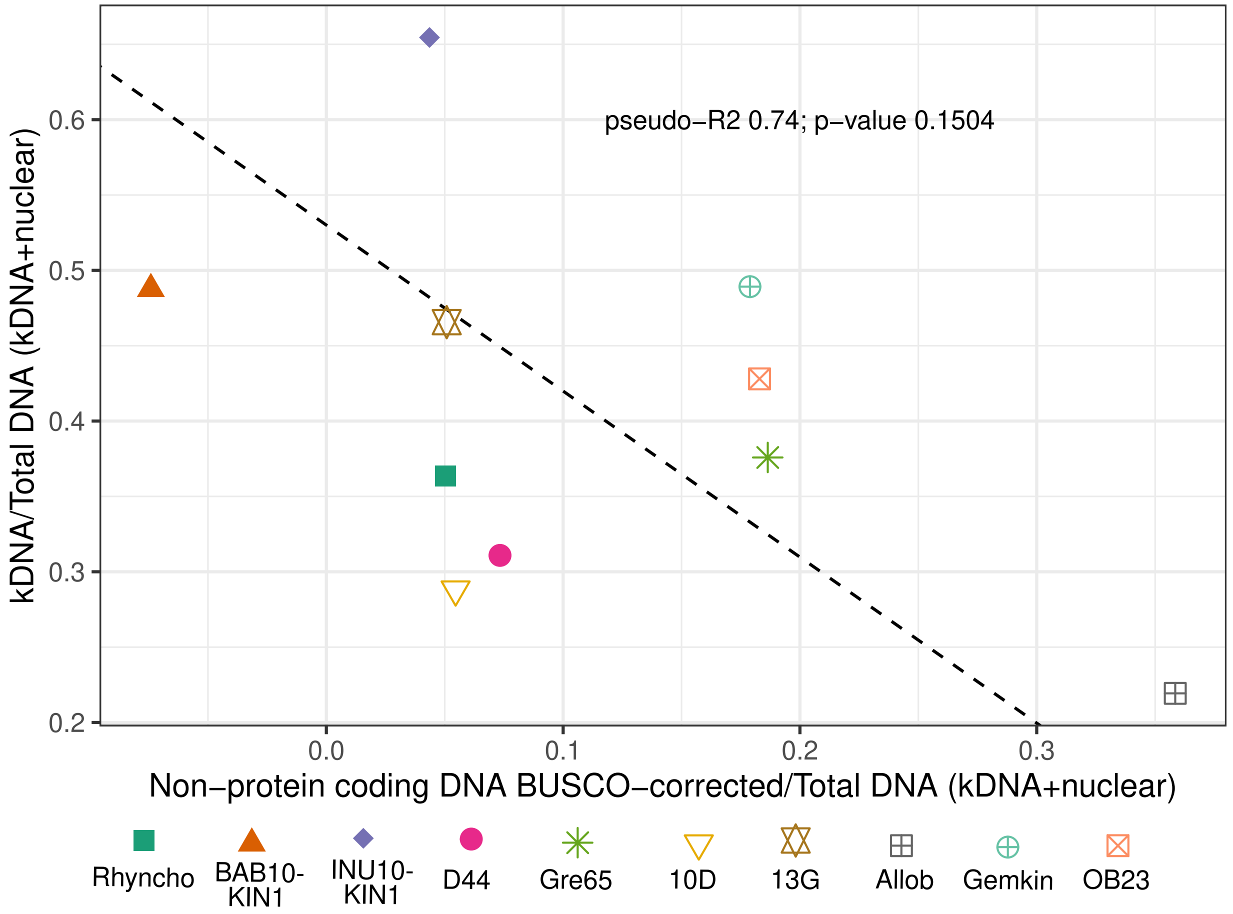


Figure S1: **Non-protein-coding nuclear and kDNA content relative to the total cellular DNA content of each isolates. The estimate of non-protein-coding nuclear and kDNA content was corrected for BUSCO completemess of non-redundant proteomes.** See Section S8.8.3 for the details on the correction for completeness; we note that such correction represents rather bold estimate, and BUSCO completeness is not the most suitable measure for assessing completeness of free-living kinetoplastid proteomes: see below.

##### Section S2.2 DNA content estimates of 10 free-living kinetoplastid isolates

Table S1: **kDNA, total nuclear DNA content, and protein-coding content of 10 free-living kinetoplastid isolates.** All DNA content values are in Mbp. For fluorescence-based estimates (columns 2-6 and 8-12), DNA content of **G1, diploid** compartments is shown. For transcriptome sequencing-based estimates, length of **haploid (G1)** protein-coding nuclear genome is shown. "Q1/Q3 kDNA/nuclear DNA" columns show 1st and 3d qunatile of predictes DNA content distributions (where predicted DNA content of single compartment of one cell represents a datapoint). "kDNA/nuclear DNA CI lower/upper" colums show the upper and confidence interval of the median of predicted DNA content.

| Isolate | Other isolate names | kDNA median | kDNA CI lower | kDNA CI upper | Q1 kDNA | Q3 kDNA | G1 kDNA count | nuclear DNA median | nuclear DNA CI lower | nuclear DNA CI upper | Q1 nuclear DNA | Q3 nuclear DNA | G1 nuclei count | protein-coding nuclear DNA HAPLOID, artefactual proteins included | protein-coding nuclear DNA HAPLOID, NO artefactual proteins | BUSCO ekaryotic | BUSCO Euglenozoa |
|---|---|---|---|---|---|---|---|---|---|---|---|---|---|---|---|---|---|
| Gemkin | *Novijibodo darinka* | 79.91 | 74.3 | 83.7 | 69.46 | 91.44 | 80 | 83.45 | 79.3 | 86.9 | 71.96 | 91.45 | 90 | 23.62 | 23.35 | 51.8 | 70.8 |
| Rhyncho | BEAP0230-"RhMon" | 52.48 | 50.2 | 54.9 | 41.07 | 64.19 | 314 | 91.87 | 88.6 | 94.4 | 80.04 | 105.58 | 311 | 35.70 | 32.50 | 49.1 | 53.9 |
| 13G | BEAP0117-"13G" | 108.45 | 105 | 114 | 98.12 | 122.20 | 107 | 124.44 | 117 | 128 | 105.64 | 141.46 | 133 | 48.04 | 46.68 | 58.4 | 65.4 |
| Gre65 | BEAP0214-"G65133" | 73.77 | 71 | 76.5 | 63.12 | 85.17 | 129 | 122.5 | 115 | 125 | 106.86 | 130.84 | 90 | 37.78 | 33.15 | 49.1 | 54.6 |
| Allob | *Allobodo yubaba*, BEAP0186-"16Ckin" | 41.34 | 37.4 | 44.1 | 31.82 | 48.92 | 75 | 147.16 | 140 | 155 | 123.86 | 170.67 | 94 | 37.39 | 36.67 | 62 | 80 |
| 10D | BEAP0071-"10D" | 55.54 | 51.2 | 59.1 | 47.87 | 62.85 | 87 | 137.25 | 133 | 141 | 125.3 | 146.79 | 114 | 54.57 | 53.04 | 59.2 | 66.9 |
| D44 | BEAP0154-"D44" | 42.28 | 40.9 | 43.7 | 36.36 | 47.67 | 180 | 93.73 | 91.4 | 95.7 | 86.99 | 101.06 | 214 | 36.53 | 36.24 | 56.4 | 71.5 |
| INU10-KIN1 | | 140.61 | 132 | 144 | 113.42 | 159.67 | 104 | 74.21 | 68.6 | 78 | 61.27 | 81.25 | 88 | 28.55 | 27.92 | 52.2 | 70.8 |
| BAB10-KIN1 | | 62.24 | 60.5 | 64.9 | 51.42 | 72.00 | 112 | 65.38 | 61.8 | 68.6 | 53.75 | 76.08 | 133 | 33.48 | 32.41 | 53.8 | 71.6 |
| OB23 | | 102.32 | 93 | 108 | 79.51 | 117.97 | 51 | 136.74 | 127 | 142 | 120.53 | 148.52 | 45 | 43.40 | 42.63 | 64.7 | 79.3 |

#### Section S3.3 Culturing conditions and information on isolates for control species with known genome size and kinetoplastids

Marine kinetoplastid isolates (Rhyncho, D44, 13G, Gre65, 10D, 13G, Allob, OB23) were cultured in "classical" RS medium with the same concentration of Nutrient component `https://mediadive.dsmz.de/medium/P5`, diluted 1:100 in of the Non-Nutrient component `https://mediadive.dsmz.de/medium/P4` (the latter mimicking seawater, 36ppt). The other three kinetoplastid isolates were also grown with Nutrient component of RS medium in 1:100 dilution as carbon source, while the Non-Nutrient component was substituted with mineral-based media of salinity and pH that corresponds to the optimal growth conditions of the isolate. For Gemkin, this was either "classical" RS medium used for other marine strains, or: CR medium, salinity 20ppt Gigeroff et al. (2023) or Non-Nutrient component of RS diluted 1:1 in distilled water. For INU10-KIN1 and BAB10-KIN1, 3N Bold's Basal Medium Bischoff (1963) and 1:100 diluted Nutrient component with distilled water as a solvent were used. For Gemkin, this was: CR medium, salinity 20ppt Gigeroff et al. (2023) or Non-Nutrient component of RS diluted 1:1 in distilled water. For INU10-KIN1 and BAB10-KIN1, 3N Bold's Basal Medium Bischoff (1963) and 1:100 diluted Nutrient component with distilled water as a solvent were used.

Additional information on kinetoplastid isolates used in this study (transcriptome sequencing data, live imaging, isolation sites, etc.) can be found in Zavadska et al. (2025).

Table S2: **Information on strains with known genome size used as a control.** Strain, species, culture medium, reference GenBank IDs and genome size used for calculations are listed. Laboratory strains derived from *Salpingoeca rosetta* strain ATCC PRA-366 are known to be both haploid and/or diploid populations Levin and King (2013); our population was predominantly haploid, according to the Sanger sequencing of polymorphic loci (see below). The genome size in the table corresponds to than of haploid assembly, 1n ("G1") cells.

| strain | species | medium | Reference assembly | Size for predictions, Mbp |
|---|---|---|---|---|
| ATCC50162 | *Diplonema papillatum* | ATCC medium 1532 | GCA_030068145.1 | 280.283133 |
| ATCC30864 | *Capsaspora owczarzakii* | ATCC medium 1034 | GCA_033442345.1 | 27.398043 |
| ATCC PRA-366 | *Salpingoeca rosetta* | Richter-Sigona (RS) medium, 1:100 NM:NNM | GCA_033442325.1 | 55.448948 |
| BY4741 (MATa his31 leu20 met150 ura30 haploid strain) | *Saccharomyces cerevisiae* | 5% Yeast Extract, 3% v/v Glycerol, 1% K2HPO4, 5% v/v EtOH medium | GCA_000766575.2 | 11.529746 |

**Section S4.4 Stain concentrations**

$$MW(PI) = 668.4g/mol \tag{3}$$

$$MW(DAPI) = 350.3g/mol \tag{4}$$

The concentrations of dyes in the staining solutions:

$$PI = 125\mu g/ml \tag{5}$$

$$DAPI = 60\mu g/ml \tag{6}$$

Assuming at least 10 µl of liquid remains on 24mmx24mm coverslip (usually it is around 12-15 µl), the number of PI molecules will be:

$$PI = 10\mu l \times 125\mu g/ml = 1.25\mu g(PI)/coverslip \tag{7}$$

$$\frac{1.25\mu g(PI)}{668.4\mu g/\mu mol} = 1.87 \times 10^{-9} mol(of PI) \tag{8}$$

$$1.87 \times 6.02 \times 10^{14} PI molecules/coverslip \tag{9}$$

The number of DAPI molecules will be:

$$DAPI = 10\mu l \times 60\mu g/ml = 0.6\mu g(DAPI)/coverslip \tag{10}$$

$$\frac{0.6\mu g(DAPI)}{350.3\mu g/\mu mol} = 1.71 \times 10^{-9} mol(of DAPI) \tag{11}$$

$$1.71 \times 6.02 \times 10^{14} DAPI molecules/coverslip \tag{12}$$

Assume the nuclei of *Diplonema papillatum*, the species with the largest genome size used in this study, are squares of 5µmx5µm, they cover all the surface of the 24mmx24mm coverslip, and all of them are in the G2 phase of the cell cycle, containing 560Mbp of DNA each. The hypothetical maximum number of *Diplonema papillatum* nuclei which fits on the coverslip would be:

$$(576 \times 10^3 \mu m^2)/25\mu m^2 = 23040 nuclei/coverslip \tag{13}$$

All nuclei that could be allocated on one coverslip (as a monolayer), would contain

$$560Mbp \times 23040 = 12902400Mbp = 12.9 \times 10^{12} bp \tag{14}$$

PI:DNA binding stoichiometry 1 PI molecule : 4-5 basepairs (as mono-intercalating dye; we take 4 as minimum) Lin et al. (2003)
DAPI:DNA binding stoichiometry 1 DAPI molecule : 4 basepairs Larsen et al. (1989)

So, given the number of DNA basepairs which could be positioned on one coverslip (organised as a monolayer of square nuclei), the number of dye molecules in the staining solution in which the coverslip is immersed, and the stoichiometry of PI and DAPI binding with DNA, the number of molecules required to bind all the DNA on the coverslip, will be, for either DAPI or PI:

$$(12.9 \times 10^{12} bp)/4 = 3.225 \times 10^{12} dye\ molecules \tag{15}$$

The number of both DAPI ($1.03 \times 10^{15}$) and PI ($1.13 \times 10^{15}$) molecules in staining solution exceeds the minimum required number of molecules, $3.225 \times 10^{12}$, by a magnitude of more than two orders, even given the most unrealistic, worst-case estimates of the amount of DNA and staining solution present on the slide.

#### Section S5.5 Image acquisition setup - interval between slices in z-stack

##### Section S5.5.1 Resolution in Z-plane

The axial resolution, or depth of field, of wide-field fluorescent microscopy can be estimated using the equation 16, where $n$ is the refractive index of the sample, $\lambda$ is the fluorescence wavelength, and $NA$ is the numerical aperture of the objective.

$$d_{ax.} \approx \frac{2n\lambda}{NA^2} \tag{16}$$

The estimated depth of focus for the emission maxima of DAPI ($\lambda_{DAPI} \approx 465\text{nm}$) and PI ($\lambda_{PI} \approx 620\text{nm}$) was 640 nm and 840 nm (17, 18).

$$d_{DAPI} = \frac{2 \times 1.33 \times 465nm}{1.4^2} \approx 630nm \tag{17}$$

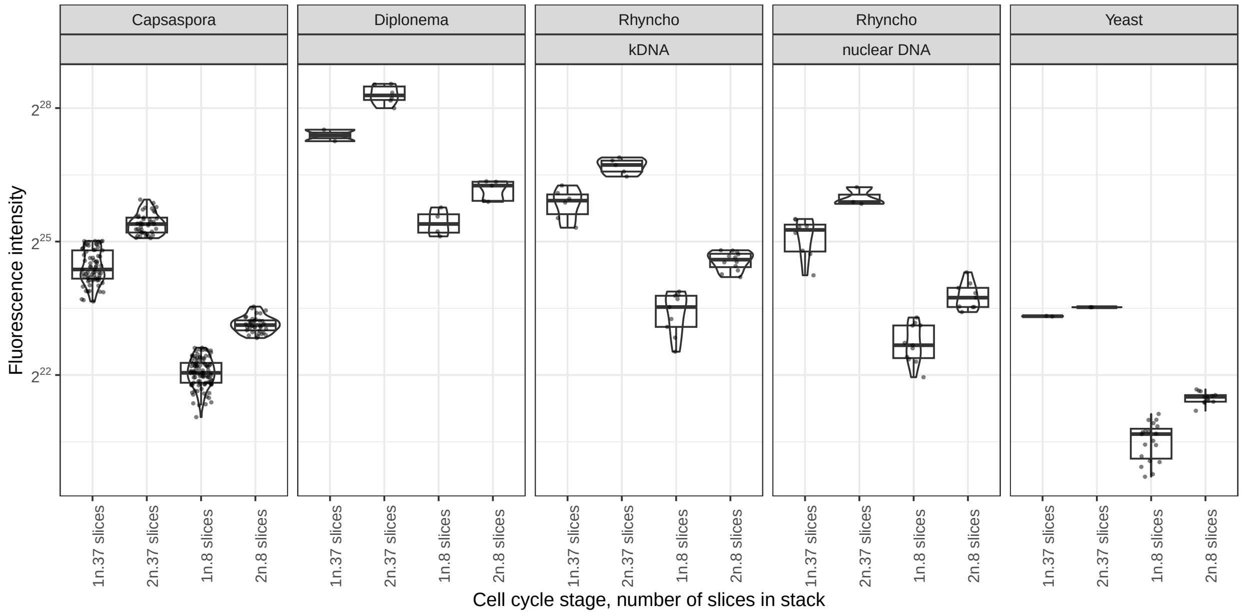


Figure S2: **Distributions of fluorescence intensities depending on the number of slices and interval of the stack**. Each facet represents values of fluorescence intensities (y-axis) obtained for a given compartment of a given species; each dot represents a measurement from one cell. On x-axis, the values are split depending on the assigned cell cycle stage ("G1" or "G2") and the stack setting ("8 slices" or "37 slices"). Note the log2 scale.

#### Section S6.6 Statistical analysis benchmark

##### Section S6.6.1 Subsets used in data analysis

The overall experimental design assumed obtaining measurements from several independent sets of cell populations (eukaryotic cell "passages") of control plus test species. These sets were passed, grown, put on the same slide, fixed, stained and imaged together. These independent sets are biological replicates of the study. Within each biological replicate, technical replicates have been present in the majority of cases: two different slides with the same populations of cells were prepared.

As a result, the raw data used in the analysis, represented by the fluorescence intensities of nuclei or kDNA of individual cells, had the underlying structure of a technical replicate (= glass slide) < biological replicate (same populations of cells and same day/batch of cell fixation and staining) Figure S3.

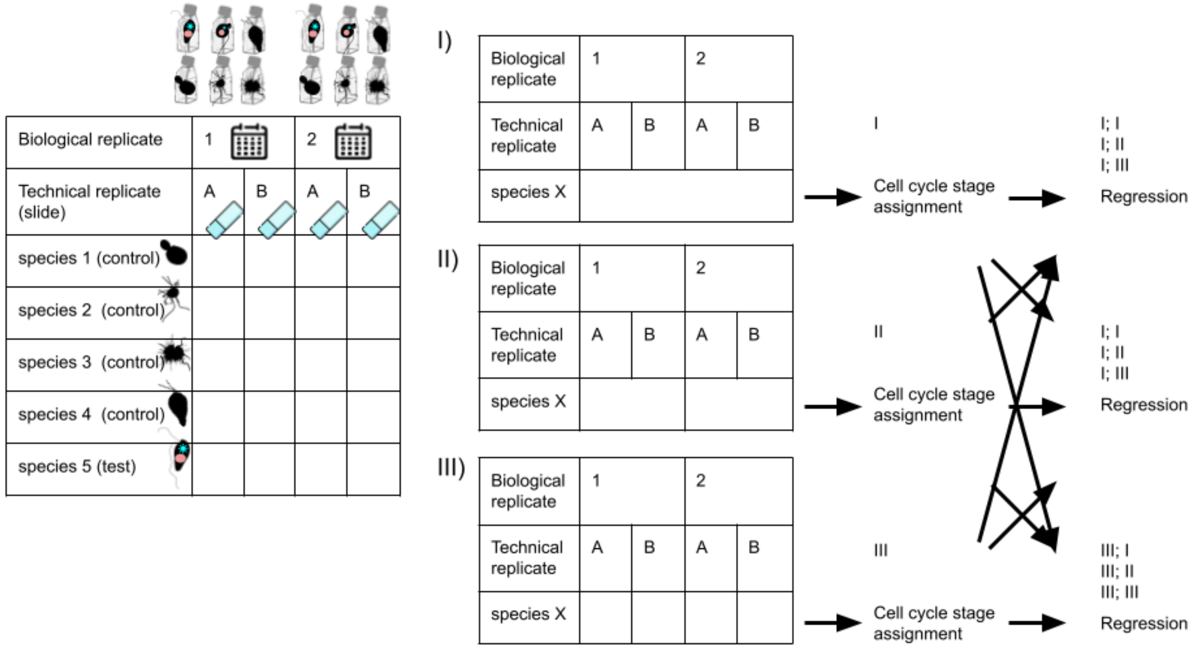


Figure S3: **Schematic illustration of data structure and nature of replicates.** Further description in the text.

Procedures of outlier filtering combined with the cell cycle stage assignment, and regression could be performed separately on different data subsets: separate sets of cells from every technical replicate (III in Figure S3), separate sets of cells from every biological replicate (II in Figure S3), or all cells with no subsetting (I in Figure S3). Furthermore, these sets could be combined: while outlier filtering with the cell cycle stage assignment is performed on a given subset (for example, all cells: "I"), the regression can be performed on a different one (for example, separately on values within each biological replicate: "II"). Using different subsets for different procedures can have advantages, given the nature of the data and the putative error sources. For example, combining cells of a given species from all biological replicates to predict the cell cycle stage can give more accurate result, as cell populations from different experiments might have the majority of cells shifted to only one stage in the cell cycle, G1 or G2.

All possible ways of subsetting data (12 combinations) were tested for the accuracy of genome size predictions the same way as described in the main text: DNA content of one out of three control species was predicted based on the regression infered using the rest three control species. The accuracy of predictions obtained for each subset combination is summarised in Figure S4 and Figure S5.

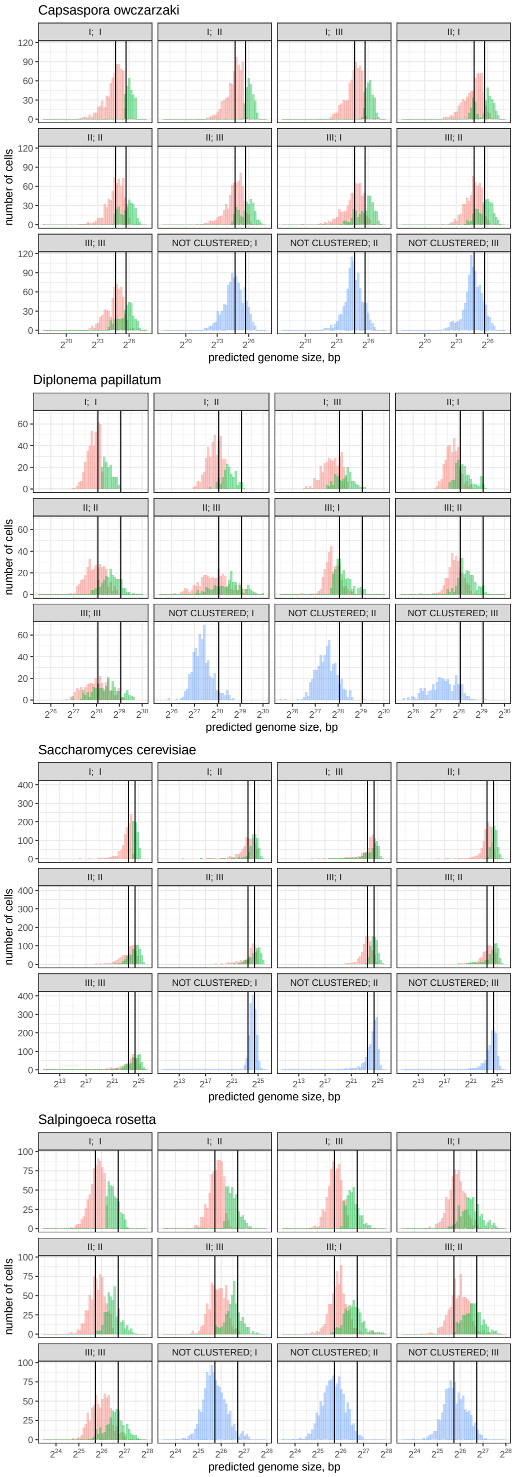


Figure S4: **Distribution of DNA content predicted for the control species based on fluorescence intensities, compared to the expected genome sizes.** FThe y-axis represents the number of G1 (red) and G2 (green) cells with a predicted DNA content (x-axis). The vertical solid lines indicate the actual sizes of control species genomes - for the G1 and G2 cells; the distributions tend to center around the vertical line, indicating that the predicted genome sizes, on average, correspond to the real ones for control species. Facets represent data subsets used for outlier filtering combined with the cell cycle stage assignment (first number in the facet header), and for regression inference (second number in the facet header); to match the number with the type of subset, refer to the Figure S3 and the description in text.

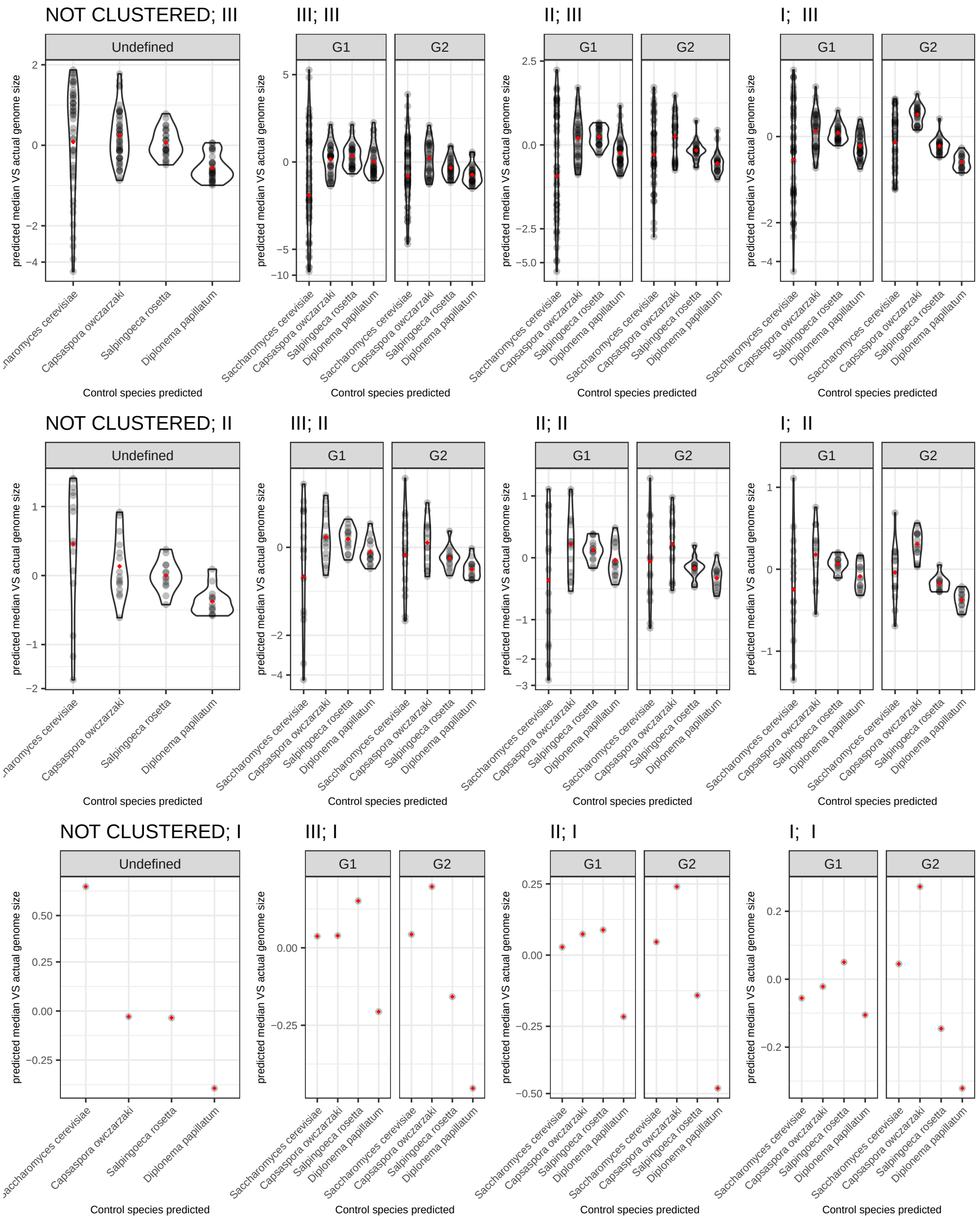


Figure S5: **Difference of medians of genome sizes predicted for the control species from the expected genome sizes, as a fraction of the expected genome size.** Each dot represents a median obtained from the predicted values of cells from either a single technical (top panel), biological (middle panel) replicates, or all the measurements taken together (bottom panel), from which the actual genome size was subtracted, and the difference is normalised by the expected genome size: $(Median_{predicted} - bp_{actual})/bp_{actual}$. Red diamonds represent the means of medians from all biological/technical replicates, from which the actual genome size was subtracted, and the difference is normalised by the expected genome size. Facets represent subsets used for outlier filtering combined with the cell cycle stage assignment (first number in the facet header), and for regression inference (second number in the facet header); to match the number with the type of subset, refer to the Figure S3. Note different scales on the y-axis. The inverse hyperbolic sin transformation was used for y-axis values in this case, to efficiently represent both scattered values for *S. cerevisiae* and less scattered values from all other species.

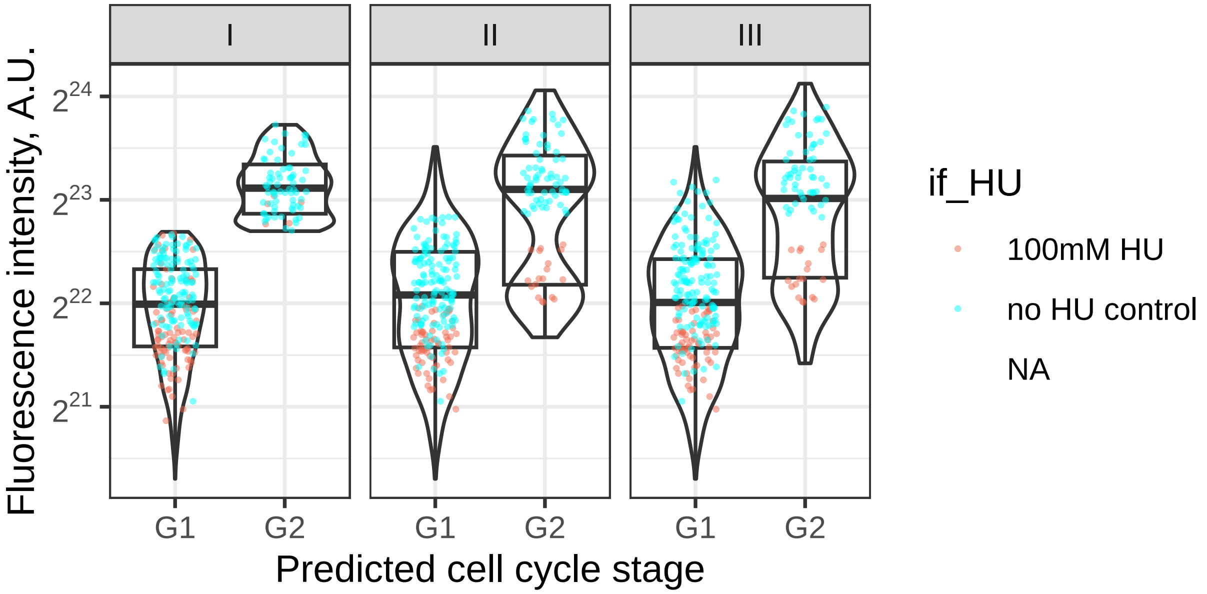


Figure S6: **Distributions of raw fluorescence intensities of *C. owczarzaki* nuclei, across cell cycle stages assigned, for different subsets used for classification.** Violin and boxplots represent the overall distribution of intensities from **all cells** measured across **all biological replicates**. The dots in red and represent the intensities measured for Hydroxyurea-treated cells and the dots in blue - intensities from control (non-treated) cells from a given biological replicate. The majority of red dots should be G1 cells, and are expected to be classified as such, while blue dots are expected to be both G1 and G2 cells. Facet headers show the subset of cells used in outlier filtering and cell cycle stage assignment procedure; to match the number with the type of subset, refer to the Figure S3. Note that the total number of cells might differ between subsets, as some values can be discarded as outliers in one subset, but not in the other.

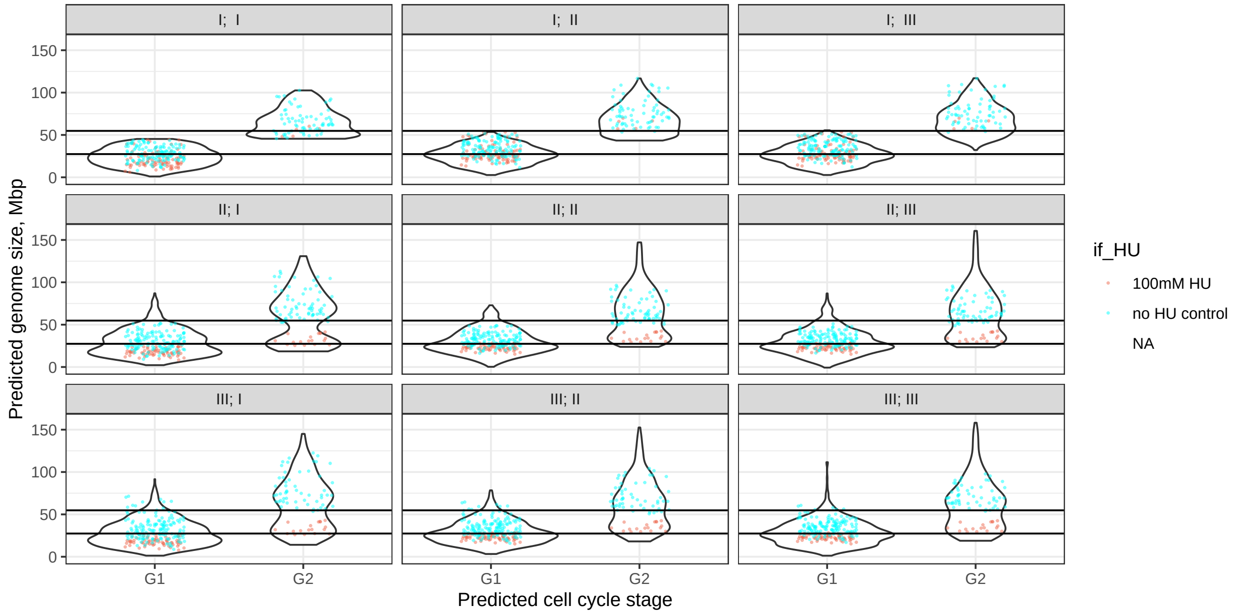


Figure S7: **Distributions of predicted DNA content of *C. owczarzaki* nuclei, across cell cycle stages assigned, for different combinations of subsets.** Violin and boxplots represent the overall distribution of predicted genome sizes from **all cells** measured across **all biological replicates**. The dots in red and represent the intensities measured for Hydroxyurea-treated cells and the dots in blue - intensities from control (non-treated) cells from a given biological replicate. The majority of red dots should be G1 cells, are expected to be classified as such and to have predicted DNA content closer to that expected for G1 cells, while blue dots are expected to be both G1 and G2 cells. Facet headers show the subset of cells used in outlier filtering and cell cycle stage assignment procedure (first number) and for regression inference (second number); to match the number with the type of subset, refer to the Figure S3. Note that the total number of cells might differ between subsets, as some values can be discarded as outliers in one subset, but not in the other.

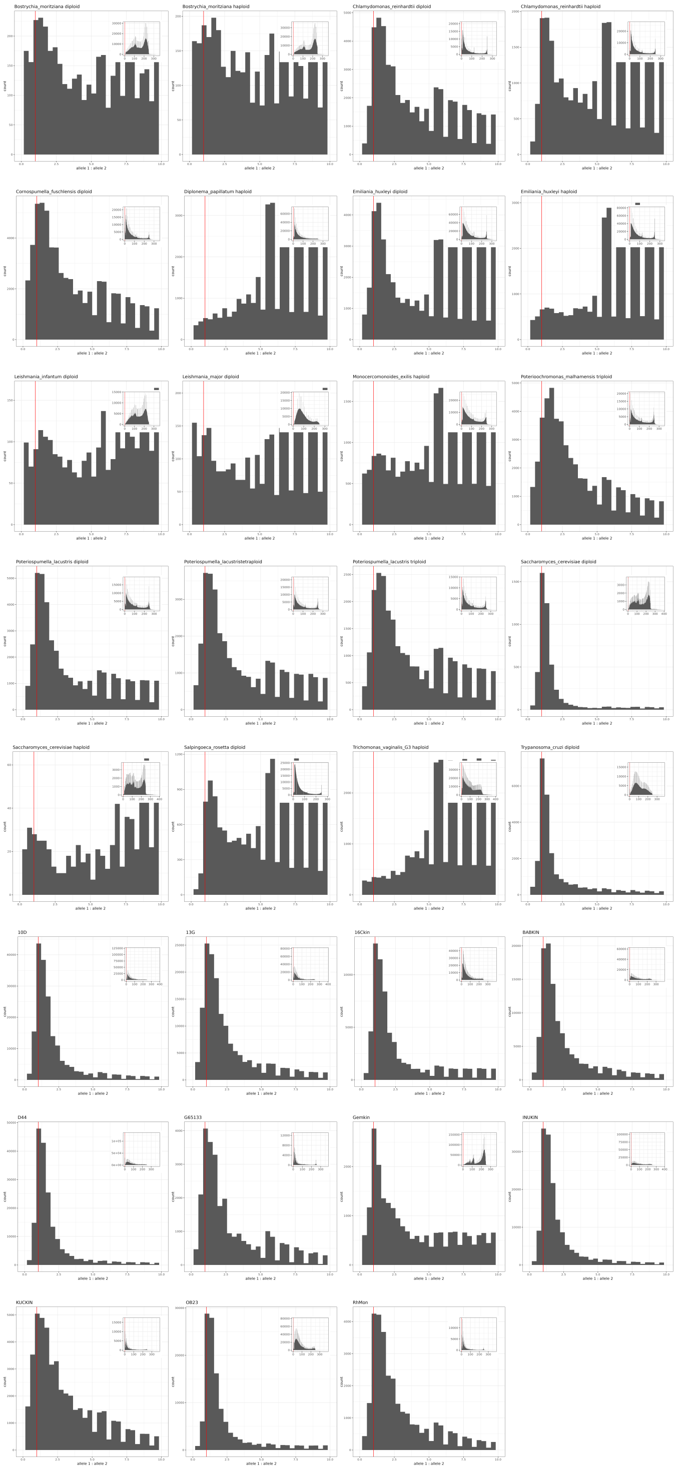


Figure S8: **Distributions of major and minor allele frequency ratios in SNPs detected by transcriptomic read mapping.** Each facet represents the distribution for each species; for the control species with known ploidy, the ploidy is also indicated. The red line corresponds to the ratio of 1: where the peak of distribution is expected for the diploid organisms. The distribution of the whole range of ratios is shown at the right upper corner of each facet, while large plots indicate the distribution of ratios ranging from 0 to 10. Frequency ratios higher than 10 mean that SNPs are less likely to result from the presence of multiple alleles (they could be artefacts of data analysis, sequencing, mutations in the process of transcription, etc).

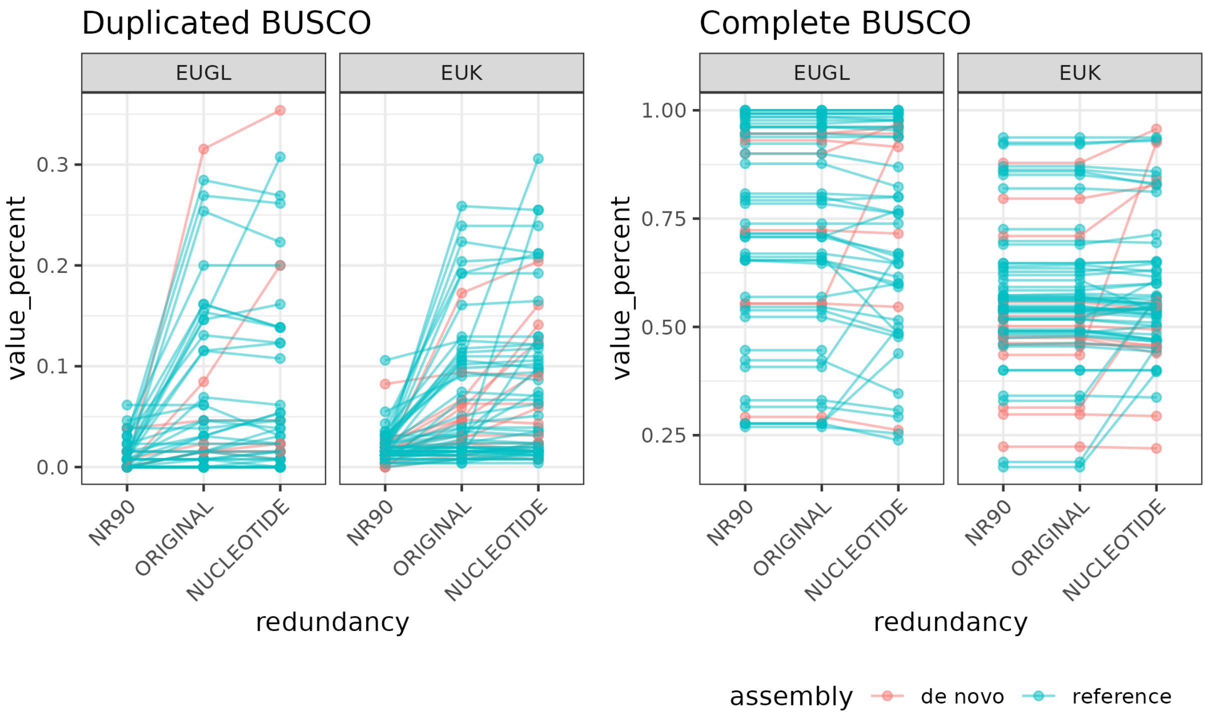


Figure S9: **Effect of redundancy reduction and data type on the number of BUSCO duplicates and Complete BUSCO genes.** Each dot represents a proteome/assembly from one species under one setting, and its position on the y-axis illustrates the percent of BUSCO genes ("Duplicated" or "Complete"). Dots from the same species are linked by lines. Colour represents whether the assembly/proteome is a part of *de novo* assembled/predicted set, used for a pipeline test in Section S8.8.3, or "reference" set. Each subplot represents BUSCO database used: "EUK" - Eukaryota BUSCO dataset; "EUGL" - Euglenozoa BUSCO dataset. The x-axis represent the combination of redundancy reduction setting and data type assessed: "ORIGINAL" - proteome with no redundancy reduction applied; "NR90" - proteome after discarding duplicates at CD-hit at 90% aa identity threshold; "NUCLEOTIDE" - nucleotide transcripts (for a few species, only CDSs were available).

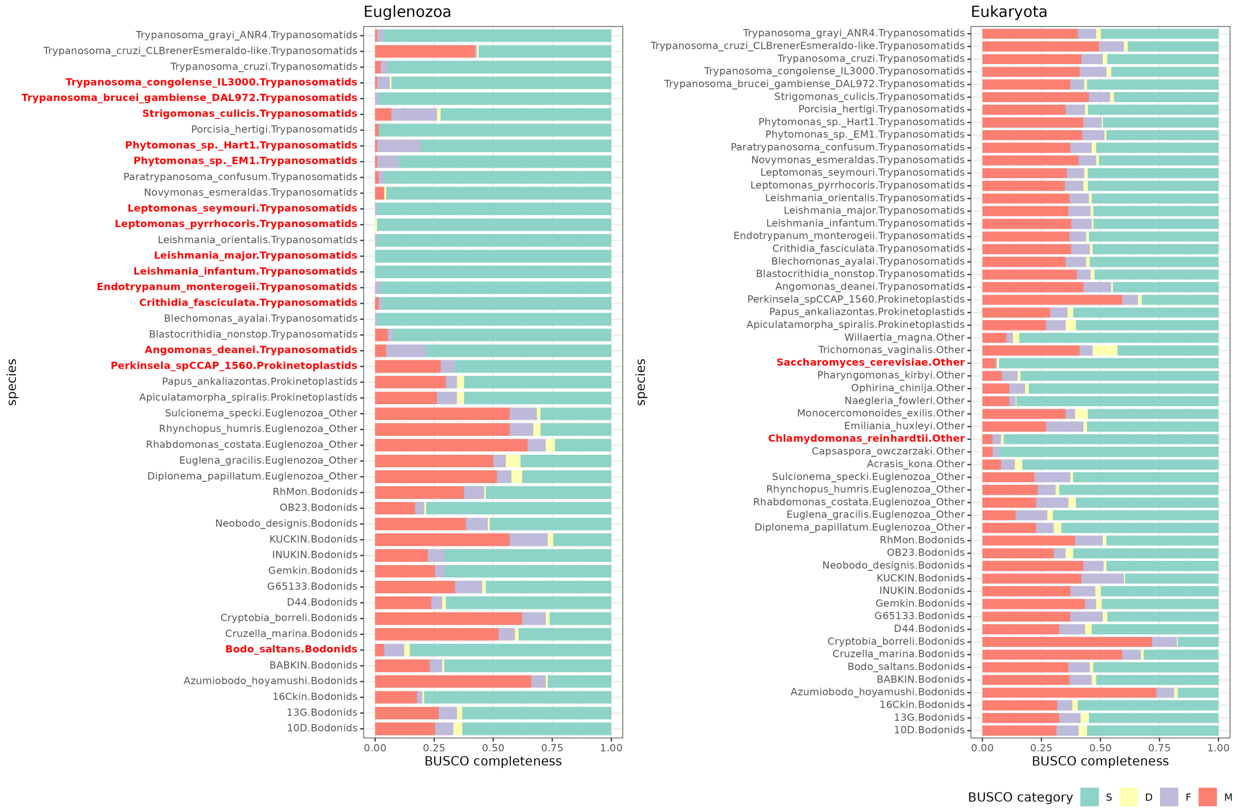


Figure S10: **BUSCO completeness of proteomes from diverse protists.** Facets represent completeness according to "Eukaryota" and "Euglenozoa" BUSCO datasets. The x-axis represents the fraction of each BUSCO gene category. Colours of the bars indicate fractions of: "S" - Complete and single-copy; "D" - Complete and duplicated; "F" - Fragmented; "M" - Missing BUSCO genes. On the y-axis, species and their taxonomic affiliation are indicated. Species which are included in "Eukaryota" or "Euglenozoa" BUSCO databases are highlighted in red.

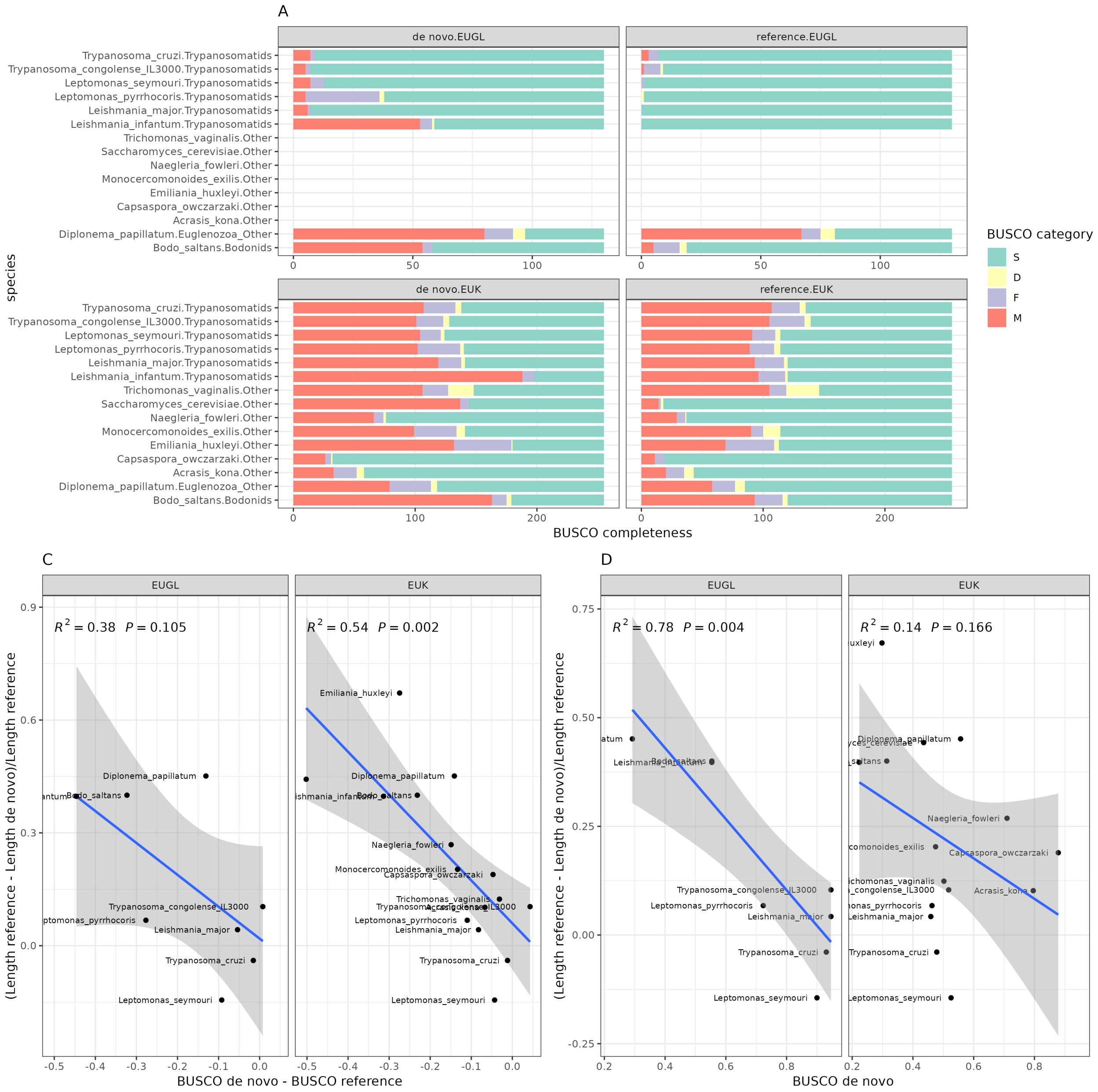


Figure S11: **A - BUSCO completeness of *de novo* predicted proteomes and reference proteomes of the same species.** "EUK" - Eukaryota BUSCO dataset; "EUGL" - Euglenozoa BUSCO dataset. Colours of the bars indicate fractions of: "S" - Complete and single-copy; "D" - Complete and duplicated; "F" - Fragmented; "M" - Missing BUSCOs. **C - Difference in length between *de novo* assembled proteome and the reference proteome, expressed as a fraction of reference proteome length (x-axis) vs difference in BUSCO completeness between *de novo* assembled proteome and the reference proteome (y-axis). D - Difference in length between *de novo* assembled proteome and the reference proteome, expressed as a fraction of reference proteome length (x-axis) vs difference in BUSCO completeness of the *de novo* assembled proteome (y-axis).** Linear regression fits on C and D were performed using the "geom_smooth" function from the ggplot2 R package, using "lm" method from stats R package for the fit.

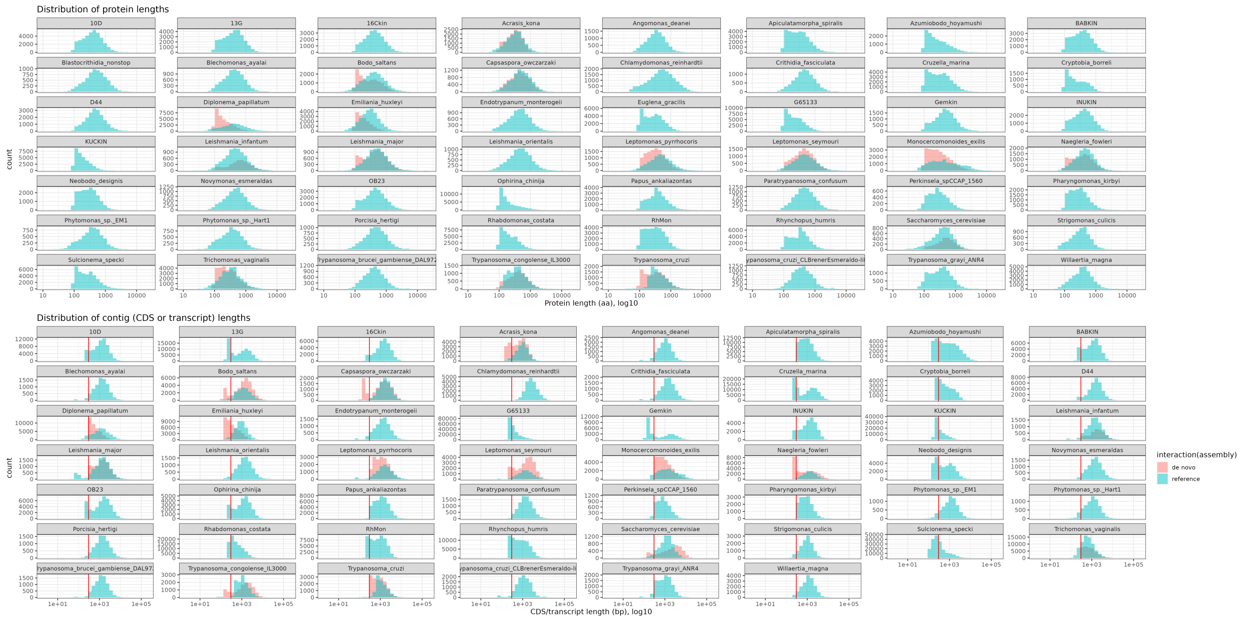


Figure S12: **Length distribution of predicted proteins and contigs.** Each subplot represents one species from the dataset; the x-axis represents lengths of proteins or contigs (note the log-10 scale). For protein length distribution, redundancy-reduced proteomes were used. The red vertical line on the contig lengths plot indicates the lengths of 300bp: proteins below 100aa are discarded at the protein annotation step; therefore, proteins from contigs shorter than 300bp are not included in the predicted proteome.

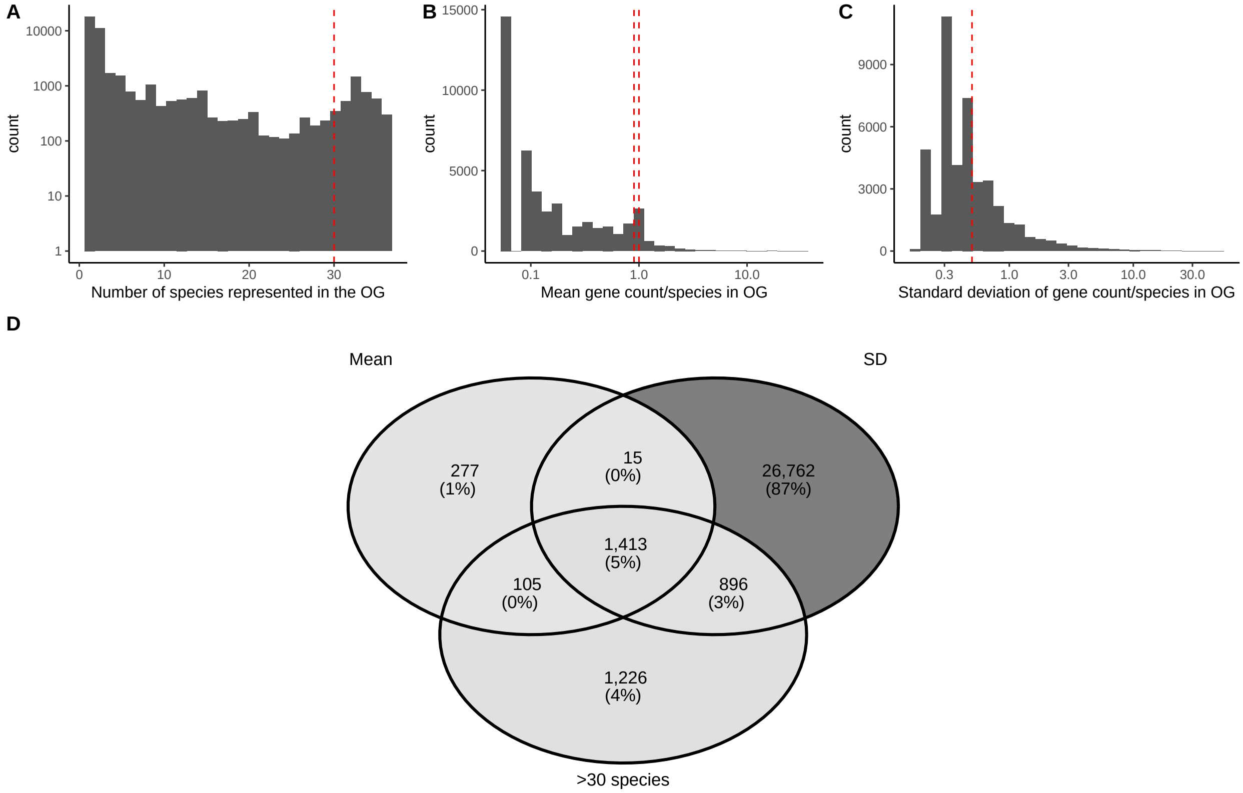


Figure S13: **Subsetting OGs. A-C**: distributions of parameters used to establish thresholds to subset OGs. The count of OGs with a given parameter value is represented on the vertical axis, and the values of the three different parameters are represented on the horizontal axis. The dashed red vertical lines in A-C represent thresholds used for subsetting OGs. Note log10 scales in A-C. **D**: the count of OGs with the mean number of proteins per single species between or equal to 1 and 0.9 ("Mean"), standard deviation of number of proteins per single species lower than or equal to 0.5 ("SD"), and the number of species with at least one protein in a given OG higher than 30 (">30 species").

#### Section S8.8.5 Short proteins in proteomes

High numbers of short proteins could affect the final estimate of total proteome length. Short proteins could either be artefacts of protein prediction software, which do not represent real sequences translated in the organism, or fragments of longer protein-coding genes, which do represent parts of "real" sequences. In the latter case, the sum lengths of

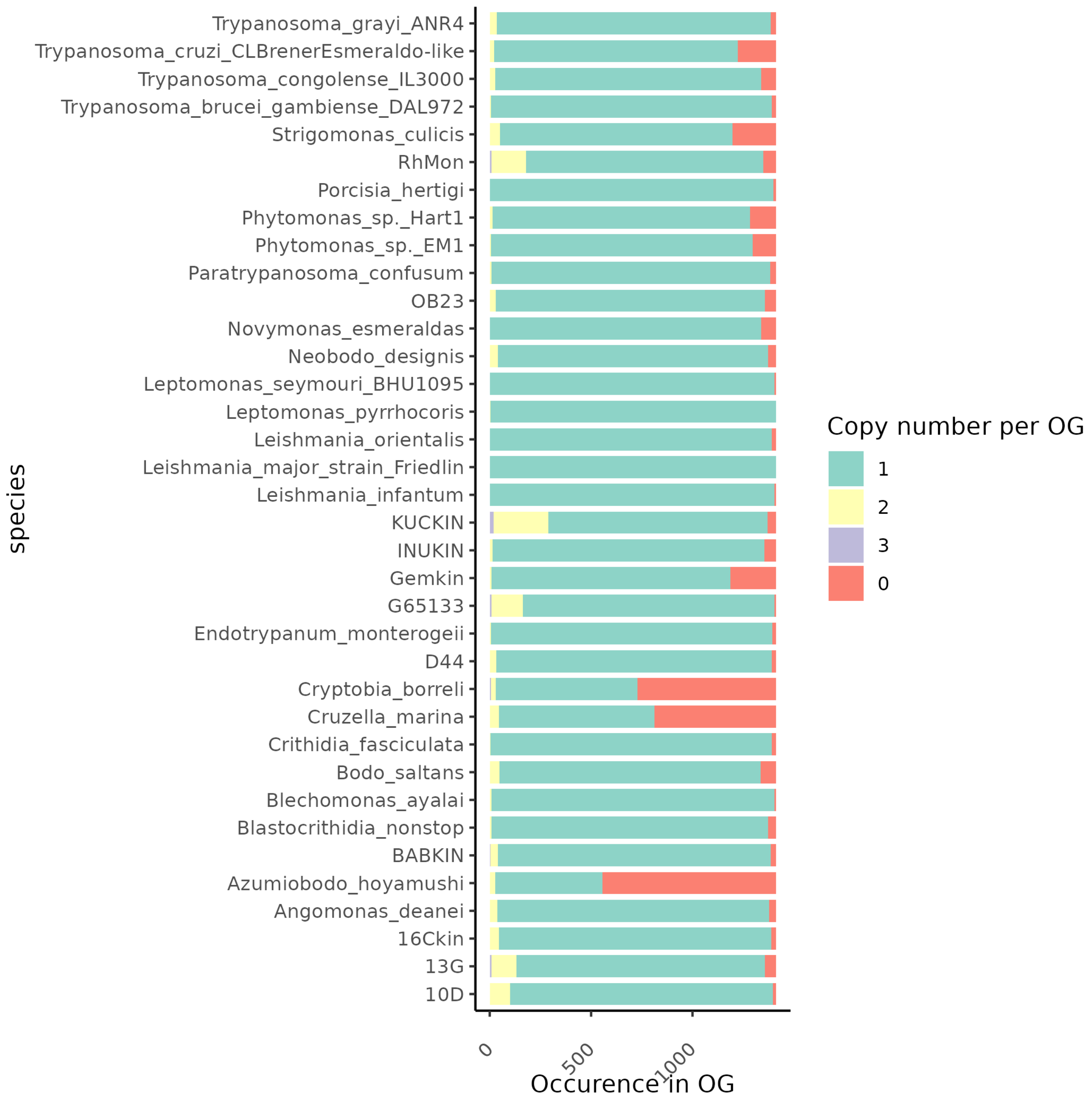


Figure S14: **Representation of genes from M3USCO OG set across Metakinetoplaastid proteomes.**

shorter proteins should equal the lengths of a single longer protein, and protein fragmentation does not affect the total length of the proteome. The overrepresentation of short sequences which do not correspond to "real" proteins could lead to an overestimation of total proteome lengths. "Real" short protein sequences can be distinguished from artefactual ones by the presence of homology to the proteins of other related species (though some genes and proteins can appear spontaneously from the non-coding ancestral sequence, the majority of protein-coding genes still evolve from ancestral protein-coding sequences). In such case, the "real" proteins are likely to be included into OG with sequences from more than one species, when classified by OrthoFinder. The artefactual proteins, in contrast, would not have homologs in the related species, and would be assigned to "singletons", or "species-specific OGs" - OGs consisting entirely of genes from one species. Alternatively, artefactual proteins would end up not being assigned to any OG.

In the Metakinetoplastid OG dataset, described above, we analysed the total lengths of proteins shorter than 200aa, which belong to either singleton OGs or do not belong to any OG Figure S15. Among the proteomes used in this study, three species: G65133, RhMon and KUCKIN have a total length of short (<200 aa) proteins contributing more than 10% towards the total length of all the proteins Figure S15. As mentioned above, KUCKIN initially had lower read coverage than other species from the study, and its proteome length is not considered in the final estimations. For G65133 and RhMon, at least half of the total length of short proteins can be attributed to proteins in singletons, or not assigned to any OG: the "artefactual" proteins, without a homology to any sequences in related species Figure S15. In G65133, only 15% of the total non-redundant proteome lengths could be attributed towards the sequences which are most likely artefacts of data analysis and do not represent the underlying biological entities. For other species used in this study, the fraction of such artefacts is negligible, in terms of their lengths relative to the whole proteome.

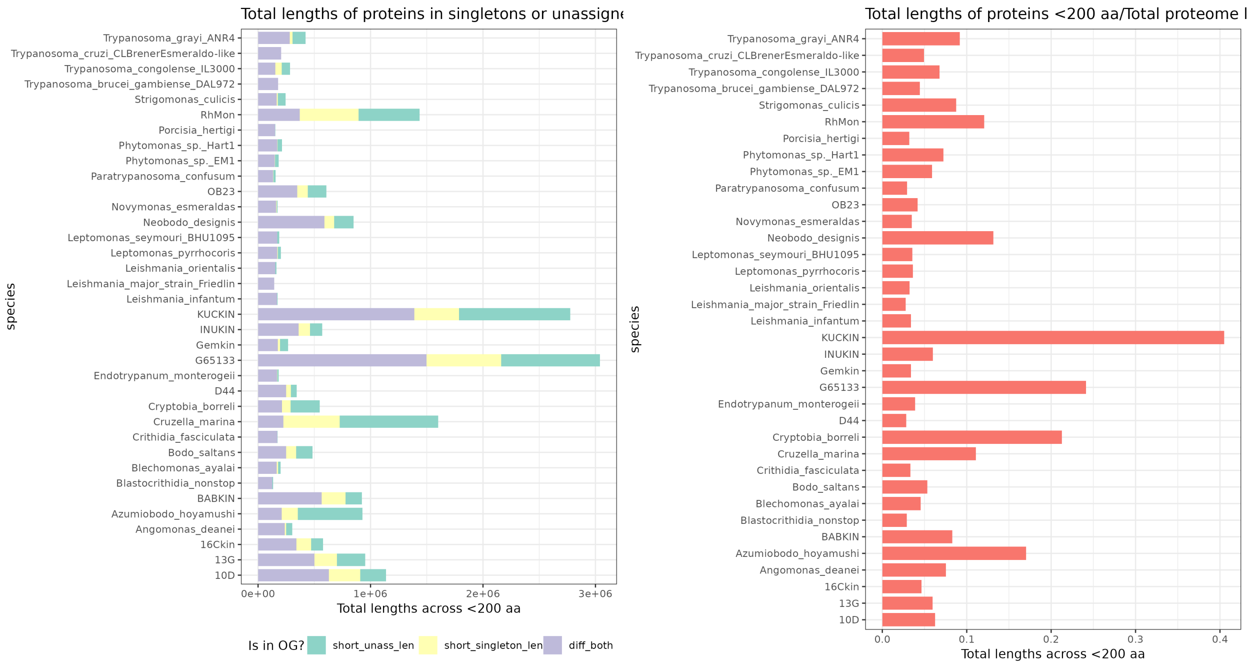


Figure S15: **The total aa length of short (<200 aa) proteins, their OG affiliation, and the fraction of total proteome length constituted by the short proteins.** The colours on the first subplot represent: "short_unass_len" - total length of short proteins which are not assigned to any OG; "short_singleton_len" - total length of short proteins which are assigned to sigletons; "diff_both" - total length of short proteins which are assigned to an OG with proteins from more than a single species.
